# Intrinsic-Dimension analysis for guiding dimensionality reduction and data-fusion in multi-omics data processing

**DOI:** 10.1101/2024.01.23.576822

**Authors:** Jessica Gliozzo, Valentina Guarino, Arturo Bonometti, Alberto Cabri, Emanuele Cavalleri, Mauricio Soto-Gomez, Justin Reese, Peter N Robinson, Marco Mesiti, Giorgio Valentini, Elena Casiraghi

## Abstract

The advent of high-throughput sequencing technologies has revolutionized the field of multi-omics patient data analysis. While these techniques offer a wealth of information, they often generate datasets with dimensions far surpassing the number of available cases. This discrepancy in size gives rise to the challenging “small-sample-size” problem, significantly compromising the reliability of any subsequent estimate, whether supervised or unsupervised.

This calls for effective dimensionality reduction techniques to transform high-dimensional datasets into lower-dimensional spaces, making the data manageable and facilitating subsequent analyses. Unfortunately, the definition of a proper dimensionality reduction pipeline is not an easy task; besides the problem of identifying the best dimensionality reduction method, the definition of the dimension of the lower-dimensional space into which each dataset should be transformed is a crucial issue that influences all the subsequent analyses and should therefore be carefully considered.

Further, the availability of multi-modal data calls for proper data-fusion techniques to produce an integrated patient-view into which redundant information is removed while salient and complementary information across views is leveraged to improve the performance and reliability of both unsupervised and supervised learning techniques.

This paper proposes leveraging the intrinsic dimensionality of each view in a multi-modal dataset to define the dimensionality of the lower-dimensional space where the view is transformed by dimensionality reduction algorithms. Further, it presents a thorough experimental study that compares the traditional application of a unique-step of dimensionality reduction with a two-step approach, involving a prior feature selection followed by feature extraction.

Through this comparative evaluation, we scrutinize the performance of widely used dimensionality reduction algorithms. Importantly, we also investigate their impact on unsupervised data-fusion techniques, which are pivotal in biomedical research. Our findings shed light on the most effective strategies for handling high-dimensional multi-omics patient data, offering valuable insights for future studies in this domain.

**Graphical Abstract:** 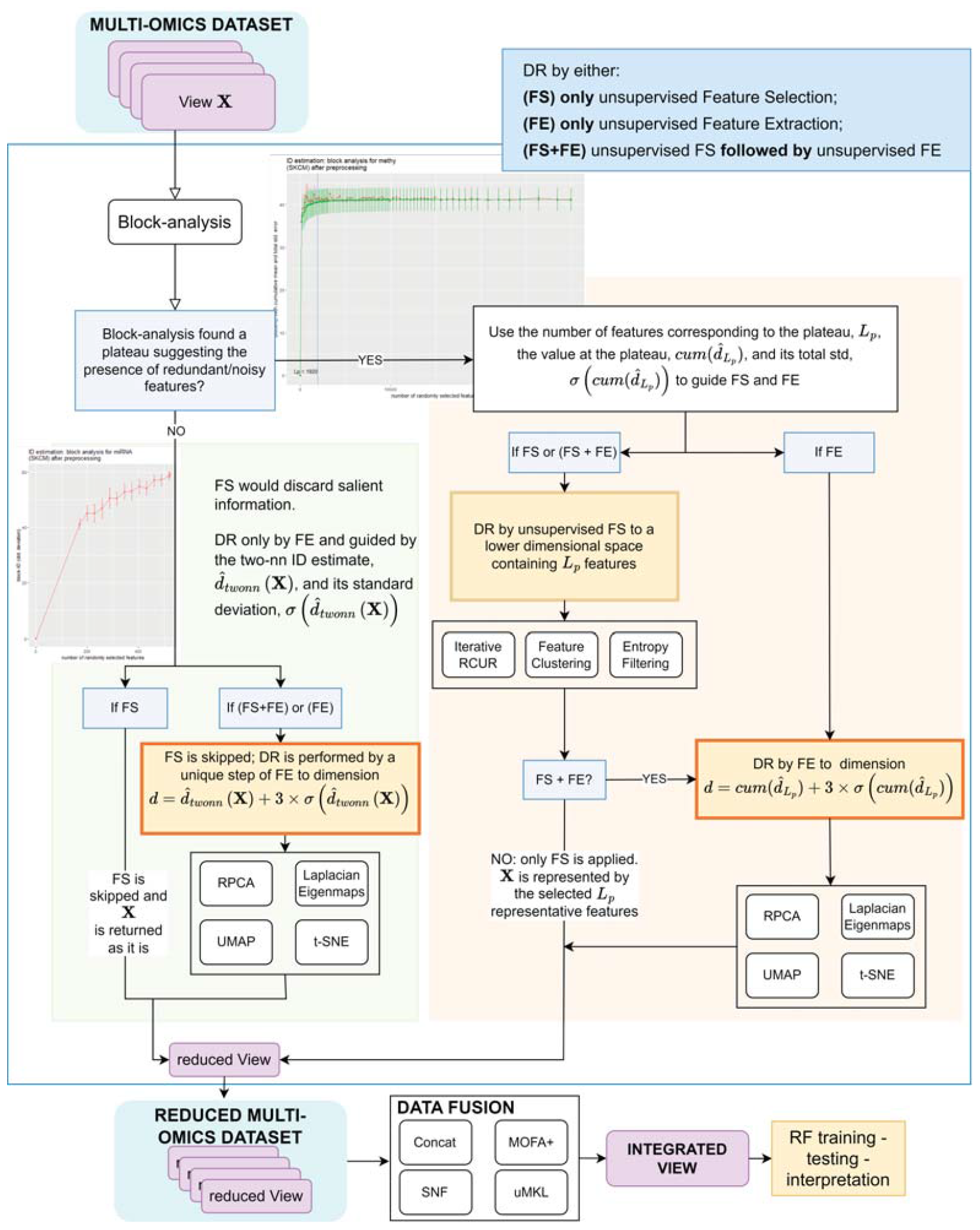

**Highlights:**

- We introduce a flexible pipeline to guide in a principled way feature selection and feature extraction methods to reduce the high dimensions and to contrast the curse of dimensionality that affects multi-omics data.
- We harness the power of cutting-edge Intrinsic Dimensionality (`id`) estimation through block-analysis, providing an unbiased estimation of the individual `id`s for each view within a multi-modal dataset.
- We use an exhaustive set of diverse multi-omics cancer datasets from the well-known TCGA dataset to show that the automatic analysis of the distribution of the block-`id`s characterizing each omics-view leverages dimensionality reduction, by (1) evidencing feature noise and redundancy, and (2) providing an unbiased estimate of the `id` for each view, to be used for setting the dimension of the reduced space. This avoids empirical or heuristic choices and allows tailoring the reduction to each data-view.
- The crucial information gained by block-analysis allowed proposing a two-step dimensionality-reduction approach combining feature selection and feature extraction. Our comparative evaluation shows the effectiveness of the proposed technique and its synergy with state-of-the-art data-fusion techniques applied in a multi-omics context.
- We show that the proposed reduction pipeline leverages traditional dimensionality reduction and state-of-the-art data-fusion algorithms. Indeed, it obtains effective performance when predicting overall survival events with simple random forest classifiers, often preferred in the biomedical field due to their robustness, efficiency, and interpretable nature.

## 1. Introduction

In the biomedical research field, the emergence of high-throughput technologies has revolutionized the acquisition of vast and diverse omics data types such as genomic, transcriptomic, proteomic, and methylomic data [1, 2]. These distinct modalities (views) provide valuable insights into the intricate molecular landscape governing biological processes and diseases; if appropriately processed and integrated they can uncover crucial disease triggers and enhance our understanding of various health conditions [3, 4, 5, 6].

However, the analysis of multi-omics data presents significant challenges due to their high dimensional nature and multi-modality. In particular, the high-dimensional nature of omics data results in high computational costs, data sparsity, and over-fitting due to the presence of noisy, uninformative, and redundant features. These problems collectively are referred to as the “curse of dimensionality”, and can bias practically all results obtained from these data. This is particularly true in bio-medical datasets, often characterized by high-dimension and small-sample-size, that is by a large number of features relative to the number of samples. Such datasets may easily reach a level of sparsity that causes samples to appear distributed on the boundaries of the hyperspace, affecting the reliability of subsequent supervised or unsupervised analyses [7, 8, 9].

To address these issues, unsupervised Dimensionality Reduction (DR) gained a lot of interest over the past decade and is now recognized as being a crucial preliminary phase in various fields [10]. DR techniques, including feature selection and feature extraction methods, mitigate the curse of dimensionality by reducing the dimension of the input dataset so that it concisely conveys similar information. In case of bio-medical multi-modal datasets, DR may be individually applied to reduce each input modality (view) and better expose its characterizing informative content. This would aid the following data-fusion task, for which several promising algorithms have been already presented in the bio-medical literature [11].

However, while feature selection and feature extraction methods have shown their own advantages and several reviews describe and eventually compare their successful results [12], to the best of our knowledge no paper investigated the following two crucial choices that should be carefully considered when analyzing and reducing high-dimensional datasets, potentially affected by the curse of dimensionality. First, there is no rule of thumb that allows claiming that a dataset is affected by the curse of dimensionality/small-sample-size problem. Second, the choice of the dimension of the reduced space is one of the most crucial choices; too low values would cause the loss of information, while too large values would not consistently reduce the curse of dimensionality. In practice, literature works in the field of bioinformatics either avoid any dimensionality reduction [13] or make some empirical/heuristic decisions [14, 15] not motivated by any theoretical justification. However, the careful design of the DR step affects all the subsequent computations and the reliability of the obtained results [16, 17, 18].

Further, when applying dimensionality reduction in a multi-omics setting, few works consider that different views might carry different amounts of information. Neglecting this fact, most works blindly apply any of the successful data-fusion techniques proposed in literature [19, 11] (supplementary section S. A.3) without any prior view-reduction, or when a reduction is applied, the same (often empirical) dimension is chosen for all the views.

Instead, a prior view-specific reduction would better emphasize and expose the information within each view, therefore improving the effectiveness of the following data-fusion task, whose aim is to uncover the salient information across views, while removing the (between-)view redundancy [20, 21].

The aim of this work is to propose a novel block-analysis technique leveraging one of the most promising and recent Intrinsic Dimensionality (`id`) estimators ([22], supplementary section S. A.2), namely the *two-nn* estimator (supplementary section S. A.2.1) to understand when and how a feature selection or feature extraction method could improve the data representation. If curse of dimensionality is detected, the block-analysis allows defining the dimension of the lower-dimensional space where each view should be transformed by any of the promising feature selection or feature extraction approaches proposed at the state of the art (supplementary section S. A.1). Further, by exploiting the information provided by block-analysis, we propose and experiment with a novel two-step DR process that improves results by combining the advantages of the first application of feature selection followed by feature extraction.

The proposed DR technique is applied in the context of multi-omics data analysis, where effective multi-omics data-fusion algorithms have been recently developed. In particular, we compared some of the most promising and effective unsupervised multi-omics data-fusion techniques (i.e. MOFA+ [21], uMKL [23], and SNF [13], all summarized in supplementary section S. A.3) to assess their strengths and compare their robustness across different settings. Indeed, while the effectiveness of these data integration approaches is undoubted, we wanted to investigate (1) the effect of using subsets of the input multi-omics views, to understand whether a subset of the input views could suffice to provide effective results, or (2) whether the integration of not-omics patients’ views, e.g. patients’ demographics views carrying a completely different semantic, would enhance the salient and discriminative information, therefore facilitating the following supervised/unsupervised analysis.

To perform our comparative evaluation we selected nine (high-dimensional) multi-omics datasets from the well-known TCGA repository (more details in section 2) and designed a supervised machine learning pipeline that reduces all the omics views in the input dataset, fuses them (by eventually integrating also the demographic view or concatenating it to the integrated view), and finally uses a random forest classifier for analyzing the fused information to predict patients’ survival. By using a supervised classification task the obtained performance can be compared using well-established performance measures, such as the area under the ROC curve (AUROC) and the area under the Precision-Recall curve (AUCPR).

Results show that, in our classification problem, DR guided by block-analysis outperforms traditional DR approaches that use heuristics to set the dimensionality of the reduced space. Further, the robustness of DR improves when a two-step DR approach is applied, or when non-omics patients’ descriptors are also integrated into the analysis. On the other hand, when a prior (and properly designed) step of DR is applied to effectively remove intra-view redundancy and noise, there is no need to spare computational time for testing the usage of subsets of the input views, because the data-fusion algorithms can produce effective integrated representations that achieve robust classification results.

## 2. TCGA datasets

To obtain reliable results, we mined the following nine multi-omics datasets from the TCGA cancer repository^1^ (see tables 1 and 2): the BLadder urothelial Carcinoma dataset (**BLCA**); the BReast infiltrating ductal CArcinoma (**BRCA1**) and the BReast infiltrating lobular CArcinoma (**BRCA2**) datasets, composed by splitting all the samples in the BReast nvasive CArcinoma dataset (BRCA); the KIdney Renal Clear cell carcinoma dataset (**KIRC**); the LUng ADenocarcinoma dataset (**LUAD**); the LUng Squamous Cell carcinoma datset (**LUSC**); the PRostate ADenocarcinoma dataset (**PRAD**); the OVarian serous cystadenocarcinoma dataset (**OV**); the SKin Cutaneous Melanoma dataset (**SKCM**).

**Table 1:** Descriptive statistics for BLCA, BRCA1, BRCA2, KIRC, and LUAD datasets. Column *N* reports the number of cases; column *D* (raw) reports the original dimension of each view; column *D* reports the dimension of each view after data pre-filtering to remove noise and high pairwise-redundancy (see supplementary file S. B); column 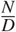 reports the ratio between the number of cases and the dimension of each view; columns *N*_*neg*_, *N*_*pos*_, and 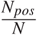 report, respectively, the number of negative (OS = 0) and positive (OS = 1) patients, and the balance ratio, measured as the ratio between the number of positive cases and all the cases in the dataset. “methy” stands for DNA methylation data.

| **Dataset** | **view** | **N** | **D (raw)** | **D** | $\frac{N}{D}$ | $N_{pos}$ | $N_{neg}$ | $\frac{N_{pos}}{N}$ |
| --- | --- | --- | --- | --- | --- | --- | --- | --- |
| **BLCA** | miRNA | 335 | 469 | 469 | 0.7143 | 151 | 184 | 0.45 |
|  | mRNA |  | 12276 | 12276 | 0.027 |  |  |  |
|  | proteins |  | 183 | 183 | 1.831 |  |  |  |
|  | methy |  | 315551 | 30000 | 0.012 |  |  |  |
| **BRCA1** | miRNA | 317 | 496 | 496 | 0.6391 | 42 | 275 | 0.13 |
|  | mRNA |  | 12242 | 12242 | 0.026 |  |  |  |
|  | proteins |  | 202 | 202 | 1.569 |  |  |  |
|  | methy |  | 289962 | 30000 | 0.011 |  |  |  |
| **BRCA2** | miRNA | 128 | 502 | 502 | 0.255 | 14 | 114 | 0.11 |
|  | mRNA |  | 8128 | 8128 | 0.016 |  |  |  |
|  | proteins |  | 192 | 192 | 0.667 |  |  |  |
|  | methy |  | 278099 | 30000 | 0.004 |  |  |  |
| **KIRC** | miRNA | 169 | 364 | 364 | 0.464 | 48 | 121 | 0.28 |
|  | mRNA |  | 7942 | 7942 | 0.021 |  |  |  |
|  | proteins |  | 186 | 186 | 0.909 |  |  |  |
|  | methy |  | 319740 | 30000 | 0.006 |  |  |  |
| **LUAD** | miRNA | 300 | 465 | 465 | 0.645 | 120 | 180 | 0.40 |
|  | mRNA |  | 11131 | 11131 | 0.027 |  |  |  |
|  | proteins |  | 179 | 179 | 1.676 |  |  |  |
|  | methy |  | 331828 | 30000 | 0.01 |  |  |  |

**Table 2:**
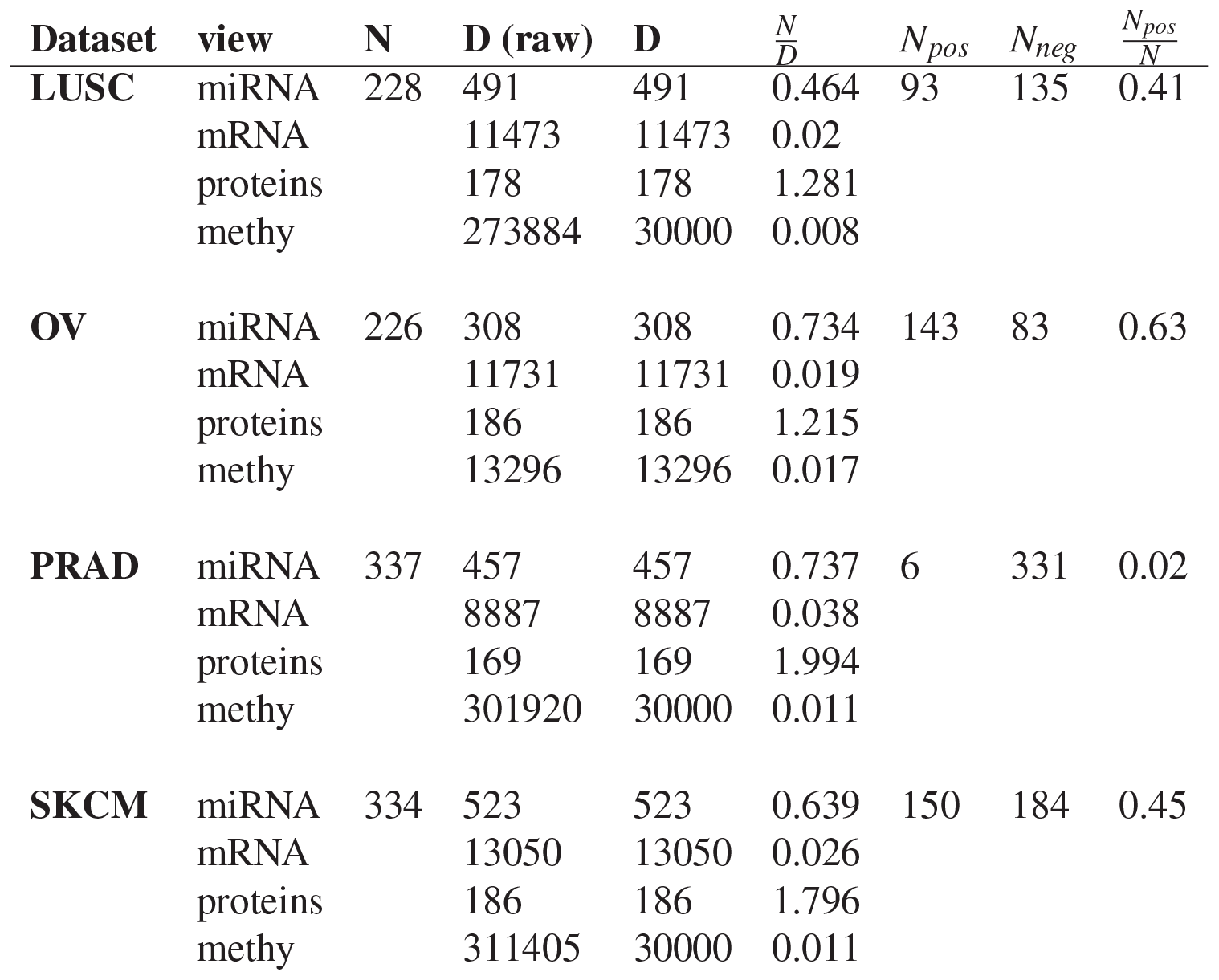
Descriptive statistics for LUSC, OV, PRAD, and SKCM datasets.

| **Dataset** | **view** | **N** | **D (raw)** | **D** | $\frac{N}{D}$ | $N_{pos}$ | $N_{neg}$ | $\frac{N_{pos}}{N}$ |
| --- | --- | --- | --- | --- | --- | --- | --- | --- |
| **LUSC** | miRNA | 228 | 491 | 491 | 0.464 | 93 | 135 | 0.41 |
|  | mRNA |  | 11473 | 11473 | 0.02 |  |  |  |
|  | proteins |  | 178 | 178 | 1.281 |  |  |  |
|  | methy |  | 273884 | 30000 | 0.008 |  |  |  |
| **OV** | miRNA | 226 | 308 | 308 | 0.734 | 143 | 83 | 0.63 |
|  | mRNA |  | 11731 | 11731 | 0.019 |  |  |  |
|  | proteins |  | 186 | 186 | 1.215 |  |  |  |
|  | methy |  | 13296 | 13296 | 0.017 |  |  |  |
| **PRAD** | miRNA | 337 | 457 | 457 | 0.737 | 6 | 331 | 0.02 |
|  | mRNA |  | 8887 | 8887 | 0.038 |  |  |  |
|  | proteins |  | 169 | 169 | 1.994 |  |  |  |
|  | methy |  | 301920 | 30000 | 0.011 |  |  |  |
| **SKCM** | miRNA | 334 | 523 | 523 | 0.639 | 150 | 184 | 0.45 |
|  | mRNA |  | 13050 | 13050 | 0.026 |  |  |  |
|  | proteins |  | 186 | 186 | 1.796 |  |  |  |
|  | methy |  | 311405 | 30000 | 0.011 |  |  |  |

For each dataset, we considered miRNA and mRNA (RNA-Sequencing expression values), protein expression (Reverse Phase Protein Arrays), and DNA methylation (Methylation Array) views, which were pre-processed to filter variables mainly carrying noise or highly redundant information (see supplementary section S. B for further details).

We also complemented the omics information with demographic patient data (age at first pathological diagnosis, gender, race, ethnicity, see supplementary tables S. B.1-S. B.3 for further details). Patients in the TCGA dataset may be classified based on their Overall Survival (OS) event, which is available from the TCGA-CDR [25] dataset. We used the overall survival label to perform a supervised classification task.

Note that some literature studies using TCGA datasets for testing classification models [26, 27, 28, 29, 30] already exist. However, these studies typically restrict their analysis to a maximum of four TCGA datasets, without providing clear justification for their choices. In contrast, our approach involved the selection of nine diverse datasets, that were chosen to encompass a wide range of heterogeneity, not only in terms of different tumor types being investigated, but also in the ratio between the number of cases and variables within each dataset, and the balance between positive (patients with OS event equal to 1) and negative patients (OS = 0). By adopting this comprehensive approach, we aimed to capture a more nuanced and representative perspective in our analysis.

## 3. Dimensionality reduction approach

In this section, we describe the block-analysis we propose to provide unbiased estimates of the `id` of a data-view (subsection 3.1).

The automated analysis of the block-`id` distribution provides a quantitative information about the amount of feature noise and redundancy affecting the view (subsection 3.2) and, based on that, it allows tailoring the dimensionality reduction of the analyzed view.

To guide the reader, Figure 1 sketches the DR pipeline guided by block-analysis.

**Figure 1:**
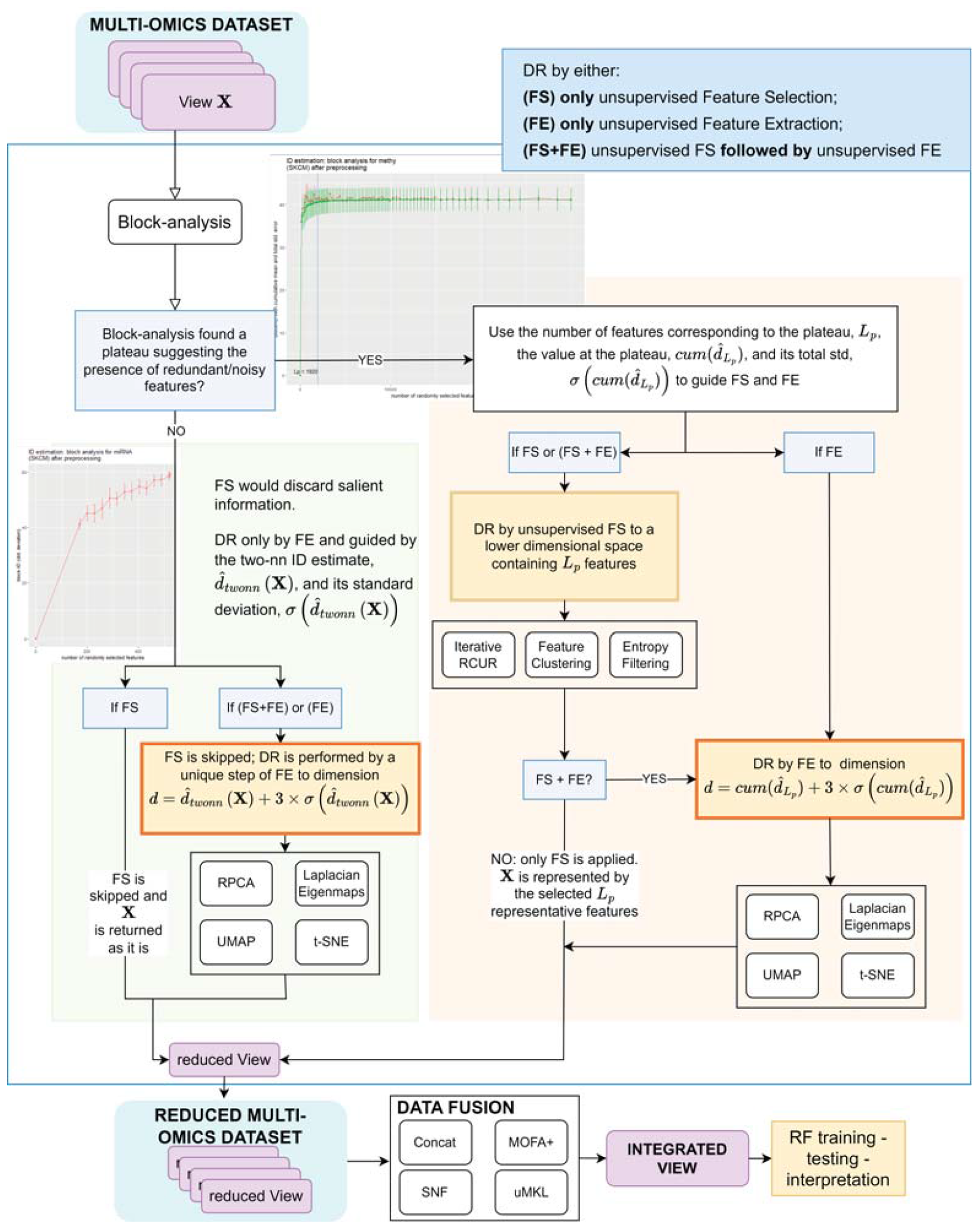
Experimented DR and data-integration pipelines.

In the whole section, we consider an input view (dataset), **X** *∈* ℜ^*N×D*^, with *N* being the number of cases, and *D* the number of features (dimension) of the view.

### 3.1. Block analysis and block-ID estimate

Several of the most promising `id` estimators produce unstable global estimates on real datasets, often affected by the small-sample-size causing sample-sparsity, outliers, and noise [31, 32, 22] (see supplementary section S. A.2 for further details). This is particularly true for nearest-neighbor `id` estimators, which base their estimation on the analysis of the distribution of points withing small data-neighborhoods. Due to the unreliability of pairwise-distances in datasets characterized by the small-sample-size, these estimators often suffer from high variance or overestimation when, e.g., the considered point-neighborhood size increases. Furthermore, since all the `id` estimators contain some randomness, most of them suffer from an added factor of variance, particularly evident when working in high dimensions.

To account for such variance as well as the presence of outlier and boundary points that could bias the estimates, authors of *two-nn* [22] proposed experiments on simulated datasets (with a large number of samples, i.e. not affected by the small-sample-size) where they apply a classic block-analysis [33]. In particular, authors compute (sub-optimal) `id` estimates (and their standard deviation) by averaging the estimates obtained on under-sampled, non-intersecting datasets composed of a number *n < N* of samples. Plotting the distribution of the obtained estimates for increasing values of *n*, a plateau is found, corresponding to an unbiased (optimal) estimate of the `id` characterizing the informative content of the dataset.

The above-mentioned approach is effective on simulated experiments, where enough samples can be generated to avoid the curse of dimensionality. On the other hand, when dealing with real bio-medical datasets, often limited in samplesize and potentially affected by the curse of dimensionality, we propose to reduce the bias due to the presence of noisy and outlier points by averaging all the *two-nn* `id` estimates computed on *M* under-sampled versions of the dataset, where the under-sampling randomly selects (with repetition) a fixed percentage, *t*, of the dataset points^2^. Choosing a proper value for the percentage *t* allows to have enough samples in each sub-dataset *M*, so that the average (and the standard deviation) of all the *M* `id`-estimates may be a first, more robust, *two-nn* `id`-estimate (and standard deviation of the estimate) of the input dataset.

In the following, any reference to the *two-nn* `id`-estimate of a dataset **X**, 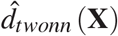 (and its standard deviation 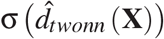 refers to this unbiased estimate.

While the aforementioned procedure mitigates the problems affecting real, noisy datasets, it still cannot cope with the possible curse of dimensionality, which practically shows up with a large number of features being noisy or redundant. Unfortunately, given an input view **X** *∈* ℜ^*N×D*^ there is no rule of thumb for deciding when a dataset characterized by low values of the ratio 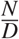 is affected by the curse of dimensionality. To provide such understanding and to obtain an unbiased `id`-estimate of the view even in the presence of noisy and redundant features we propose applying the block-analysis feature-wise, as detailed in this section.

In particular, we start by using the *two-nn* `id`-estimate for the input view, 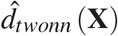, to set the dimension *L*_0_ of the smaller block as 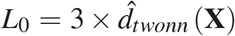. Though we are aware that this `id` estimate might still be biased by redundant and noisy features, if any, it can be a valid aid to guarantee that even smaller blocks can contain enough information to produce reliable estimates.

Once *L*_0_ is set, we perform the block-analysis by iterating over blocks with increasing dimensions, estimating the *two-nn* `id` of each block, and then analyzing the distribution of all the block-`id`s.

More precisely, at the *j*^*th*^ iteration (*j*-th block **B** _*j*_), when the block size is *L*_*j*_ = *L*_0_ + *j ×L*_0_, *L*_*j*_ ≤ *D*, we estimate the `id` (and its fluctuations) for **B** _*j*_ by:

I. creating *n*_*try*_ blocks, 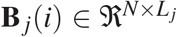, *i ∈* [1,…, *n*_*try*_], each representing all the samples in the input view with *L*_*j*_ randomly sampled features;
II. estimating the *two-nn* `id` of each **B** _*j*_(*i*), 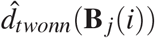 and then computing the mean (and variance) of all the computed estimates to obtain the block-`id` estimate, 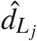 (and its variance, 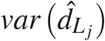 for **B** _*j*_, being *var* the variance operator.

This step essentially provides an estimate of the `id` (and its variance) that would be obtained if the data was represented by *L*_*j*_ randomly selected features^3^.

### 3.2. Automatic analysis of Block-ids allows tailoring dimensionality reduction

The red dotted lines in figure 2 (and supplementary figures S. C.1-S. C.4) plot the block-`id`s, 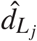 for increasing block dimensions in the miRNA and pro-tein views (SKCM dataset); red bars in the figure repres<u>ent stand</u>ard deviations, computed as the square root of the variance 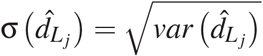.

**Figure 2:**
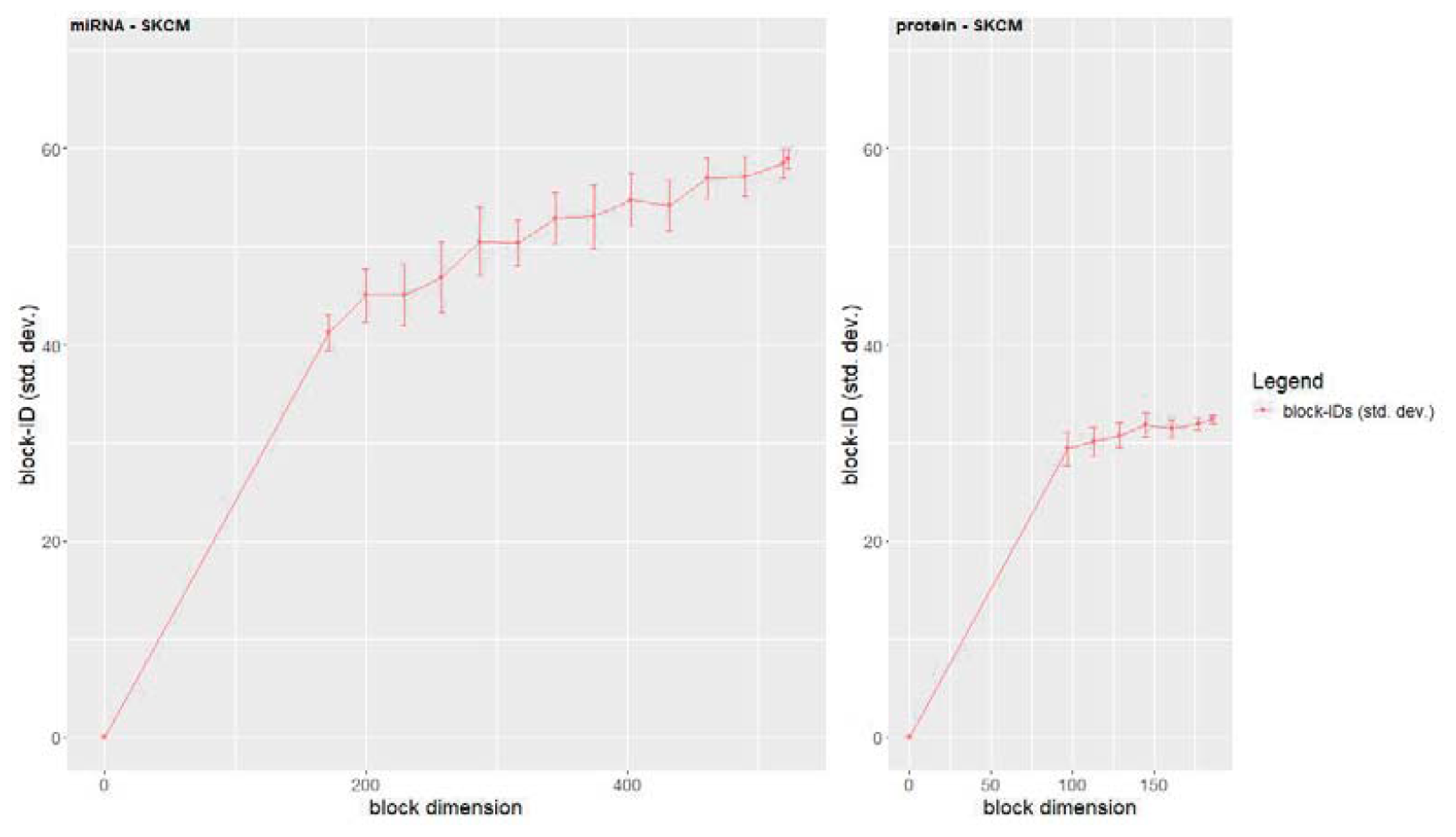
Block-analysis performed by using the *two-nn* estimator on the SKCM dataset. Left: miRNA view (SKCM dataset). Right: protein view (SKCM dataset). Point *L*_*j*_ of the red-dotted line (and the vertical bars) represents 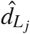 (and its standard deviation 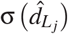, that is the estimated block-id for block **B** _*j*_, computed as the mean (and standard deviation) of the ids estimated on *n*_*try*_ blocks with dimension *L*_*j*_ (and its standard deviation). The block-id increases as the block dimension increases, suggesting that each of the added features increases the amount of information. Therefore, the id of the whole view, i.e. the id of the block covering all the features, is a reliable estimate of the dimensionality of the space where the data should be transformed by a feature-extraction algorithm (figure 1 - light green box - FE option). On the other hand, considering that each feature adds novel information, if feature-selection is the chosen dimensionality reduction approach (figure 1 - light green box - FS option), the view is not reduced and it is returned as it is; in other words, feature-selection is avoided because it would necessarily spare information.

When observing the miRNA and protein views, which are characterized by higher ratios 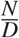 when compared to the mRNA and methylation data-views, we note that the block-`id` keeps increasing until the block size includes all the features in the view, *L*_*j*_ = *D*, that is, the block-`id` equals the *two-nn* `id`-estimate of the whole view: 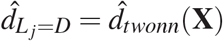 and 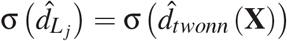.

This suggests that each new feature adds novel information. In other words, the view contains a limited amount of noise and redundancy, supposedly due to the data-view belonging to a real dataset. In this case, the *two-nn*-`id` of the whole view, 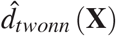 (and its standard deviation, 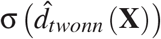 is an unbiased estimate of the `id` of the whole view (and its fluctuations).

In practice, *when no plateau is automatically detected by block-analysis* (see figure 1 - light green box) no DR via feature-selection is applied because the selection of a subset of features would surely cause loss of information. Instead, we allow performing DR via feature extraction, which considers (and combines) all the features in the dataset (all the original information in the dataset) while computing the reduced view. In this case, the dimension of the reduced space, *d* where the view is transformed is computed by using the `id` estimate of the whole view: 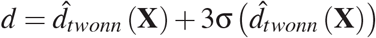.

On the other hand, for the mRNA and the methylation view (figure 3) the distribution of the block-`id`s (red-dotted line) is more noisy, and increases until it reaches a (noisy) plateau. The dimension *L*_*p*_ of the block where the plateau starts (horizontal axis in figure 3, automatically detected as described in supplementary section S. C) can be regarded as an estimate of the minimum number of (salient) features that can be used to represent the salient information in the data-view, and after which the addition of extra features mainly adds redundancy and/or noise.

**Figure 3:**
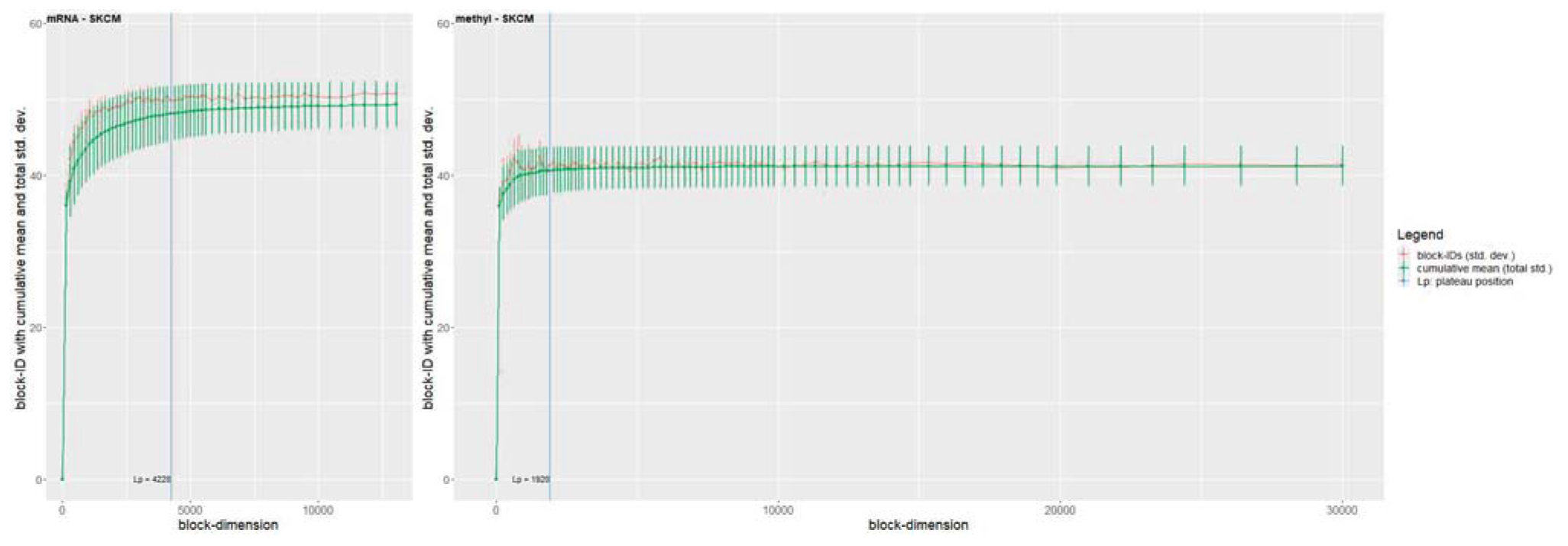
Block-analysis performed by using the *two-nn* estimator on the SKCM dataset. Left: mRNA; right: methylation data. The block-ids (red-dotted line) are more noisy than those in the miRNA and protein views. The effect of noise is reduced by the computation of the cumulative mean (green-dotted line), soon reaching stability (plateau), providing a reliable estimate of the view id. The analysis of the cumulative mean allows the automatic detection of the position of the plateau (*L*_*p*_ = 4228 and *L*_*p*_ = 1920 for, respectively, the mRNA and the methylation view), corresponding to the number of features that may be selected from the dataset to reduce the information loss. In other words, the block-analysis of the mRNA and methylation views allows to detect signs of the curse of dimensionality in terms of the presence of feature redundancy. Moreover, it provides an unbiased estimate of the id characterizing the information content of view, and an estimate of the number of features that could be retained by any unsupervised feature selection algorithm to avoid information loss.

In practice, *L*_*p*_ is the number of the original features to be selected by an unsupervised feature selection algorithm to reduce noise and redundancy. While this step reduces the curse of dimensionality effects, the value of *L*_*p*_ is often high. To reduce the computational costs of the following algorithms and compute a data-representation concisely conveying similar information, we therefore propose applying a **two-step DR** approach where the reduced *L*_*p*_-dimensional view is input to a feature extraction algorithm^4^.

The feature extraction transforms the data into a space whose dimension, *d*, is computed based on an unbiased estimate of the view `id`, computed by considering that the block-`id` distribution is very noisy for views affected by the curse of dimensionality. To reduce noise effects by averaging, we compute the cumulative mean of the block-`id`s (green-dotted line in figure 3).

More precisely, the cumulative mean for block **B** _*j*_, 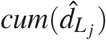, is computed as the average of all the block-`id`s computed for blocks 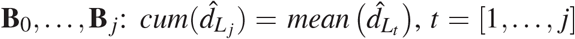. Eve’s law of total variance [34] allows computing the total variance of 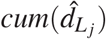, as the sum of the (unexplained) variance 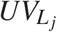 due to the `id`-estimator, and the (explained) variance 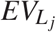 due to the sampling process^5^.

In practice, each point of the cumulative mean represents the average block-`id` that would be obtained on a view composed by randomly sampling a number of features that is equal to (or lower than) the dimension of block **B** _*j*_.

Further, assuming some features are mostly carrying noise and/or redundant information, the random under-sampling of features that is performed to compose blocks with varied and increasing dimensions, as well as the evaluation of the `id` for increasing block dimensions, is able to reduce (by averaging) biasing effects due to noise and redundancy. This is also visible in the plot of the cumulative mean, which approaches the block-`id` (red-dotted line) plot and is less noisy. This suggests that the (cumulative) value corresponding to the plateau of the block-`id`, 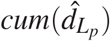, and its total standard deviation, 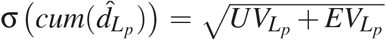, can be considered as an unbiased estimate of the `id` (and its fluctuations) of the whole view.

Summarizing, *when a plateau is automatically detected in position L*_*p*_ *of the block-*`id` *distribution* (see figure 1 - light orange box) we reduce the curse of dimensionality by applying any of the following three DR options: (1) if feature selection is the preferred approach, we select *L*_*p*_ salient features; (2) if feature extraction is the preferred DR approach, the data-view is transformed to a lower dimensional space with dimension 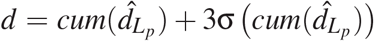; (3) if a two phase DR is chosen, feature selection is applied to select *L*_*p*_ features and the re-duced view is input to a feature extraction algorithm that transforms the dataset into a space with dimension 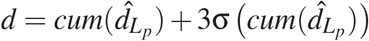.

Tables 3 and 4 report, for each dataset and view used in our experiments (section 2), the number of features *L*_*p*_ corresponding to the plateau, if any is found, the `id` estimate 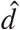, which equals either 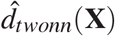 - when no plateau is found - or 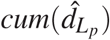 - when a plateau is found, and its total standard deviation, 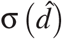, computed as the square root of the total variance, 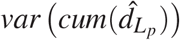.

**Table 3:** Block-id estimates for BLCA, BRCA1, BRCA2, KIRC and LUAD datasets. For each dataset-view the table reports the number of features *L*_*p*_ corresponding to the plateau (if any is found), and the id estimate 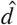 with its corresponding standard deviation 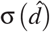.

| **Dataset** | **view** | $L_p$ | $\hat{d}$ | $\sigma(\hat{d})$ |
| --- | --- | --- | --- | --- |
| **BLCA** | miRNA |  | 53.72 | 1.81 |
|  | mRNA | 2835 | 42.75 | 2.85 |
|  | proteins |  | 28.83 | 1.19 |
|  | methy | 4238 | 52 | 3.48 |
| **BRCA1** | miRNA |  | 51.11 | 1.68 |
|  | mRNA | 4640 | 46.61 | 3.18 |
|  | proteins |  | 35.04 | 1.26 |
|  | methy | 5738 | 48.57 | 3.55 |
| **BRCA2** | miRNA |  | 35.21 | 3.69 |
|  | mRNA | 4216 | 42.26 | 3.57 |
|  | proteins |  | 27.75 | 1.23 |
|  | methy | 3000 | 39.87 | 3.04 |
| **KIRC** | miRNA |  | 53.26 | 5.55 |
|  | mRNA | 4340 | 39.28 | 3.35 |
|  | proteins |  | 31.1 | 1.6 |
|  | methy | 2208 | 47.35 | 4.74 |
| **LUAD** | miRNA |  | 44.15 | 1.12 |
|  | mRNA | 3168 | 43.32 | 2.72 |
|  | proteins |  | 33.26 | 1.44 |
|  | methy | 1677 | 43.27 | 3.36 |

**Table 4:** Block-id estimates for LUSC, OV, PRAD and SKCM datasets. For each dataset-view the table reports the number of features *L*_*p*_ corresponding to the plateau (if any is found), and the id estimate 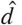 with its corresponding standard deviation 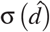.

| **Dataset** | **view** | $L_p$ | $\hat{d}$ | |
| --- | --- | --- | --- | --- |
| **LUSC** | miRNA |  | 50.77 | 1.41 |
|  | mRNA | 4350 | 48.38 | 3.55 |
|  | proteins |  | 35.79 | 0.84 |
|  | methy | 3400 | 43.19 | 3.26 |
| **OV** | miRNA |  | 33.97 | 2.22 |
|  | mRNA | 3173 | 55.19 | 4.78 |
|  | proteins |  | 38.07 | 2.08 |
|  | methy | 3614 | 44.14 | 2.93 |
| **PRAD** | miRNA |  | 54.04 | 1.95 |
|  | mRNA | 3645 | 43.57 | 2.66 |
|  | proteins |  | 27.43 | 0.97 |
|  | methy | 7760 | 61.92 | 4.23 |
| **SKCM** | miRNA |  | 57.11 | 3.14 |
|  | mRNA | 4228 | 48.6 | 3.39 |
|  | RPPA |  | 32.45 | 1.07 |
|  | methy | 1920 | 40.86 | 2.88 |

## 4. Results

In this section we first summarize the DR+data-fusion pipelines we devised and experimented (subsection 4.1); next, we detail the experimental settings of the supervised classification task we exploited to objectively compare the different pipelines (subsection 4.2); finally we report and discuss the results obtained by our comparative evaluation (subsection 4.3).

### 4.1. Dimensionality reduction guided by block-analysis and multi-omics data fusion

In this section, we detail the (one-step or two-step) DR+data-fusion pipelines we designed and compared by their application for supervised prediction. To help readers’ comprehension, figure 1 sketches all of them.

DR is performed by either a unique step of unsupervised feature selection (FS in figure 1), a unique step of unsupervised feature extraction (FE in figure 1), or by a 2-step DR process where the ouput of feature selection is input to feature extraction (FS+FE in figure 1).

The unsupervised feature selection algorithms we adopt were chosen based on their documented promising results, their limited computational costs, and considering preliminary experiments we ran, which showed their robustness with respect to datasets characterized by a limited cardinality. For interested readers, a brief literature background about unsupervised feature selection is reported in supplementary section S. A.1.1. In particular, the following algorithms were selected, and eventually optimized to reduce their computational costs:

- A parallel feature clustering algorithm returning the features that are the centroids of the identified clusters. Given the high computational costs of feature clustering methods at the state-of-the-art, the algorithm we implemented splits the input view into non-intersecting feature subsets that are distributed on multiple cores. Each core applies the Genie agglomerative clustering algorithm [35] to cluster the input feature subset, and then returns the features that are centroids of each cluster (feature medoids). The main algorithm recollects and concatenates all the feature medoids and iterates the algorithm on the concatenated feature medoids to perform a further selection until a number *L*_*p*_ of feature medoids is reached. More details are reported in supplementary section S. D.1.
- An iterative version of the RCUR [36] algorithm, whose parallel schema is similar to the one applied for feature clustering (more details are reported in supplementary section S. E); it allows selecting an *L*_*p*_-dimensional subset of the original features, based on their potential to represent the information in the input view.
- A simple entropy filtering algorithm that selects the *L*_*p*_ features with the highest entropy.

When a unique step of unsupervised feature selection is applied to reduce all the multi-omics views in the input dataset, only views for which the block-analysis identified a plateau (that is, views affected by feature redundancy - light red box in figure 1) are reduced by selecting a number of features corresponding to the position *L*_*p*_ of the plateau of the block-`id`. The other views are kept as they are to avoid loss of information (light-green box in figure 1).

Feature-extraction algorithms were similarly chosen based on their promising and successful results (supplementary section S. A.1.2). In detail, we compared Randomized PCA (RPCA, alias RSVD), laplacian eigenmaps, UMAP, and t-SNE and defined the dimension of the lower-dimensional space as 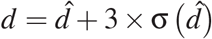, where 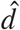 (and 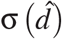) is the estimated `id` (and its standard deviation). We recall that (section 3.1) a reliable `id` estimate (and standard deviation) of views not affected by feature redundancy is obtained as the mean (and standard deviation) of the *two-nn* estimates computed on *M* undersampled sets (light-green box in figure 1). When instead the block-analysis detects feature redundancy (light-red box in figure 1), the cumulative value at the plateau of the block-`id` distribution (and its total standard deviation) provides a reliable `id` estimate.

When we apply the two-step DR pipelines we simply perform a preliminary unsupervised feature selection method among those listed above followed by any of the unsupervised feature extraction methods listed above. This practically means that only views where a plateau is found (light-red box in figure 1) undergo feature-selection and then feature-extraction; the other views undergo only feature-extraction.

Once all the views in the input dataset have been individually reduced, they are input to a data-fusion algorithm to leverage the related and complementary information across views and produce an integrated view that may be input to any further analysis. Besides the basic concatenation of the reduced views, which can be considered as a simple benchmark for comparison, we exploited and compared data-fusion approaches that showed their promise in several multi-omics data-analysis tasks and are applied in different stages of the data analysis [37, 11] (details in supplementary section S. A.3). In particular, we experimented with: (1) an input-data fusion technique, namely MOFA+ [21], which applies a Bayesian approach to derive a set of latent factors capturing and representing the information content of the input multi-modal representation; two Patient-Similarity-Network (PSN) fusion techniques, that are (2a) the widely used Similarity Network Fusion algorithm (SNF, [13]), which applies a smart diffusion process that merges the similarities between pairs of samples that have “shared” neighbors across views, and (2b) an unsupervised Multiple Kernel Learning technique (uMKL, [23]), which outputs the (integrated) kernel that best aligns with all the unimodal kernels (i.e. the Gram matrices) representing the topological structure of each input view.

Overall, we experimented with nineteen DR methods; seven of them (one-step DR approach) applied either one of the three unsupervised feature-selection methods (feature clustering, iterative RCUR - *par rcur* in the following, entropy filtering) or four unsupervised feature-extraction methods (RPCA, Laplacian Eigenmaps, t-SNE, and UMAP); twelve were two-step DR pipelines obtained by all the combinations of the four feature-selection algorithms and the three feature-extraction algorithms. Considering that the reduced data is input to any of the four data integration methods we experimented (SNF, uMKL, MOFA+, and the simple concatenation), for each multi-omics dataset we run about eighty different DR+data-fusion pipelines (experiments).

### 4.2. Experimental settings

Each DR+data-fusion pipeline was tested on a binary classification task across all the nine multi-omics cancer datasets (section 2). To this aim, a random forest classifier (RF, [38]) was trained and tested to predict the overall survival event of patients.

Besides their interpretable nature [39], their often superior effectiveness with respect to even the (less efficient) deep neural network models [40], and their capability of handling a set of heterogeneous variables [38], we chose RF classifiers due to their robustness to the input feature set and the choice of hyper-parameter values. This makes it easier to apply them consistently across different datasets, DR, and data-fusion approaches, and allows an objective assessment of the informativeness of the (reduced) input-data representation and the effectiveness of the data-fusion algorithms, without the confounding effects due to the prior application of supervised feature selection or hyper-parameter tuning steps^6^.

To obtain an unbiased evaluation, the RF training and testing phase was repeated across fifteen stratified holdouts (80:20 train:test ratio) that obviously differed for each dataset but were kept fixed across all the experiments run on the same dataset. To avoid confounding effects that could hamper an objective comparison, we avoided the application of any supervised feature selection algorithm and we instead set all the RF parameters to their default values.

Paired-samples Wilcoxon test, alias Wilcoxon signed-rank test, at the 95% of confidence (i.e. α = 0.05) was used for comparison. If not specified, the test was performed by pooling the results obtained on all the nine TCGA datasets (supplementary files report also the details per dataset); for each comparison, we considered AUCPR and AUC for hypothesis testing and exploited win-tie-loss tables to summarize the statistical comparison between each method against all the others^7^. In particular, when two specific DR+data-fusion experiments were compared, we paired the results obtained on each of the nine TCGA datasets and the fifteen stratified holdouts. When, instead, we performed more generic comparisons to asses each DR approach (or each data-fusion method), we paired the results obtained across the nine different datasets, the fifteen holdouts, and the four data-fusion methods (or nineteen DR pipelines).

Wilcoxon signed-rank test summarize and compare the performance of different pipelines across multiple settings. Therefore, pipelines that achieve the highest/lowest number of wins/losses can be regarded as being, on the average of all the experimented settings, the top-performing and most robust.

However, under specific settings, some other pipelines may achieve promising results; to provide a more exhaustive and detailed description of the obtained results, for each of the considered comparative evaluations we collected and analyzed the list of the top-performing experiments; in simpler words, for each of the nine TCGA datasets, we collected the three (DR+data-fusion) experiments that obtained the highest AUC or AUCPR values. This allowed counting the frequency of the DR and data-fusion pipelines occurring among the top-performers.

### 4.3. Comparative evaluation results

After performing data-fusion tests with no prior DR that evidenced the need of a properly designed DR approach (supplementary section S. F.1), we conducted tests to compare the proposed DR approach (guided by block-analysis) to methods that exploit heuristics or empirical measures to set the dimension of the lower dimensional space (section 4.3.1). Next, we compared the results obtained by the block-analysis guided DR+data-fusion pipelines to gain insights about different data-fusion settings, ranging from the traditional multi-omics fusion setting (subsection 4.3.2), to those settings where we fused subsets of omics (subsection 4.3.3), and omics plus non-omics views (subsection 4.3.4).

#### 4.3.1. When compared to heuristics, the usage of block-analysis and the id estimate obtain better results

Besides comparing the described DR+data-fusion pipelines, in our experiments we also aimed to assess the effectiveness of using the `id` estimate to set the dimension of the lower dimensional space.

To this aim, we initially experimented with DR pipelines exploiting a unique step of either feature-selection or feature-extraction (1-step DR, section 4.1) to choose the better performing among two heuristically set dimensions, 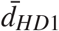 and 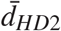. In particular, the heuristic dimensions we chose to compare are based on the rationale that most of the feature extraction algorithms allow to compute a reduced space whose number of dimensions is lower or equal than *min*(*N, D*) − 1 [36, 41, 42]. Based on this consideration, we run all the one-step DR+data-fusion pipelines by using two heuristics, *HD*_1_ and *HD*_2_. In particular, *HD*_1_ sets the dimension of the reduced space to 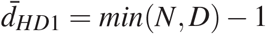; *HD*_2_ halves 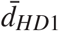, i.e. for *HD*_2_ we used 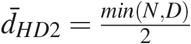.

In supplementary section S. F.2 we report details about the experiments we performed to compare the results obtained by using *HD*_1_, *HD*_2_, and our block-analysis to guide the reduction. Besides avoiding empirical or heuristic choices, the assessment showed the promise of our proposal.

Further, the obtained results hint that the most robust and effective results are obtained by a two-step DR pipeline combining the iterative version of RCUR we implemented with RPCA.

On the other hand, when comparing the performance and robustness of the data-fusion algorithms, SNF is undoubtedly among the most promising techniques in all the comparative evaluations; howver, also uMKL and MOFA+ show their promise.

#### 4.3.2. Comparison of DR and data integration pipelines guided by the block-analysis

To assess and compare the robustness of the DR+data-fusion pipelines guided by block analysis we applied the paired samples Wilcoxon test to compare: (1) each DR pipeline against each other, by pairing all results across datasets, hold-outs, and data-fusion algorithms; (2) each data-fusion algorithm against each other, by pairing all results across datasets, holdouts, and DR pipelines. Figures 4 and figure 5 show the win-tie-loss tables obtained when the AUC or the AUCPR measures are used for comparison.

**Figure 4:**
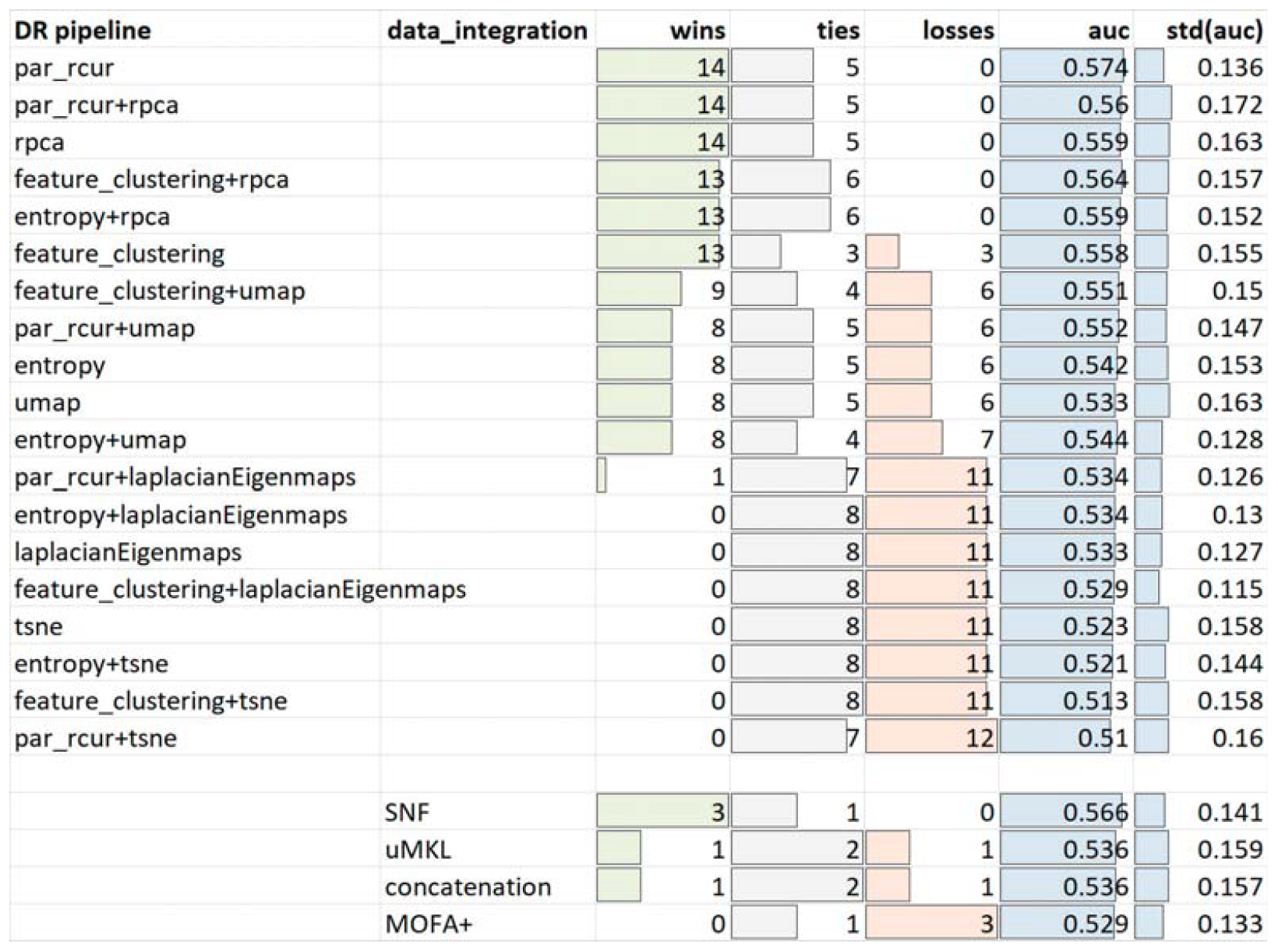
Win-tie-loss tables obtained by Wilcoxon signed-rank test when using the AUC measure to compare (top table) all the DR pipelines exploiting the information provided by block-analysis to guide the reduction of each of the four omics views, and (bottom table) the data-fusion algorithms that fuse the reduced views (bottom table).

**Figure 5:**
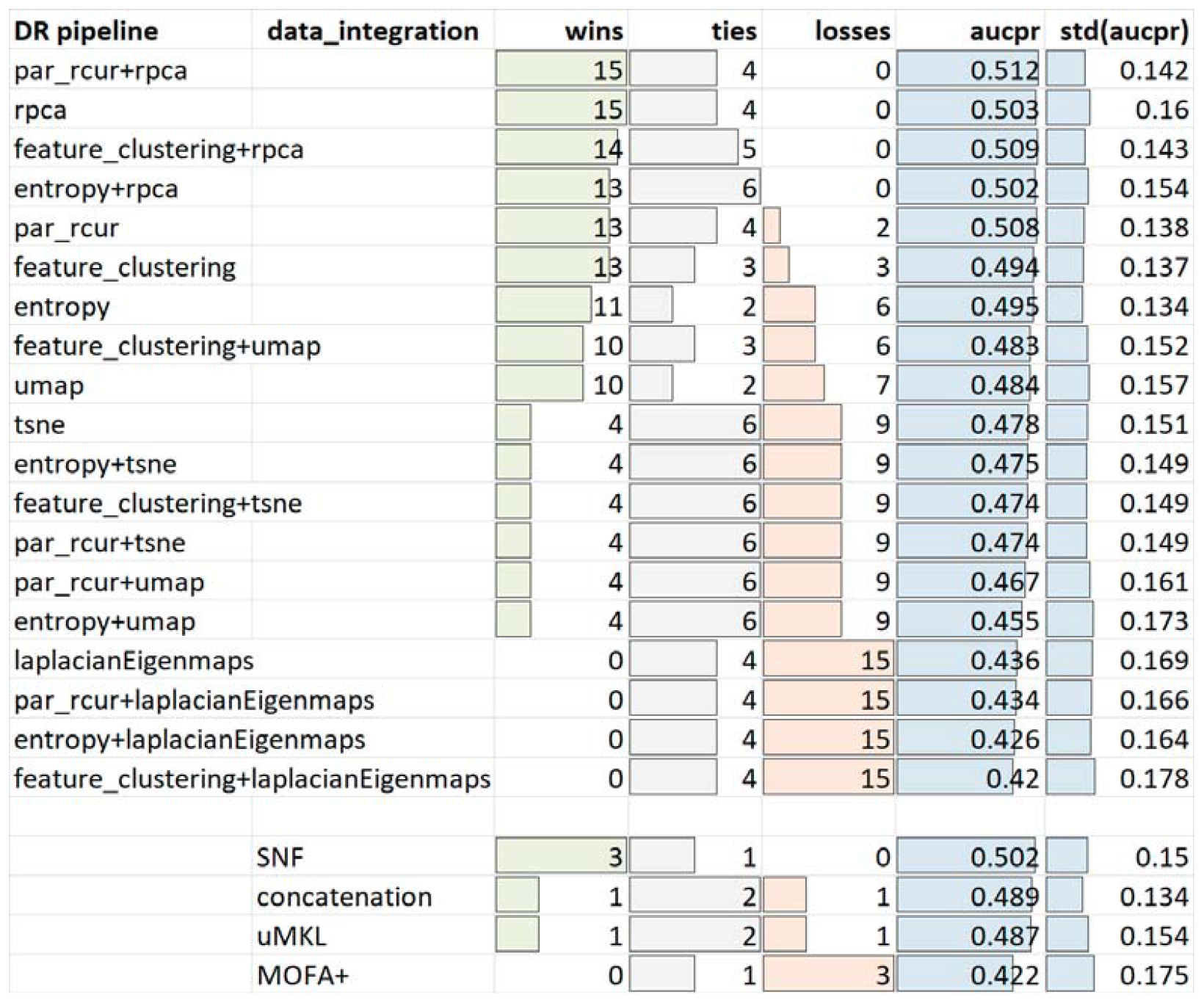
Win-tie-loss tables computed by Wilcoxon signed-rank test when using the AUCPR measure to compare all the DR pipelines guided by block-analysis (top table) and the data-fusion algorithms (bottom table). All the four omics are reduced and then integrated.

For what regards DR approaches, the two-step DR approaches using RPCA are in the list of top-winning pipelines that have zero losses (three up to five approaches when the AUC is used, and, most importantly, three up to four approaches when the AUCPR measure is used); generally speaking, all DR methods exploiting RPCA are the top-winners, confirming the experiments reported in [18]. The superiority of two-step DR approaches using RPCA was also confirmed by the Wilcoxon signed-rank tests we ran to compare each DR+data-fusion pipeline against each other (supplementary figures S. F.13 and S. F.14 and supplementary tables S7 and S8).

Among the data integration methods, SNF seems the most robust algorithm with respect to different settings. However, when observing the pairwise DR+datafusion comparisons where the AUCPR is used as the evaluation measure (supplementary figure S. F.14), uMKL also shows its promise.

Further, for each dataset, we collected the top-performing pipelines, that is the list of DR+data-fusion pipelines that obtain the three highest AUCs or AUCPRs. Next, we counted the frequency of occurrence of each DR and data-fusion algorithm in the top-performing list (the detailed list of top-performers is reported in supplementary file S9). Figure 6 shows that the majority of top-performers use a two-step reduction schema, including RPCA and fusing data by means of SNF.

**Figure 6:**
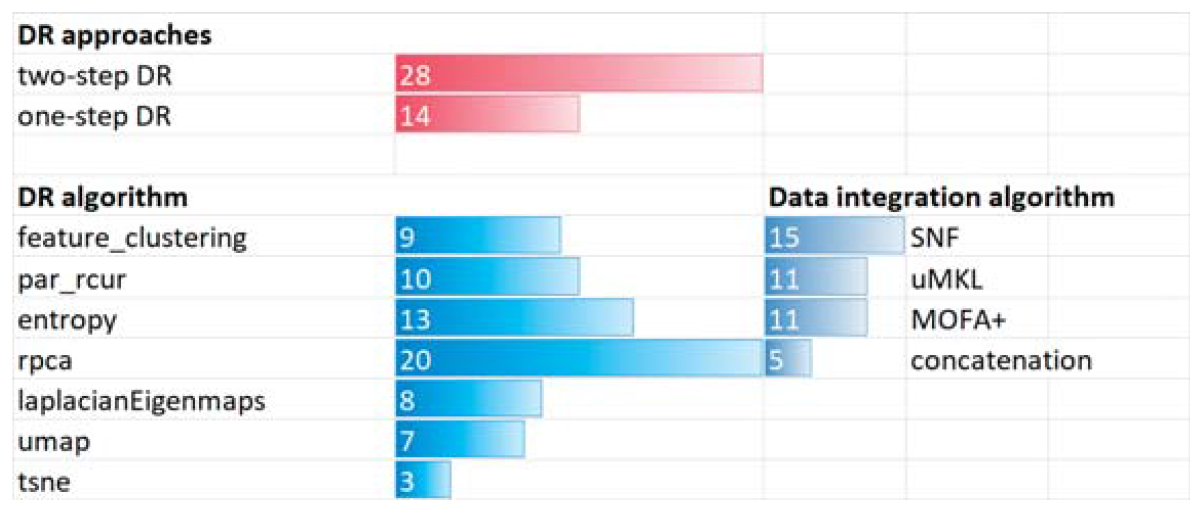
Frequencies of DR algorithms, DR approaches (one-step versus two-step), and data integration methods that appear among the best models when block-analysis guides the DR of the four omics composing the multimodal dataset.

#### 4.3.3. The usage of all the available omics improves robustness with respect to noise and data unbalance

While performing the experiments we reasoned that the usage of all four omics might provide redundant and/or misleading information for the problem at hand. Moreover, some data-fusion algorithms might profit when fewer views are integrated. Therefore, we ran experiments to compare the usage of all the available (four) omics to the usage of multi-omics datasets containing at least two omics. Win-tie-loss tables comparing results obtained when using multi-omics combinations, DR pipelines, and data-fusion algorithms are shown in figures 7 (when the AUC is used for evaluation) and figure 8 (when the AUCPR is used for evaluation). Extracts of win-tie-loss tables comparing the whole DR+data-fusion pipelines (also specified by the input multi-omics combination) are shown in supplementary figures S. F.15 and S. F.16, while supplementary files S10 and S11 report the complete win-tie-loss tables.

**Figure 7:**
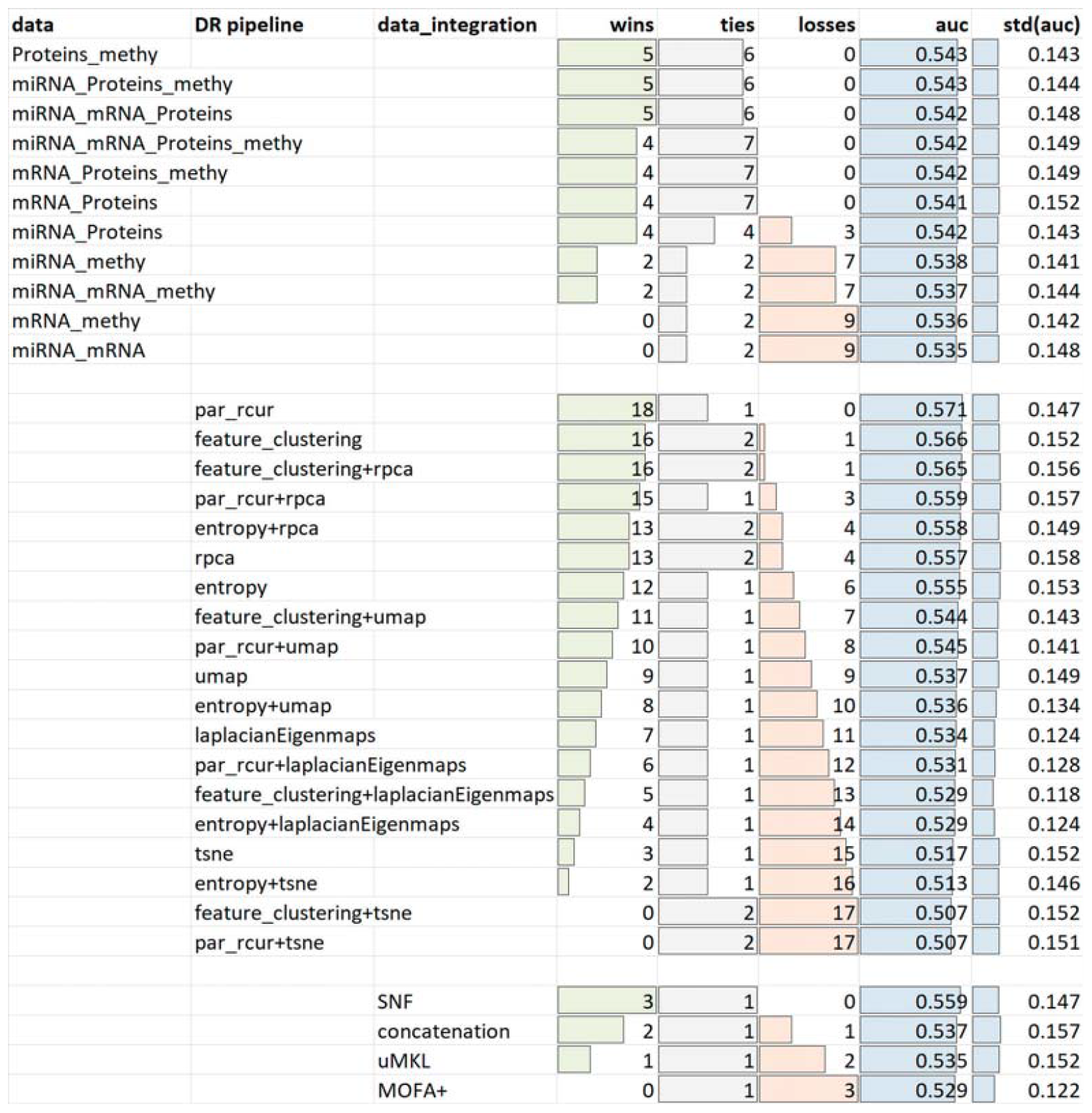
Win-tie-loss tables computed by Wilcoxon signed-rank test when using the AUC measure to compare models guided by block-analysis, integrating at least two omics, and neglecting demographic data.

**Figure 8:**
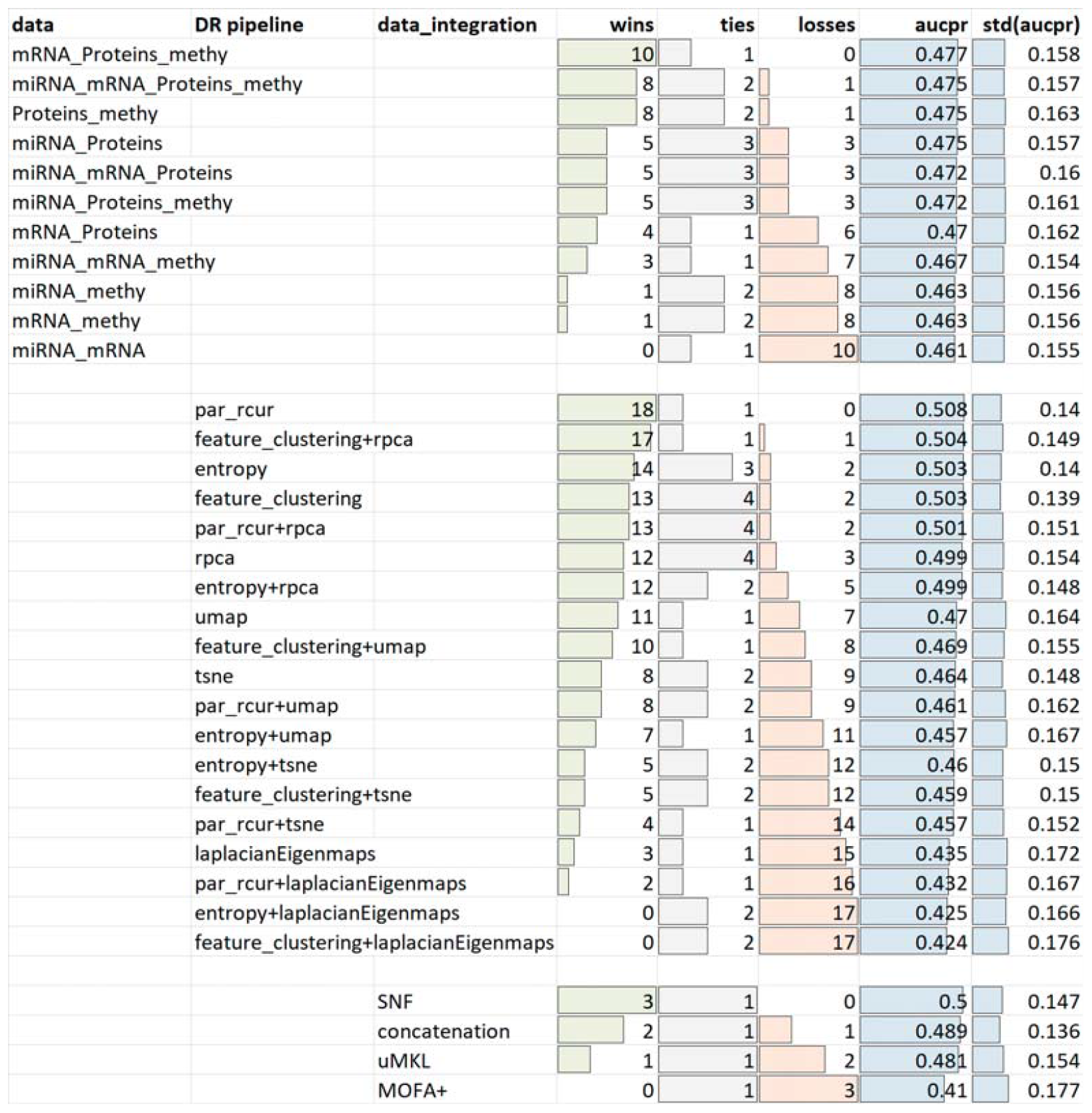
Win-tie-loss tables computed by Wilcoxon signed-rank test when using the AUCPR measure to compare models guided by block-analysis, integrating at least two omics, and neglecting demographic data.

Regarding the comparison of the input-views (top tables in figures 7 and 8) win-tie-loss tables obtained with AUC seem suggesting that specific combinations of (mainly) three omics achieve results that are comparable and slightly better than those obtained when using four omics. However, when observing the win-tie-loss table obtained by using the AUCPR we note that the usage of four omics is comparable to only one combination of three omics (which, again, scores slightly better) and one combination of two omics. Considering that most of the datasets used for our tests are unbalanced, i.e. the AUCPR measure is less biased and more informative while the AUC might be over-optimistic [43], we can infer that the usage of a superior number of omics, i.e. a superior number of features potentially adding more informative content but also some noise and redundancy, does not affect performance but instead achieves results comparable to specific combinations of omics and guarantees robustness with respect to data unbalance. Note that, this performance is indirectly related to the effectiveness of the prior DR step. Indeed, we recall that, when no DR is applied at all (supplementary section S. F.1), the algorithms that fuse all the omics and then apply supervised feature selection and hyper-parameter tuning achieve poor results (supplementary section S. F.1). Therefore, provided that a proper DR is applied, the usage of all the available omics allows to obtain robust results and to avoid costly experiments to choose the most suitable combination of omics given the problem at hand.

Among the DR pipelines, we again note that feature clustering followed by RPCA, the iterative RCUR (either alone or in combination with RPCA), or RPCA alone were still the most robust DR approaches. When instead the paired-samples Wilcoxon test was used to compare the four data-fusion algorithms, SNF confirmed its superiority for both AUC and AUCPR measures. On the other hand, the simple integration via concatenation seemed to benefit from the reduction of omics, which is probably due to the lower number of features when less than four omics are considered.

To provide an exhaustive description of the obtained results, for each dataset we collected the list of experiments that had the highest AUC or AUCPR. Figure 9 plots the frequency of appearance of each multi-omics combination, DR pipeline, and data-fusion algorithm (supplementary table S12 lists all the best models and their performance). For what regards the composition of the input multi-omics combinations, all views but the miRNA-view equally contributed in obtaining a good performance. Moreover, while less robust when compared to combinations of three/four omics by Wilcoxon signed-rank tests, also combinations of two omics could appear among the top-performers; this suggests that, when having enough samples and computing power, the combination of input views could be regarded as a further hyper-parameter to be tuned to optimize performance.

**Figure 9:**
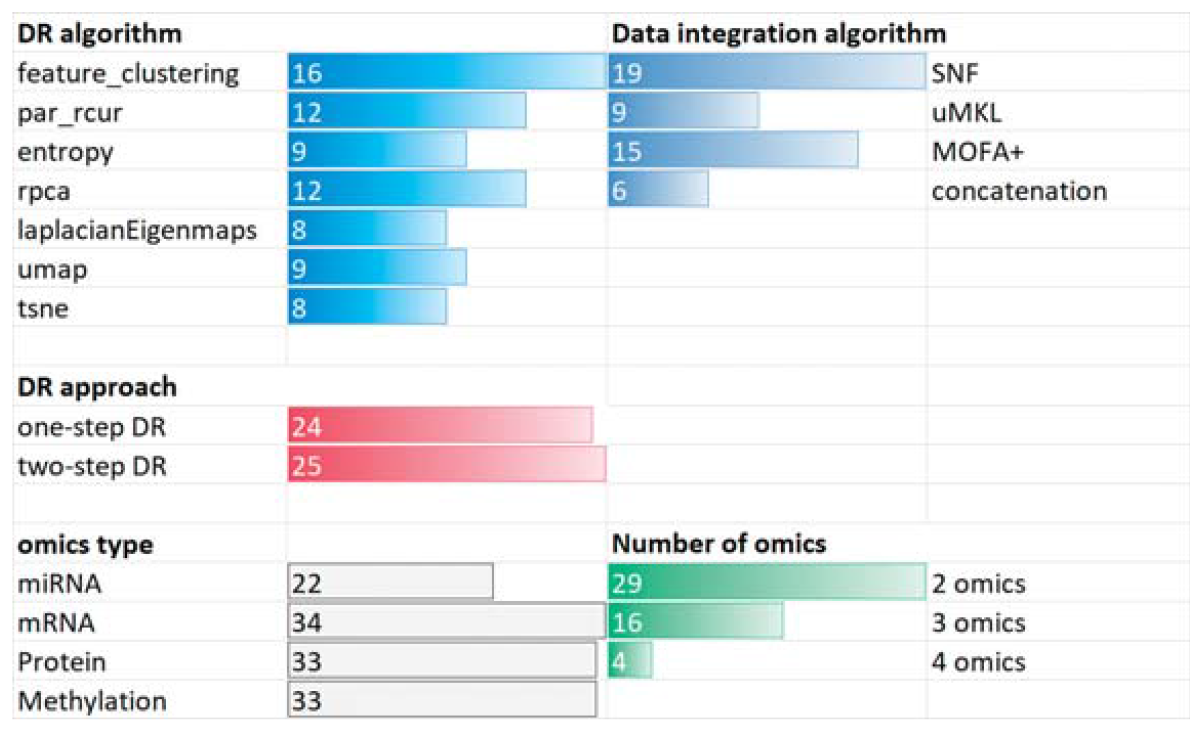
Frequencies of multi-omics combinations, DR pipelines, and data-fusion methods appearing among the best models when at least two omics are fused and the block-analysis guides the DR.

#### 4.3.4. Integration of demographic patient data may further improve results

The available TCGA datasets contain demographic descriptions (gender, age at diagnosis, ethnicity, race) that might provide further useful information to improve the performance of the supervised classification task.

Since several bio-medical studies provide both omics and non-omics patients’ descriptors, and considering the documented literature interest about challenges regarding the integration of omics and non-omics views [44] (supplementary section S. F.3.1), it was interesting to understand not only if patients’ descriptors other than multi-omics may improve results of our supervised analysis, but also if a simple approach that concatenates patients’ descriptors to the fused multi-omics view could be more effective than using the patients’ descriptors as a further view to be integrated.

In our classification pipeline, once the multi-omics views are integrated, concatenation of demographic descriptors is possible because RF can process heterogeneous data. On the other hand, to use SNF, uMKL for integrating multi-omics and demographic views we used the Gower similarity^8^ to compute pairwise similarities, and then used them as the fifth kernel to be integrated by SNF and uMKL. When using MOFA+ we simply provided the demographic view as a further view to guide the discovery of the latent components.

Based on the results from the previous experiment (section 4.3.3), in this comparative evaluation we limited the number of experiments to those that used all the available omics.

Figures 10 and 11 report the win-tie-loss tables computed by Wilcoxon signedrank test when comparing the integration of omics and non-omics descriptors to the usage of only omics-views. Supplementary figures S. F.17 and S. F.18 show the win-tie-loss tables obtained when comparing full pipelines characterized by a specific set of views being integrated, and a specific DR+data-fusion pipeline (complete tables can be found in supplementary tables S13 and S14). In the figures, “4 omics+ pt” refers to the usage of omics and non-omics variables; “MOFA+ + PT data”, “SNF + PT data”, and “uMKL + PT data” refer to the data-fusion algorithms also integrating the demographic view; “MOFA+”, “SNF”, “uMKL”, and “concatenation” refer to the traditional application of the data-fusion algorithms for integrating multi-omics, followed by concatenation with the demographics views.

**Figure 10:**
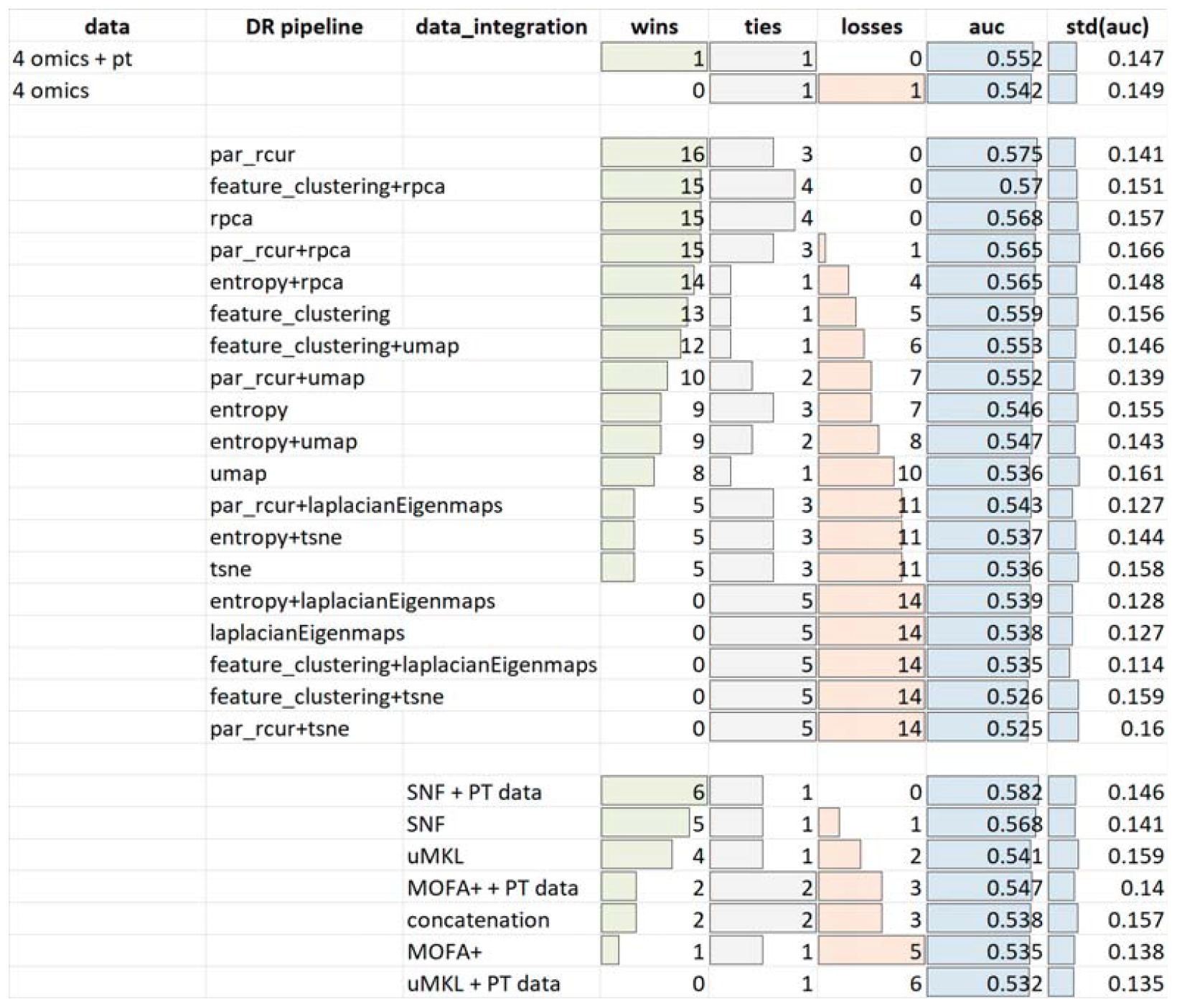
Top table: Win-tie-loss tables obtained when assessing the integration of omics and non-omics descriptors (“4 omics+pt”) by comparing it to the usage of only the omics descriptors (“4 omics”). Center table: DR pipelines are compared when 4 omics and demographic descriptors are integrated. Bottom table: data-fusion algorithms are compared when 4 omics and demographic descriptors are integrated either by concatenating demographic descriptors to the integrated representation (“SNF”, “uMKL”, “MOFA+”, and “concatenation”), or by using the non-omics descriptors as a further view to be integrated (“SNF + PT data”, “uMKL + PT data”, “MOFA+ + PT data”). AUC is used when computing the Wilcoxon signed-rank test.

**Figure 11:**
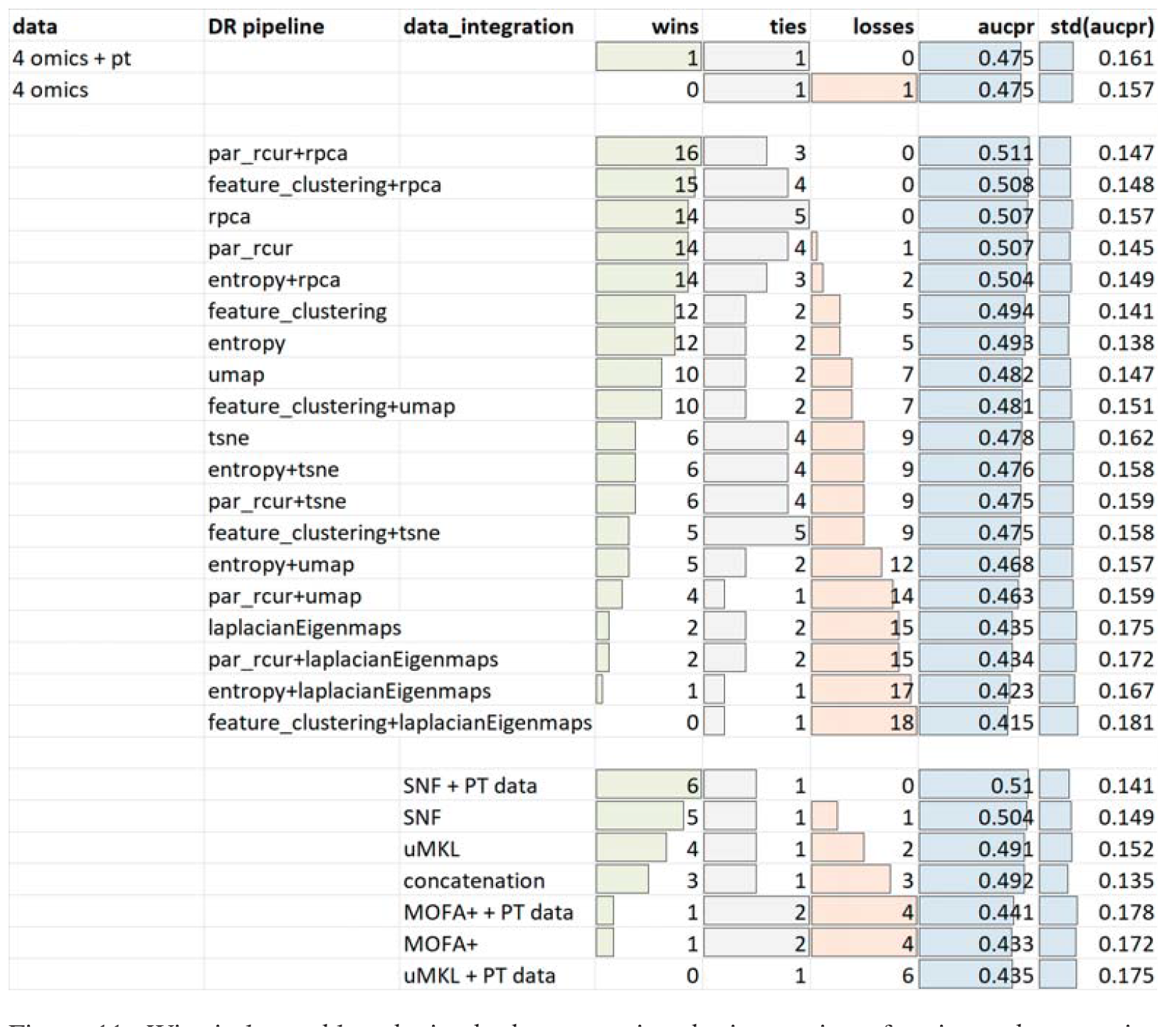
Win-tie-loss tables obtained when assessing the integration of omics and non-omics descriptors (“4 omics+pt”) by comparing it to the usage of only the omics descriptors (“4 omics”). AUCPR is used when computing the Wilcoxon signed-rank test.

Figure 12 plots the frequency of each data view, DR algorithm, and data-fusion method appearing in the top-performing experiments (listed in supplementary table S15).

**Figure 12:**
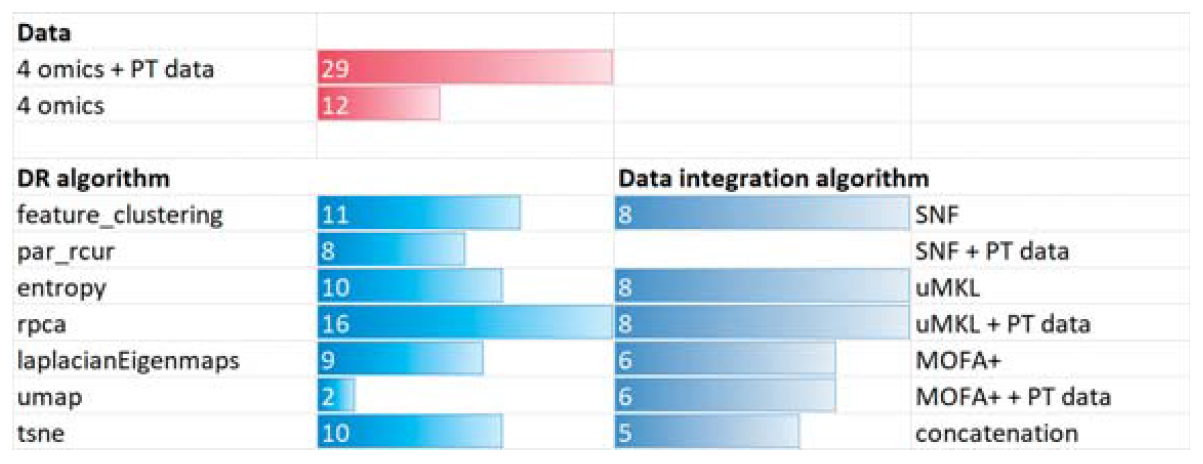
Frequencies of DR methods, and data integration algorithms appearing among the top-performing models when all the views (omics and non-omics) are analyzed.

Observing the results we understand that surely the inclusion of patient data improves performance; indeed, for both AUC and AUCPR, combinations of views including patient data always win with respect to combinations neglecting demographic predictors. The comparative performance of DR pipelines remains unaltered with respect to previous experiments; indeed, iterative RCUR (par rcur) and feature clustering followed by RPCA, or the usage of a unique step of par rcur or RPCA are the DR methods achieving the most robust results for both AUC and AUCPR.

Regarding the data integration models, SNF is still among the top-performing models. When comparing the two (multi-omics plus demographic) integration approaches, we note that both “SNF + PT data” and “MOFA + PT data” (using the demographic descriptors as a further view to be integrated) score better than their counterparts (“SNF” and “MOFA+”) that simply concatenate the demographic variables to the integrated multi-omics views. This is not true for uMKL, which, according to the win-tie-loss tables, obtained more robust results when we first integrated omics descriptors and then concatenated demographic variables to the fused kernel representation. This might be due to the fact that the Gower similarity measure is not a proper kernel similarity, as required by uMKL. Despite this fact, we note that “uMKL” and “uMKL + PT data” appear with the highest frequency in the list of top-performing models (supplementary table S15 and figure 12), which evidence the potentials of the uMKL data-fusion strategy and suggests that the transformation of Gower similarity into a kernel matrix might further improve results.

Overall, these results suggest that integration of omics and non-omics variables, when opportunely designed, might be a promising way. Indeed, When considering the more detailed comparison between DR+data-fusion pipelines (extracts of the top twenty-five winners in figures S. F.17 and S. F.18) we note that the best fusion algorithm is MOFA+ integrating demographic descriptors. Considering that one of the advantages of MOFA+ relies on its ability to integrate heterogeneous data type, its superiority is not a surprise and further supports our belief that the integration of omic and non-omics data is a promising way that needs a careful design.

## 5. Discussion and Conclusions

In this paper, we have described a novel application of block-analysis to leverage any of the most promising `id` estimators and obtain an unbiased `id`-estimate of the views in a multi-modal dataset. We also proposed an automatic analysis of the block-`id` distribution computed by the block-analysis to detect feature noise and redundancy contributing to the curse of dimensionality and therefore evidence the need to apply a view-specific dimensionality reduction phase (guided by the `id` estimate) prior to any subsequent analysis to reduce curse-of-dimensionality effects.

Using the proposed `id` analysis we can therefore automatically take view-specific decisions so that views containing higher levels of noise and redundancy can undergo a two-step dimensionality reduction approach combining the advantages of feature-selection and feature-extraction; on the other hand, views less affected by the curse of dimensionality may be reduced by traditional feature-extraction approaches.

Besides assessing our proposal by using nine heterogeneous multi-omics cancer datasets from the TCGA repository, we analyzed and compared the DR effects on the subsequent application of data-fusion techniques that have shown their promise in the field of multi-omics. To this aim, we used the fused view to predict overall survival events by means of RF classifiers, often preferred in the biomedical field due to their interpretable nature, relative robustness to hyper-parameter settings, superior efficiency, and effectiveness in many competitions [40].

The results we obtained first evidenced that a properly designed DR step is crucial and should never be neglected when complex multi-omics data is processed. Secondly, we showed that DR approaches guided by block-analysis were superior to traditional DR approaches setting the dimension of the reduced space by some heuristics or by preliminary empirical experiments.

When analyzing the impact of the proposed DR approach on different multiomics fusion settings we first noticed that the two-step DR approach can be an effective solution. Particularly, in our classification task the most robust and effective results were achieved when combining the iterative version of RCUR we implemented (or feature clustering) with RPCA, whose formulation is simpler and more explainable than, e.g., UMAP and t-SNE.

When observing the robustness and performance of the experimented datafusion algorithm we confirmed the efficiency and effectiveness of SNF, which showed its robustness with respect to different settings. MOFA+ also showed its promise, though its robustness and efficiency were lower than that of SNF. However, the advantage of MOFA+ relies on its ability to deal with heterogeneous data types, so that it can effectively integrate omics and non-omics views.

Regarding two different and crucial data-fusion settings we were interested in investigating, we noted that comparable results can be obtained when using all four omics or specific subsets of (at least) two omics. This suggests that, in our experiments, the DR and data-fusion steps are able to cope with the presence of potentially redundant information within and between the four omics views, so that all the available omics types can be used without the need to try all their different combinations. In other words, the design of a proper DR can facilitate the following data-fusion task by effectively removing redundancies within individual data types, while better exposing their characterizing informative content. This facilitates the task of the following data-fusion algorithms, which must only deal with redundancies across views while uncovering the shared and individual informative content of the multiple omics. This allows to avoid costly empirical experiments to choose the subset of omics to be integrated.

We further assessed the addition of patients’ demographic descriptors in the analysis and showed that it effectively increases the classification performance.

Concluding, all the experiments led us to the definition of a DR+data-fusion pipeline that obtains promising results without the need for empirical and heuristically based choices. In particular, considering that all the results reported in our experiments (besides being averaged across all the nine TCGA datasets) were obtained when applying neither supervised feature selection nor hyper-parameter tuning to avoid confounding effects that would bias the comparative assessment, we ran the last experiment where we used all the four omics as input to data fusion, we concatenated the fused representation with demographic descriptors and then optimized the RF performance by supervised feature selection through RF importance [39] and hyper-parameter tuning through internal stratified holdout validation (more details are reported in supplementary section S. F.4).

This procedure allowed obtaining more than satisfactory results (see figures 13 and 14), with AUCs often greater than 0.70*/*0.75 and large AUCPRs even for datasets characterized by a low (≤ 0.2) ratio between the number of positive cases and the total number of cases. Of note, these results greatly outperform the results we obtained when we avoided any DR step prior to data-fusion, eventual concatenation of demographic descriptors, and RF optimization via supervised feature selection and hyper-parameter tuning (supplementary figure S. F.6 and figure 15).

**Figure 13:**
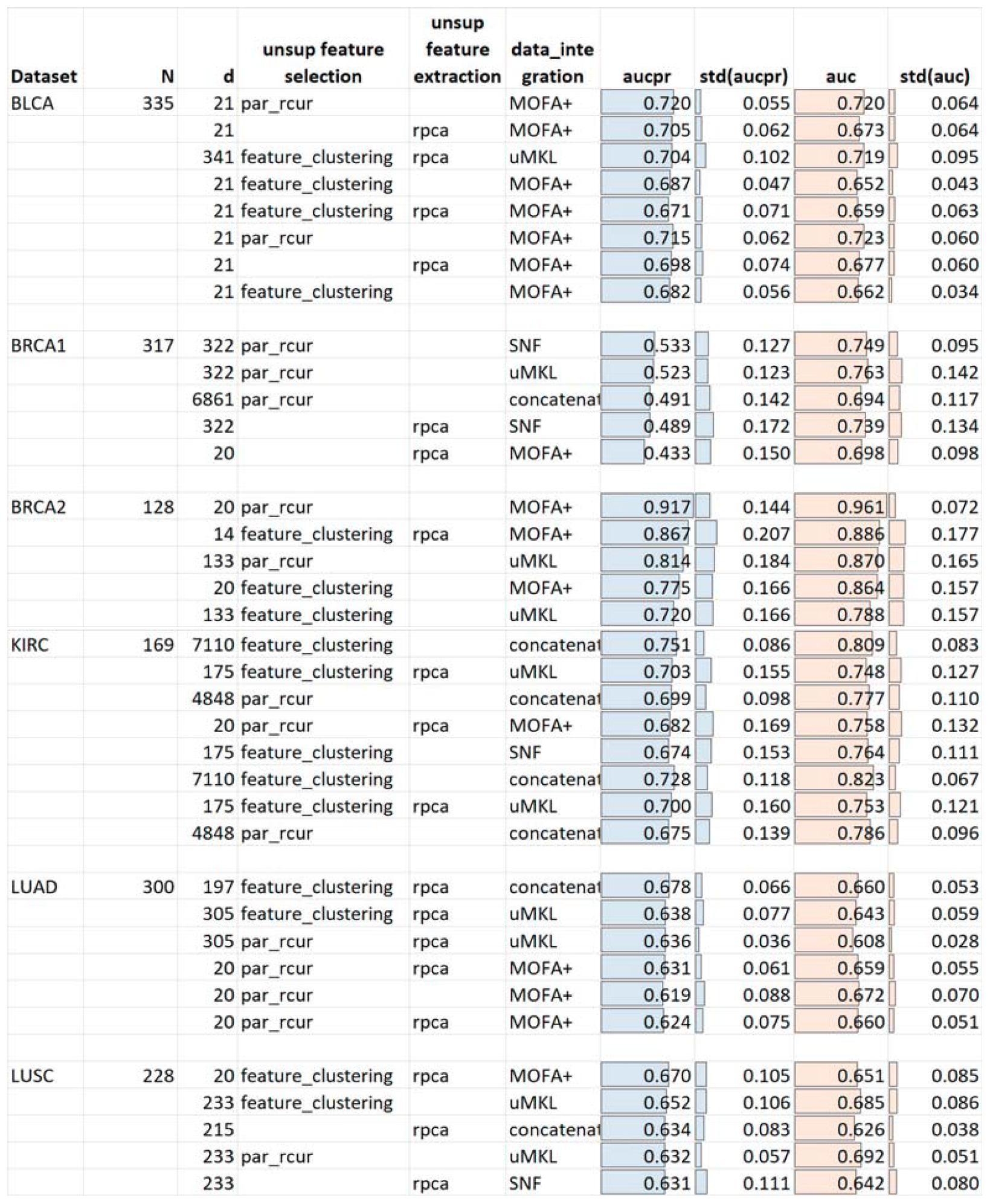
Classification performance obtained one the BLCA, BRCA1, BRCA2, KIRC, LUAD, and LUSC datasets when using the block-analysis to guide the DR of all four omics. After reduced data-fusion the demographics views are concatenated and used as input for supervised feature selection, RF tuning, RF training, and sample classification.

**Figure 14:**
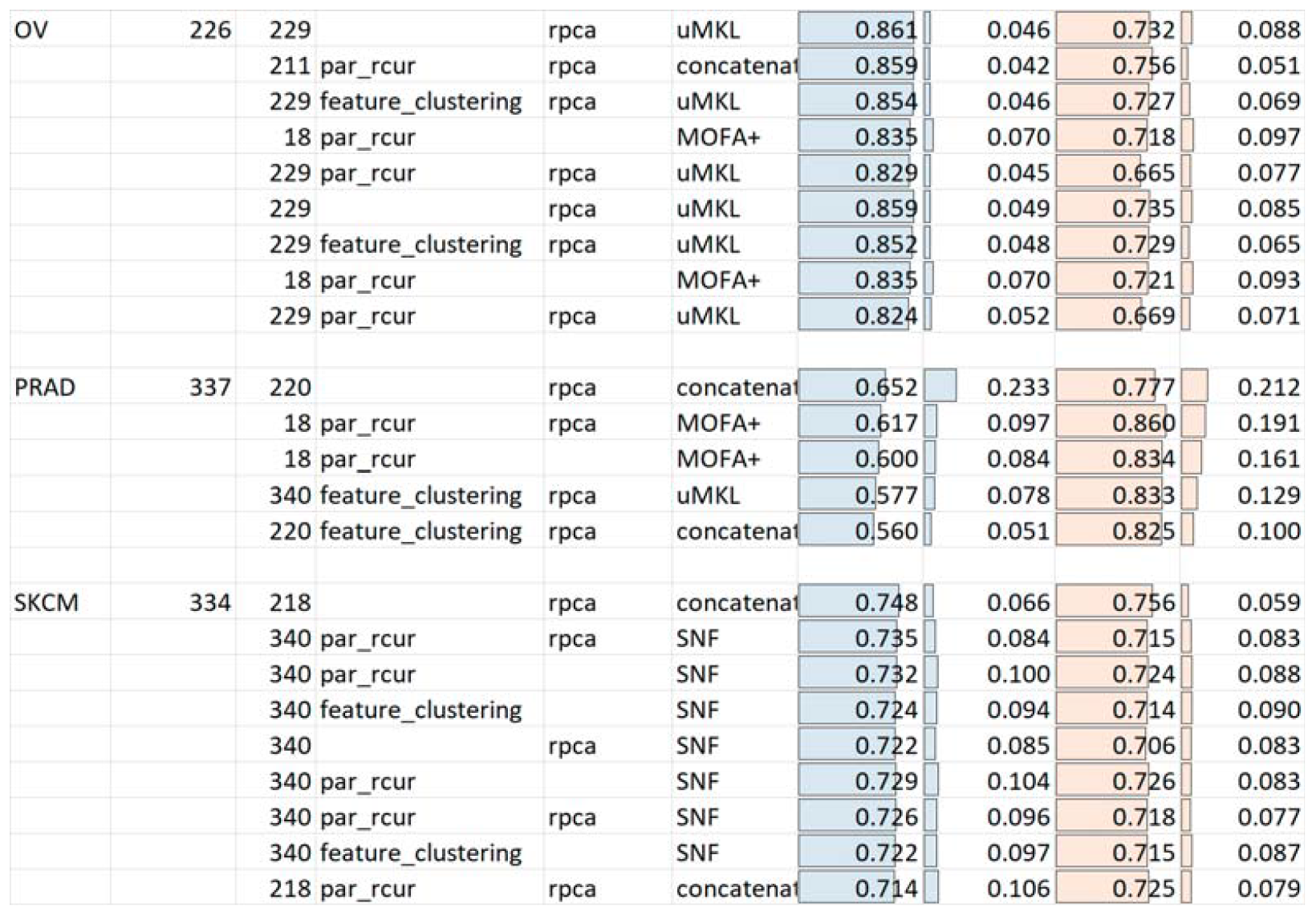
Classification performance obtained on the OV, PRAD, and SKCM datasets when using the block-analysis to guide the DR of all four omics. After reduced data-fusion the demographics views are concatenated and used as input for supervised feature selection, RF tuning, RF training, and sample classification.

**Figure 15:**
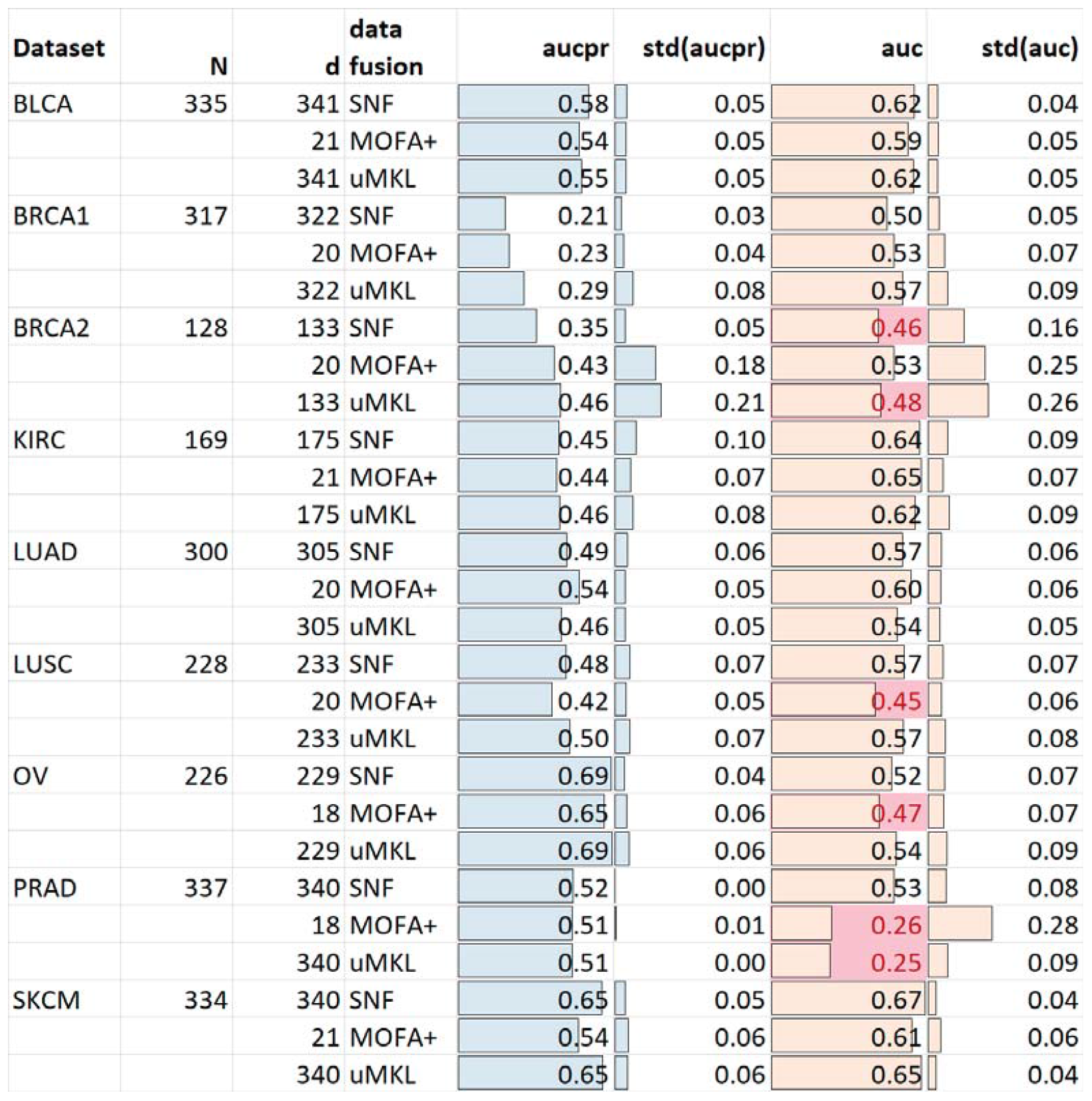
Classification performance obtained when integrating all four omics and then concatenating the demographics views. No prior DR is performed but supervised feature selection and hyper-parameter tuning are applied prior to RF training.

## Supporting information

Supplementary images

## Acknowledgements

Authors would like to thank Prof. Juho Rouso (lead of KEPACO Lab - Department of Computer Science, Aalto University, Espoo, Finland) for his invaluable support, and PhD Riikka Huusari (Department of Computer Science, Aalto University, Espoo, Finland) for her precious comments.

## Footnotes

1 The R package “curatedTCGAData” [24] was used to download the tumor datasets from the TCGA repository (dataset version 2.0.1).

2 In all our experiments we set *M* = 11 and *t* = 90%. The low value of *M* limits the computational-time costs of the algorithm; however, the higher this value, the lower the variability of the estimate and the higher the precision of the estimate. The value of *t* = 90% is chosen to obtain under-sampled datasets with enough samples.

3 We set *n*_*try*_ = 31 to reduce time costs of the algorithm; however, the higher this value, the higher the precision of the estimate.

4 Note that most feature-extraction techniques are based on the computation of pairwise sample distances, which are biased under the curse of dimensionality due to the high level of sample-sparsity. Besides the reduction of computational costs, the prior application of a feature-selection algorithm reducing the amount of redundancy and noise facilitates the task of the following feature-extraction algorithm by allowing the computation of more reliable pairwise sample-distances.

5 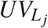 is computed as the mean of the block-id variances for blocks 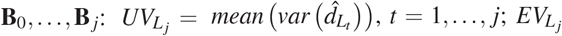 is computed as the variance for blocks 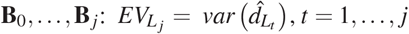

6 If the input-data contains discriminative information and a limited amount of redundancy, default RF parameters can achieve decent results. Moreover, while it is undoubted that supervised feature selection and hyper-parameter tuning increase RF performance, it is also true that the increase also depends on some randomness; by avoiding preliminary feature selection and hyper-parameter tuning, we could more directly focus on comparing the performance of the DR+data-fusion pipelines, which are not confounded by the effects of feature selection and hyper-parameter tuning choices.

7 When using sided-hypothesis tests to compare the performance of two methods *A* and *B*, a win/loss (or tie) is assigned if the sided-test is below (above) the α-value. When assessing multiple methods, all pairwise comparisons are performed, and a three-column table is computed that lists, for each method, the number of wins, ties, and losses

8 Gower distance/similarity is a measure of dissimilarity or similarity between two individuals (or data points) described by a set of heterogeneous variables, including categorical, binary, or-dinal, and numerical variables. The Gower distance/similarity is computed as the average of all the distances/similarities measured on each variable, taking into account the data types of those variables.

