## Supplementary images for "Intrinsic-Dimension analysis for guiding dimensionality reduction and data-fusion in multi-omics data processing"

### Supplementary material

#### S. A. Background

In this section we review DR approaches that are among the most used in literature (supplementary section S. A.1), and we group them into feature selection (supplementary section S. A.1.1) and feature extraction techniques (supplementary section S. A.1.2). Then, we report the state-of-the-art of `id` estimation (supplementary section S. A.2) and we detail one of the most promising `id` estimators, namely *two-nn* (supplementary section S. A.2.1), which we leverage to obtain unbiased estimates by block-analysis; finally, we briefly recall successful state-of-the-art data-fusion algorithms (supplementary section S. A.3).

##### *S. A.1. Dimensionality Reduction*

DR techniques can be grouped in two main classes: *Feature Selection* algorithms select an "informative" subset from the original feature-set; *Feature Extraction* techniques analyze the input feature set to extract a reduced feature-basis that allows coding the sample set in a salient subspace maintaining most of the original information.

###### *S. A.1.1. unsupervised Feature selection*

Feature selection algorithms [45] may be either *supervised* or *unsupervised*. Supervised feature selection algorithms are applied prior to (or are embedded into) a supervised classification task. They select features that best discriminate between the available classes; these methods can be split into algorithms that are independent of the following classification model, e.g. Minimum Redundancy Maximum Relevance (MRMR, [46]), or depend on the following model, e.g. recursive feature elimination (RFE, [47]) and permutation importance selection algorithms (e.g. Boruta [48]). Supervised feature selection algorithms often improve performance when the data sample has enough cases compared to the data dimensionality; however, under the curse-of-dimensionality phenomenon, they are computationally intensive, often impracticable and, above all, they often select a feature subset that can easily bring to overfitting with poor generalization (this is also shown by our preliminary experiments, reported in supplementary section S. F.1). Therefore, even in a supervised classification task, a first unsupervised reduction is often needed, after which supervised feature selection algorithms may be successfully applied.

*Unsupervised* feature selection algorithms, sometimes called “filter methods”, [49, 50] do not make use of data-labels but instead remove noisy, redundant, and not-informative features to obtain a reduced, salient, and representative feature subset. Feature selection methods in this class contain a wide range of algorithms, spanning computationally cheap algorithms that are therefore practical also in high dimensions, such entropy filtering or CUR/RCUR decomposition [36, 51, 52], to computationally expensive algorithms that do not scale in high dimensions, such as feature-clustering methods [53] (where the features that are cluster centroids are selected), or Spectral Feature Selection (SPEC [54]). It has to be noted that, though often unpractical, feature clustering and spectral feature selection approaches have obtained the most promising results in literature [49, 50]. However, while feature clustering can be redesigned to reduce the computational costs by parallel implementations that obtain practical computational times (a sketch of our parallel implementation is reported in supplementary section S. D.1), spectral clustering cannot be distributed. Therefore, in our tests we experimented feature clustering (supplementary section S. D.1), entropy filtering, and an iterative version of RCUR (supplementary section S. E)

*S. A.1.2. unsupervised Feature extraction*

*Feature extraction* techniques assume that the samples in each high-dimensional view of the multi-modal dataset are drawn from an originally lower-dimensional manifold that has been twisted and curved by a smooth mapping, so that the observed high-dimensional space contains the original information but also a good amount of redundancy and noise. Hence, the true information can be outlined by many fewer coordinates than those of the high-dimensional space. To recover the original low-dimensional data state-of-the-art feature extraction methods transform/embed the original high-dimensional data (manifold) by either applying factor-analysis techniques or manifold-embedding approaches.

In particular, factor analysis techniques linearly/non-linearly combine the original features to project the data into a reduced space where redundancies are removed while salient information is preserved and eventually emphasized [55, 51]. In the multi-omics data analysis field, the mostly used factor analysis techniques are the classic PCA algorithm, mainly used for exploratory data analysis [16, 17], and its randomized variation (randomized SVD [56, 55, 18]).

On the other hand, manifold-embedding approaches solve optimization problems that find the lower-dimensional manifold maximally preserving the local and global topological structure of the dataset. Examples of manifold-embedding methods are the Laplacian Eigenmaps (LE, [42]) algorithm, which is the first and effective DR step of the spectral clustering algorithm used by many authors to identify novel patients' subtypes [57], or t-distributed stochastic embedding (t-SNE, [58]) and Uniform Manifold Approximation and Projection (UMAP, [59]), which are mainly used for high-dimensional data-visualization purposes and, due to their promising results and solid theoretical base, have been also successfully experimented for RNA sequencing data reduction [12]. In particular, the comparison reported in [12] has highlighted their superior performance with respect to classic linear models (e.g., PCA), and to more complex methods, such as deep variational autoencoder models (VAE, [60]), and a multiple-kernel-learning technique (SIMLR [61]) for a simultaneous reduction and integration. The experimental comparison shows the instability of VAE and SIMLR on biomedical sets highly affected by the small-sample-size.

Computational complexity and instability in high-dimension are two problems often affecting feature extraction models. While computational complexity is due to the large search space where the best (reduced) solution must be found, instability is due to the fact that most feature extraction algorithms base their analysis on pairwise-point relationships, which are biased when incurring the small-sample-size problem and the curse of dimensionality. Some algorithms, e.g, UMAP and t-SNE, cope with both problems by applying a preliminary DR step, where PCA or other linear models are applied to produce a lower-dimensional data representation that is more tractable and less affected by the curse of dimensionality. In this work we similarly propose a sequential combination of a feature selection algorithm, which provides a first reduction by selecting representative features, and a feature extraction algorithm, which combines the representative features to obtain the reduced dataset (see Section 4.1).

#### *S. A.2. ID estimation*

The dimension of the lower-dimensional space where to project the reduced data is a value that should be carefully chosen [17]; while excessively large values would bring to the computation of still sparse, noisy, and redundant datasets, excessively low values would cause the loss of salient information. Such a choice is still an open problem. In this paper we propose guiding the DR step by the `id` of the dataset [62, 32]. The `id` is the minimum number of dimensions of a lower dimensional space where the data can be projected (by a smooth mapping) in order to minimize the information loss and maintain its characterizing structure.

Unfortunately, the estimation of the topological dimension of a manifold using a limited set of points, (that are assumed to be) uniformly drawn from it, is a challenging and not yet solved task. All the state-of-the-art `id` estimation techniques exploit differing underlying theories, according to which they are often grouped into the following four main categories [32]: *Projective `id` estimators*, *Topological `id` estimators*, *Fractal `id` estimators*, and *Nearest-Neighbors (NN) based `id` estimators*.

*Projective `id` estimators* [63, 64, 65] basically process a dataset to identify a somehow appealing lower dimensional subspace, e.g. the one minimizing the information loss, where to project the data and whose vector space dimension is viewed as the `id` estimate.

*Topological `id` estimators* [66, 67], assume the $D$-dimensional samples in the available dataset $X_n = \{\mathbf{x}_i\}_{i=1}^n \subset \mathbb{R}^D$ have been uniformly drawn from a manifold $\mathcal{M}$ with topological dimension equal to $d$ that has been embedded in a higher $D$-dimensional space through a nonlinear smooth mapping. Under this assumption, the `id` is estimated as the manifold's topological dimension, defined through Lebesgue's Covering Dimension [68].

Since topological dimension is very difficult to be practically estimated, several authors implicitly assume that the original manifold $\mathcal{M}$ has a somehow fractal structure [69] and estimate the `id` by employing *fractal dimension estimators*. Roughly speaking, since the basis concept of all fractal dimensions is that the volume of a $d$-dimensional ball of radius $r$ scales with its size as $r^d$ [69, 70], all state-of-the-art fractal `id` estimators [71, 72] are based on the idea of counting the number of observations in a neighborhood of radius $r$ to (somehow) estimate the rate of growth of this number. If the estimated growth is $r^d$, then the estimated fractal dimension of the data is considered to be equal to $d$.

*NN-based `id` estimators* [31, 22] describe data-neighborhoods' distributions as a function of the manifold topological dimension $d$, and then estimate $d$ by analyzing the $k$-nearest neighborhoods in the available datasets, assuming they approximate small $d$-dimensional hyperspheres with radius $r \to 0 \in \Re^+$.

In the bioinformatics field, the available datasets are often noisy and complex. In this context NN estimators often outperform Projective, Topological and Fractal `id` estimators; indeed, projective and topological-based `id` estimators produce reliable estimates for data drawn from manifolds with mainly low curvature and low `id` values, while fractal `id` estimators fail when the points are noisy and/or not uniformly drawn from the underlying manifold. On the other hand, NN estimators such as DANCo [31] have shown their robustness on not-uniformly drawn, noisy and complex datasets, where the two main assumptions at the base of fractal, topological, and projective `id` estimators are often violated. Indeed, (1) the points cannot be assumed to be uniformly drawn from the manifold where they are assumed to lie, and (2) the complexity of the available datasets allows assuming that the points lie on more-than-one, eventually intersecting manifolds, each characterized by a specific topological dimension.

To account for the aforementioned issues, NN estimators often compute a reliable “global” `id` estimator by integrating all the “local” IDs estimated over point-neighborhoods. In a recent work, authors performed preliminary experiments to compare **DANCo** [31] and ***two-nn*** ([22] , see supplementary section S. A.2.1), two recent state-of-the-art NN-based `id` estimators. The reported preliminary results confirmed the results also reported in [22]; while DANCo is more robust with respect to noisy, real datasets, often containing outliers, *two-nn* has a much lower computational cost and no crucial parameters to be chosen. Besides, the usage of the block-analysis we describe in section 3.1 allows to reduce the biases affecting real datasets, improving the robustness of *two-nn* to noise and outliers and obtaining estimates comparable to DANCo. For this reason, in this paper we use the *two-nn* estimator to guide the DR of each view.

*S. A.2.1. The two-nn `id` estimator*

*two-nn* [22] is a NN `id` estimator that has been developed by considering that several state-of-the-art `id` estimators compute estimates that are either influenced by the cardinality of the considered point neighborhoods, or are not robust with respect to not-uniformly distributed point-neighborhoods, which is often the case of real-world bioinformatics datasets. To tackle the aforementioned problems, the authors proposed theories that highlight how, assuming local uniformity across 2-NN neighborhoods, the volume of the shell between the first and the second NNs of each point in a manifold is dependent on the (local) manifold ID. This ultimately brings to a Pareto law relationship linking the (true) manifold ID, $d$, and the ratio, $\mu_i = \frac{r_{2i}}{r_{1i}}$ of the distances between the $i-$th point and its second and the first NN: $\mathcal{L}(\mu_i; d) = d\mu_i^{(d-1)}$. Based on this relationship an `id` estimate, $\hat{d}$, can be derived by fitting the empirical cumulative distribution of the $\mu_i$s through a maximum likelihood estimator.

The *two-nn* estimator reaches an estimation accuracy comparable to the state-of-the-art estimator DANCo Note that the *two-nn* formulation requires assuming a uniform distribution only across 2-NN neighborhoods, and, compared to the state-of-the-art DANCo estimator, should therefore be less affected by twisted and curved datasets. Indee Moreover, when testing DANCo against *two-nn* [73] we noted DANCo has a much higher computational complexity, it is strongly influenced by the neighborhood size parameter, and has a greater variance of the computed estimates. On the other hand, authors of *two-nn* showed the superior performance of DANCo in the presence of boundary points [22], often characterizing datasets where the imbalance between dimension and sample cardinality is extremely large. To cope with this weakness *two-nn* authors propose computing an unbiased estimate by a block-analysis technique. Considering the appealing properties of *two-nn* in this work we preferred it to DANCo and we used the iterative estimation scheme proposed by the authors to cope with boundary points. More precisely, given an omics view, we randomly undersampled it $M = 11$ times to keep the $t = 90\%$ of cases, we computed the *two-nn* estimate of all the undersampled sets, and we averaged them to obtain a estimate robust with respect to noisy and outlier points.

#### *S. A.3. Multi-omics data integration methods*

Patient similarity Networks (PSNs) are emerging as a new approach for analyzing patient data to develop predictive approaches. Beside the advantages of using PSNs in the bio-medical field, Gliozzo et al. [11] survey multi-omics data-integration techniques that allow to build or fuse PSNs. In their survey, data-fusion techniques are split into three categories, *PSN-fusion*, *input-data fusion* models, and *output-data fusion* algorithms.

Among *PSN-fusion* techniques, the first literature attempts proposed using supervised multiple kernel learning algorithms (MKL, [74]), which integrate multiple kernels by leveraging support-vector-machine classifiers; though promising, these methods often become problematic when limited sample size is available, due to the need of performing an internal validation to tune crucial hyper-parameters. A limited validation-set size often leads to poor generalization and impacts the classifier reliability. Moreover, the most promising supervised MKL algorithms work on numeric data types and only a few attempts [75] have been proposed to integrate heterogeneous views.

To cope with the limited sample size, even in the context of supervised classification, most authors prefer to apply an unsupervised integration step, followed by a supervised classification. Under this rationale, in [23] the authors propose a successful unsupervised MKL (hereafter named uMKL) algorithm finding a consensus kernel that is able to deal with kernels computed on heterogeneous types.

A completely different PSN-fusion approach is used by the well-known Similarity Network Fusion (SNF, [13]) algorithm and by all its variants. SNF builds similarity kernels encoding the global and local patient-similarity network information, and applies an iterative diffusion process that performs a weighted average of the multi-view similarities between two patients (nodes in the similarity networks) if they have "shared" neighborhoods in all the views. SNF and its variants are probably the most used approaches in the context of multi-omics data integration, where they have been always applied for unsupervised clustering to, e.g., identify novel patient subtypes.

*Input-data fusion* techniques aim to combine the sample information from multiple views to create a more comprehensive and integrated representation for subsequent unimodal analysis. While most literature works have presented input-data fusion methods that exploit joint factor analysis techniques (e.g. PCA-based techniques - JIVE [20], or joint non-negative matrix factorization techniques [76, 77, 78]) and use them for exploratory-data analysis or unsupervised clustering, preliminary experiments reported by MOFA+ [21], a Bayesian approach exploiting a latent variational-inference estimation of factors, show that the integrated view has the potential to improve the performance of supervised/unsupervised modeling. MOFA+ decomposes the $M$ input views into a matrix of shared latent factors for each sample and $M$ weight matrices, one for each data modality. The decomposition is guided by the assumption of prior-distributions that impose a weight-regularization disentangling variation across data sets while yielding interpretable factors. MOFA+ uses a two-level regularization: the first level encourages view- and factor-wise sparsity, thereby allowing to directly identify which factor is active in which view. The second level encourages feature-wise sparsity, thereby typically resulting in a small number of features with active weights.

In this work, we aimed at performing a comparative evaluation of the most used and successful data integration techniques, to also understand the impact of dimensionality reduction on their predictive performance. Due to their documented performance, we performed experiments by comparing the results obtained by the base view concatenation to those obtained by SNF, uMKL, and MOFA+.

#### S. B. Dataset preparation

The TCGA repository groups cancers by their type (see [79] and the list of cancer types in the dedicated web page), but often clusters together samples having different histological types, which may add dispersive information biasing any analysis.

To improve the homogeneicity in the considered datasets, for the following cancers, we selected a subset of the available histological types:

- BLCA (Bladder Urothelial Carcinoma): only "muscle invasive urothelial carcinoma (pt2 or above)" samples are considered.

- BRCA (Breast invasive carcinoma): this dataset was split into two separated datasets (BRCA1 and BRCA2) comprising “infiltrating ductal carcinoma” and “infiltrating lobular carcinoma”, respectively.

- LUAD (Lung adenocarcinoma): only “lung acinar adenocarcinoma”, “lung adenocarcinoma mixed subtype”, “lung adenocarcinoma- not otherwise specified (nos)”, “lung bronchioloalveolar carcinoma mucinous”, “lung bronchioloalveolar carcinoma nonmucinous”, “lung micropapillary adenocarcinoma” and “lung papillary adenocarcinoma” are considered.

- LUSC (Lung squamous cell carcinoma): only “lung squamous cell carcinoma - not otherwise specified (nos)” are considered.

- PRAD (Prostate adenocarcinoma): only “prostate adenocarcinoma acinar type” samples are considered.

The Kidney renal clear cell carcinoma (KIRC) dataset, the Ovarian serous cystadenocarcinoma (OV), and the Skin Cutaneous Melanoma dataset (SKCM) contained cancers belonging to the same histological type.

For Breast Invasive Carcinoma (BRCA), we consider only female patients which are the vast majority. Moreover, only primary solid tumors are exploited for all cancer types except Skin Cutaneous Melanoma (SKCM). Most of the patients in SKCM dataset are metastatic (following the rapid progression of the disease), thus we retain both primary and metastatic samples (when both are available we choose the primary). We controlled for the absence of technical replicates in the datasets and we further selected only samples not stored using “formalin-fixed paraffin-embedded” (FFPE) technique that leads to lower quality nucleic acids (DNA and RNA) that are fragmented and chemically modified [80]. miRNA, mRNA, protein expression and DNA methylation data are downloaded for each cancer type. In particular, the following assays are exploited for each data view:

- *miRNA*: Gene-level log2 RPM expression values from RNA-Sequencing

- *mRNA*: RSEM TPM gene expression values from RNA-Sequencing

- *protein*: normalized protein expression values from Reverse Phase Protein Array (RPPA)

- *DNA methylation*: Probe-level methylation beta values from Methylation Array.

In the case of DNA methylation, we used data coming from "Infinium HumanMethylation 450K BeadChip" in all cases except for the Ovarian serous cystadenocarcinoma (OV) dataset, where only few samples where available for this array and thus we opted for data with lower resolution coming from "Illumina HumanMethylation 27K BeadChip".

After downloading all the datasets we generated each view by aplying the following filters, which allow to prune variables carrying practically no information.

1. **Remove variables mainly due to noise by filtering near-zero variance features**: Features with near zero variance were defined as: (1) features with very few unique values with respect to sample cardinality (in our experiments we removed features that had less than $\frac{N}{10}$ unique values, being $N$ the number of cases in the dataset); (2) features for which the frequency of the most common value is twenty times larger than the frequency of the second most common value.[9]

2. **Remove high pairwise-feature correlations**. Pairs of features showing a high pairwise correlation (i.e. Pearson correlation $> 0.75$) were filtered to remove the features having the highest mean correlation with all the other features in the dataset[10]. The parallel algorithm we implemented to perform this task is outlined in supplementary section S. B.1.

3. **Data normalization**: each variable was normalized via standard scaling (Z-score normalization).

4. **Entropy Filtering**: the entropy of a signal is considered as a measure of the information content of the signal; to obtain practical computational times views containing more than 30000 variables were trimmed to 30000, features by selecting those features characterized by the highest entropy. Note that most authors [14, 15] use a similar quick filtering approach to both reduce computational costs and minimize the curse of dimensionality. However, they reduce all the input data-views to the same, much lower (e.g. 4000 [14] or 5000 [15] features) heuristic or empirically chosen, dimensionality.

[9]The function "nearZeroVar" from R package caret was used with default arguments.

[10]From a biological perspective, the correlation between omics variables could often be meaningful. However, from a statistical point of view, highly correlated variables lead to inflated estimates, therefore affecting the reliability of statistical estimators. Thus, their removal is often advisable.

> Thanks to the approach we are proposing we could instead choose a much larger value to obtain practical computational costs; effects due to the curse of dimensionality were instead reduced by more informed DR techniques.

Further we downloaded demographics patient variables reporting age at first pathological diagnosis, gender, ethnicity, and race. Tables S. B.1-S. B.3 report statistics about the patient demographics.

| | | OS = 0 | OS = 1 | p-value |
|---|---|---|---|---|
| **BLCA** | | | | |
| **n** | | 184 | 151 | |
| **age at first diagnosis** | | 66.00 [57.75, 75.00] | 70.00 [64.00, 77.50] | $< 0.001$* |
| **gender (%)** | male | 141 (76.6) | 110 (72.8) | 0.504 |
| | female | 43 (23.4) | 41 (27.2) | |
| **race (%)** | white | 137 (78.3) | 124 (85.5) | 0.001* |
| | black or african american | 7 (4.0) | 13 (9.0) | |
| | asian | 31 (17.7) | 8 (5.5) | |
| **ethnicity (%)** | not hispanic or latino | 159 (95.8) | 138 (100.0) | 0.04* |
| | hispanic or latino | 7 (4.2) | 0 (0.0) | |
| **BRCA1** | | | | |
| **n** | | 275 | 42 | |
| **age at first diagnosis** | | 56.00 [47.50, 64.00] | 59.00 [45.50, 73.25] | 0.314 |
| **race (%)** | white | 167 (60.9) | 37 (88.1) | 0.002* |
| | black or african american | 86 (31.4) | 5 (11.9) | |
| | asian | 21 (7.7) | 0 (0.0) | |
| **ethnicity (%)** | not hispanic or latino | 243 (95.7) | 41 (100.0) | 0.361 |
| | hispanic or latino | 11 (4.3) | 0 (0.0) | |
| **BRCA2** | | | | |
| **n** | | 114 | 14 | |
| **age at first diagnosis** | | 61.00 [49.25, 69.00] | 67.00 [60.50, 74.25] | 0.067 |
| **race (%)** | white | 98 (86.7) | 10 (76.9) | 0.553 |
| | black or african american | 8 (7.1) | 2 (15.4) | |
| | asian | 7 (6.2) | 1 (7.7) | |
| **ethnicity (%)** | not hispanic or latino | 105 (94.6) | 13 (100.0) | 0.86 |
| | hispanic or latino | 6 (5.4) | 0 (0.0) | |

Table S. B.1: Statistics for demographic variables in BLCA, BRCA1, and BRCA2 datasets. Numeric variables (age and age at first pathologic diagnosis) that were all non-normally distributed; therefore, kruskal-wallis test was used. For categorical variables, $\chi$-square test was used. Asterisks mark p-values that are lower than 0.05 (a statistically significant difference was found when comparing the distribution of cases different survival events.

| | | OS = 0 | OS = 1 | p-value |
|---|---|---|---|---|
| **KIRC** | | | | |
| **n** | | 121 | 48 | |
| **age at first diagnosis** | | 61.00 [51.00, 68.00] | 63.00 [54.75, 73.25] | 0.055 |
| **gender (%)** | male | 76 (62.8) | 33 (68.8) | 0.583 |
| | female | 45 (37.2) | 15 (31.2) | |
| **race (%)** | white | 92 (77.3) | 40 (85.1) | 0.108 |
| | black or african american | 27 (22.7) | 6 (12.8) | |
| | asian | 0 (0.0) | 1 (2.1) | |
| **ethnicity (%)** | not hispanic or latino | 98 (95.1) | 42 (97.7) | 0.807 |
| | hispanic or latino | 5 (4.9) | 1 (2.3) | |
| **LUAD** | | | | |
| **n** | | 180 | 120 | |
| **age at first diagnosis** | | 64.00 [59.00, 71.00] | 67.00 [58.00, 74.00] | 0.265 |
| **gender (%)** | male | 82 (45.6) | 54 (45.0) | 1 |
| | female | 98 (54.4) | 66 (55.0) | |
| **race (%)** | white | 136 (83.4) | 92 (87.6) | 0.611 |
| | black or african american | 24 (14.7) | 12 (11.4) | |
| | asian | 3 (1.8) | 1 (1.0) | |
| **LUSC** | | | | |
| **n** | | 135 | 93 | |
| **age at first diagnosis** | | 67.00 [60.25, 73.00] | 70.00 [64.00, 74.00] | 0.037* |
| **gender (%)** | male | 101 (74.8) | 74 (79.6) | 0.499 |
| | female | 34 (25.2) | 19 (20.4) | |
| **race (%)** | white | 103 (91.2) | 72 (88.9) | 0.084 |
| | black or african american | 6 (5.3) | 9 (11.1) | |
| | asian | 4 (3.5) | 0 (0.0) | |

Table S. B.2: Statistics for demographic variables in KIRC, LUAD, and LUSC datasets. Numeric variables (age and age at first pathologic diagnosis) that were all non-normally distributed; therefore, kruskal-wallis test was used. For categorical variables, $\chi$-square test was used. Asterisks mark p-values that are lower than 0.05 (a statistically significant difference was found when comparing the distribution of cases different survival events.

| | | OS = 0 | OS = 1 | p-value |
|---|---|---|---|---|
| **OV** | | | | |
| **n** | | 83 | 143 | |
| **age at first diagnosis** | | 57.00 [50.00, 63.00] | 59.00 [53.00, 69.50] | 0.035* |
| **PRAD** | | | | |
| **n** | | 331 | 6 | |
| **age at first diagnosis** | | 62.00 [57.00, 66.00] | 62.50 [53.75, 69.00] | 0.948 |
| **SKCM** | | | | |
| **n** | | 184 | 150 | |
| **age at first diagnosis** | | 58.00 [48.00, 71.75] | 56.50 [48.00, 71.00] | 0.824 |
| **gender (%)** | male | 97 (52.7) | 99 (66.0) | 0.019* |
| | female | 87 (47.3) | 51 (34.0) | |
| **race (%)** | white | 173 (97.2) | 142 (95.9) | 0.755 |
| | asian | 5 (2.8) | 6 (4.1) | |
| **ethnicity (%)** | not hispanic or latino | 173 (97.2) | 146 (98.6) | 0.603 |
| | hispanic or latino | 5 (2.8) | 2 (1.4) | |

Table S. B.3: Statistics for demographic variables in OV, PRAD, and SKCM datasets. Numeric variables (age and age at first pathologic diagnosis) that were all non-normally distributed; therefore, kruskal-wallis test was used. For categorical variables, $\chi$-square test was used. Asterisks mark p-values that are lower than 0.05 (a statistically significant difference was found when comparing the distribution of cases different survival events.

*S. B.1. High-pairwise correlation analysis and filtering*

We considered as highly-correlated pairs of features with a Pearson correlation coefficient $corr_P > 0.75$. For each pair of correlated variables, we discarded the one whose mean absolute pairwise-correlation with respect to all the other features is higher. Since the application of pairwise-correlation filtering is computationally demanding, we implemented a parallel-distributed algorithm that applies the following consecutive steps.

1. The feature-set is split into non-intersecting feature sub-subsets; in other words, given the input dataset $\mathbf{X} \in \Re^{N \times D}$, where $N$ is the number of cases and $D$ is the number of features, we create $M$ non intersecting $d$-dimensional feature-subsets $\mathbf{X}^j_{sub} \in \Re^{N \times d}, \bigcap_{j=1,\dots,M} \mathbf{X}^j_{sub} = \emptyset$. The value of $d$ was set to $d = 5000$ to obtain practical computational costs.

2. Each $\mathbf{X}^j_{sub}$ is assigned to a core, where it is processed to remove high-pairwise correlations, therefore obtaining a filtered subset $\mathbf{X}^j_{subF}$.

3. All the filtered subsets are recollected by the main thread and are recomposed by concatenation along the feature-dimension to obtain a new (partially filtered) dataset $\mathbf{X}_F = \left[\mathbf{X}^1_{subF}, \ldots, \mathbf{X}^M_{subF}\right]$.

4. The position of features in $\mathbf{X}_F$ is shuffled and the process (points 1 to 3) is iterated on $\mathbf{X}_F$ until a maximum number of iterations (set to 10) has been reached or no high-correlated features have been removed in any of the subsets.

### S. C. Automatic detection of plateau in the distribution of block-`ids` and block-`ids` plots

To automatically identify a plateau on the distribution of the block-`ids` and reduce the computational costs by stopping computation when a plateau is found, we view the distribution of the block-`ids` as a signal and process it by applying:

(1) the standard deviation filter, with support $wl = 10$ and anchored on the right-most element of the window;

(2) the box-derivative filter, which is realized by applying a derivative filter, with support $wl = 3$ and anchored on the central point of the filtered window, followed by a box (or average) filter, with support $wl = 10$ and anchored on the right-most element of the filtered window.

Scanning the results of the two filters from left (smallest block) to right (largest block) a plateau is found at the first block where the derivative filter outputs a value lower than 0.25 or the average-derivative filter outputs a value lower than 0.05.

Placing the anchor of each filter on the point corresponding to the largest (right-most) block means that we can apply the two filters after the computation of each block-`id`. As a plateau is found, we check that the `id` of the block where the plateau is detected is less than three standard deviations away from the two-nn `id` of the whole view, $\hat{d}_{twonn}(\mathbf{X})$. If this happens, the computation is stopped; otherwise it continues and searches for the next block.

In figures S. C.1-S. C.4 we plot the results of the block analysis for all the considered TCGA datasets. For visualization purposes, the plots visualize the computation over all the blocks, until the block size covers all the input view.

### S. D. Hierarchical feature clustering

Besides exploration of relationships between features, hierarchical feature clustering allows selecting a lower-dimensional subspace composed of "representative" features, that are feature-cluster medoids or singletons (that is, features that do not belong to any cluster). In this work, feature clusters were composed by an agglomerative hierarchical clustering method [81] named Genie [35]. Besides being one of the fastest hierarchical clustering techniques, Genie guarantees robustness with respect to noise and boundary points by computing pairwise-points distances leveraging the mutual reachability distance [82]. Further, it applies a Single Linkage (SL) criterion [83] to merge closer clusters but also provides a strategy to penalize the formation of small clusters. In particular, at each iteration of the algorithm, the inequality between the cluster cardinalities is evaluated by the Gini index. If the inequality is above a user-set threshold Genie starts favoring the merge of the smallest clusters with their nearest clusters. In high dimensions the computational costs of Genie become impracticable; hence we implemented a distributed-hierarchical algorithm (see supplementary section S. D.1) and we applied it to extract a number of feature clusters (i.e. the dimension of the reduced space) equal to the desired dimension of the lower dimensional space.

#### *S. D.1. Distributed-clustering through Genie*

To optimize the feature clustering, we designed a parallel algorithm similar to the one implemented for high-pairwise correlation filtering (supplementary section S. B.1) that iteratively clusters the feature set until the space dimension is greater than the desired number of clusters, $\mathcal{C}$. In more detail, the following steps are applied:

1. The feature-set is split into non-intersecting feature sub-subsets; in other words, given the input dataset $\mathbf{X} \in \Re^{N \times D}$, being $N$ the number of cases and $D$ the number of features, we create $M$ non intersecting $d$-dimensional feature-subsets

   $\mathbf{X}_{sub}^{j} \in \Re^{N \times d}, \bigcap_{j=1,\ldots,M} \mathbf{X}_{sub}^{j} = \emptyset$. The value of $d$ was set to $d = 1000$ to obtain practical computational costs.

2. Each $\mathbf{X}_{sub}^{j}$ is assigned to a core, where Genie is applied to apply hierarchical clustering and then extract $c$ clusters ($c$ is chosen at the beginning of the algorithm in order to ensure a maximum number of iterations are

applied). The $c$ features that are the medoids of the $c$ clusters are the representative features that are selected to represent $\mathbf{lowX}^j_{sub} \in \Re^{N \times c}$, the lower-dimensional representation of $\mathbf{X}^j_{sub}$.

3. All the $\mathbf{lowX}^j_{sub}$ are collected by the main thread and are concatenated along the feature dimension to obtain
   $\mathbf{lowX} = \left[\mathbf{lowX}^1_{sub}, \ldots, \mathbf{lowX}^M_{sub}\right]$, the lower dimensional representation of $\mathbf{X}$.

4. The position of features in $\mathbf{lowX}$ is shuffled and points 1 to 3 are iterated until the dimension of $\mathbf{lowX}$ becomes lower or equal than $C$.

### S. E. Parallel implementation of iterative RCUR

The CUR decomposition, and its more promising randomized version RCUR [36, 51, 52], have been proposed as an interpretable dimensionality reduction alternative to PCA and RPCA (Randomized PCA, alias Randomized SVD); indeed, the novel dimensions computed by PCA and RPCA have a meaning that is difficult to interpret [55]. CUR factorizes the initial matrix $\mathbf{X} \in \Re^{N \times D}$ into 3 matrices: $\mathbf{C} \in \Re^{N \times d}$, $\mathbf{U} \in \Re^{d \times d}$, and $\mathbf{R} \in \Re^{d \times D}$, where $\mathbf{C}$ and $\mathbf{R}$ are formed by a subset of columns and rows from $\mathbf{X}$, which are chosen based on their capability of maintaining the structure and informative content of the $\mathbf{X}$, and $\mathbf{U}$ is constructed so as to guarantee that the product CUR is close to $\mathbf{X}$. When dimensionality reduction by CUR (or RCUR) is applied, the lower-dimensional dataset is represented by the $d$-dimensional $\mathbf{C}$ matrix. The $d$ features composing $\mathbf{C}$ are selected based on their potential to represent the information in $\mathbf{X}$; this potential is quantified by the *normalized statistical leverage score*, a coefficient computed based on the top $d$ right singular vectors of $\mathbf{X}$ (for more details see [36]). To compute the singular vectors, the SVD or truncated SVD algorithm are generally applied, which means that the CUR and RCUR algorithms will, at most, select a number of features $d \leq min(N,D) - 1$.

To apply RCUR for feature selection and obtain a number of selected features that is $L_p \geq min(N,D) - 1$ we used a parallel algorithm that applies RCUR several times (e.g. 1000) and, after each application, assigns a win to any of the features selected by RCUR. Once all the RCUR iterations have been performed, the algorithm selects the $L_p$ features that obtained the largest number of wins.

### S. F. Comparison between experiments applying no DR, experiments using heuristics to define the dimension of the lower dimensional space, and experiments where block-analysis guides the DR

In this section we firstly provide the preliminary experiments that evidenced the urgent need of applying a properly designed DR approach (supplementary section S. F.1). Next, in supplementary section S. F.2 we report details about the experiments that allowed us to understand that the usage of the block-analysis to guide the DR is a promising way, avoiding difficult empirical choices.

#### *S. F.1. Application of data fusion without any prior DR step*

Our first experiment regarded the straightforward application of data-fusion algorithms on the nine TCGA datasets that were pre-processed to remove high-pairwise correlations and low variance, and were then input to entropy filtering (supplementary section S. B) to have a maximum of 30000 variables per view. Figure S. F.5 shows results obtained on the fifteen stratified holdouts when we avoided hyper-parameter tuning and supervised feature selection prior to RF training. Though pre-processing allowed obtaining an AUC surpassing the one of a random classifier for several datasets and data-fusion algorithms, results were far from being satisfactory; when supervised feature selection and hyper-parameter tuning were applied prior to RF training (see figure S. F.6) as described in supplementary section S. F.4, the performance improvement was almost unnoticeable, probably due to the high-number of variables relative to the limited cardinality of the samples in the internal training holdouts.

This cleared that the high-data sparsity, affecting especially the DNA methylation and mRNA views, was a source of large bias misleading even supervised tuning steps; an effective DR approach was therefore necessary.

#### *S. F.2. Comparison between heuristics $HD_1$, $HD_2$ and models using the block-analysis to guide DR*

The next step aimed at assessing the effectiveness of using the `id` estimate to set the dimension of the lower dimensional space. To this aim, we ran experiments where the one-step (feature-selection or feature-extraction) DR pipelines described in section 4.1 used heuristically set dimensions for the lower dimensional space, $\bar{d}_{H*}$. In particular, the heuristic dimensions we chose to compare are based on the rationale that most of the feature extraction algorithms allow to compute a reduced space whose number of dimensions is lower or equal than $min(N,D)-1$ [36, 41, 42]. Based on this consideration, we ran all the one-step

DR plus data integration pipelines by using two heuristics, $HD_1$ and $HD_2$. In particular, $HD_1$ sets the dimension of the reduced space to $\hat{d}_{H1} = min(N,D) - 1$; $HD_2$ halves $\hat{d}_{H1}$, i.e. for $HD_2$ we used $\hat{d}_{H2} = \frac{min(N,D)}{2}$.

Our first comparative evaluation (supplementary section S. F.2.1) was aimed at identifying the best performing among the two heuristic definitions ($HD_1$ and $HD_2$) of the dimension of the lower dimensional space. The chosen heuristic was then compared to the results obtained when block-analysis was used to guide the DR (supplementary section S. F.2.2).

#### *S. F.2.1. Comparative evaluation of pipelines using heuristics $HD_1$ and $HD_2$ for setting the dimension of the reduced spaces*

To obtain an initial straightforward comparison, we firstly ran a Wilcoxon signed-rank test to compare all the results obtained by $HD_1$ to those obtained by $HD_2$. In other words, we paired the results based on the dataset being considered, the employed DR and data-fusion algorithms, and the holdout. The obtained p-values showed that there is no statistically significant difference when the results are compared based on the AUC (p-value $p < 0.0871$); on the other hand, Wilcoxon signed-rank test based on the AUCPR highlighted that the lower dimensional space defined by heuristic $HD_2$ provides more accurate results (p-value $p < 0.0238$).

To choose between the two heuristics we ran more exhaustive pairwise comparisons by using the sided Wilcoxon signed-rank test to compare all the pipelines experimented for $HD_1$ to those experimented for $HD_2$. Extracts of the win-tie-loss tables summarizing the results of the application of the sided Wilcoxon signed-rank test are shown in figure S. F.7, when the AUC is used for comparison (detailed win-tie-loss tables are reported in supplementary file *S*1), and figure S. F.8 (supplementary table *S*2) when the AUCPR is used for comparison. The obtained win-tie-loss tables supported the belief that, on average, $HD_2$ obtains more reobust AUCPR results.

To provide a more comprehensive comparison in Supplementary table *S*3, for each dataset, we collected the three DR+data-fusion pipelines that obtained the best AUC and/or the best AUCPR. As shown in figure S. F.9, among the forty top-performing models we collected, twenty-five experiments (63% of all the experiments we run) used $HD_2$, while fifteen (37%) used $HD_1$. This further supported our choice of using heuristic $HD_2$ as a benchmark to assess our proposal of using the block-analysis and the estimated `id` to guide DR.

*S. F.2.2. Comparative evaluation of DR and data-fusion pipelines using heuristic $HD_2$ to set the dimension of the reduced space versus pipelines guided by block-analysis*

When we compared heuristic $HD_2$ to our proposal, we firstly analyzed each cancer dataset and, for each of them, we extracted the three DR+data-fusion pipelines (experiments) that obtained the highest AUC and the highest AUCPR values (see supplementary table S4). The 76% of the top-performing DR+data-fusion experiments (32/42) are those that use the block-analysis to guide the DR (figure S. F.10).

This results are confirmed by the sided Wilcoxon tests we ran to compare all the DR+data-fusion pipelines (on all the nine TCGA datasets) guided by $HD_2$ to those guided by the block-analysis. For this comparison, the Wilcoxon test was unpaired because block-analysis allows to also implement two-step DR approaches, for which there is no pairing when using $HD_2$. Further, before running the tests we considered that we were pooling results obtained on all the datasets, which are characterized by different balance between positive and negative samples (tables 1 and 2). While AUCs have a clear baseline value across different datasets, which corresponds to the AUC $= 0.5$ of a random classifier, the baseline for the AUCPR measure, i.e. the AUCPR of a random classifier, generally corresponds to the ratio of positive examples [84] in the dataset, and therefore varies across the nine datasets we are considering. For this reason, for the unpaired Wilcoxon test we used $\Delta(\text{AUCPR})$ measure, computed as the difference between the AUCPR and the ratio of positive examples in the dataset.

The obtained p-values (p-value $p < 0.013$ when using the AUC, and p-value $p < 0.033$ when using the $\Delta(\text{AUCPR})$) are a first evidence of the superiority of models guided by block analysis.

Next, we performed detailed pairwise comparisons between all DR+data-fusion pipelines. Note that, in this case, we could pair the holdouts and cancer datasets. Extracts of the win-tie-loss tables obtained when using the AUC and the AUCPR measures to run sided Wilcoxon signed-rank tests are shown in figures S. F.11 S. F.12 (detailed tables are reported in supplementary tables S5 - when AUC is used for hypothesis testing, and S6 - when AUCPR is used for hypothesis testing). They highlight that the usage of the block-analysis to guide the DR is a promising way, especially when the crucial AUCPR measure is used, which is often the more restrictive evaluation measure in unbalanced datasets.

#### *S. F.3. Comparing different settings when applying DR+data-fusion pipelines guided by the block-analysis*

In this section, we report details about the settings we experimented when using block-analysis to guide DR.

For each setting, we report extracts of the win-tie-loss tables (obtained with the AUC and AUCPR measures) containing the twenty-five top-winners DR+data-fusion experiments. When comparing two DR+data-fusion results, Wilcoxon signed-rank test pairs the results achieved on the same TCGA dataset and holdout.

In particular, we experimented with the following four settings:

- all the available omics being integrated (section 4.3.2): extracts of twenty five top-winning (most robust) DR+data-fusion pipelines are reported in figures S. F.13 and S. F.14. The complete win-tie-loss tables are reported in supplementary tables S7 (win-tie-losses obtained when the AUC measure is used), S8 (win-tie-losses obtained when the AUCPR measure is used), and S9 (list of top-performing models according to the AUC or AUCPR measures).

- at least two omics being integrated (section 4.3.3): extracts of twenty five top-winning DR+data-fusion pipelines are reported in figures S. F.15 and S. F.16. The complete win-tie-loss tables are reported in supplementary tables S10 (win-tie-losses obtained when the AUC measure is used), S11 (win-tie-losses obtained when the AUCPR measure is used), and S12 (list of top-performing models according to the AUC or AUCPR measures).

- all the available omics plus patients' demographic predictors (section 4.3.4) extracts of twenty five top-winning DR+data-fusion pipelines are reported in figures S. F.17 and S. F.18). The complete win-tie-loss tables are reported in supplementary tables S13 (win-tie-losses obtained when the AUC measure is used), S14 (win-tie-losses obtained when the AUCPR measure is used), and S15 (list of top-performing models according to the AUC or AUCPR measures).

  For what regards this setting, in the next supplementary subsection S. F.3.1 we describe the ascertain bias and the data fairness issues that must be carefully considered when interpreting classifiers using patient data other than non-omics for supervised/unsupervised classification.

*S. F.3.1. Ascertain bias and data fairness*

Several multi-omics studies provide patients' descriptors (demographic, clinical, biological data [3]) differing from multi-omics, which could aid the following unsupervised/supervised analysis, when opportunely integrated.

Indeed, the results we report in section 4.3.4 show that a simple integration by concatenation of demographic descriptors may improve the classification performance.

We however warn that, when non-omics data is integrated to omics views, the interpretation and explanation of the obtained predictions to, e.g., uncover triggers of mortality risks, should be carefully performed due to ascertain bias, data quality issues, and dataset/model fairness.

Ascertain bias is related to individuals enriched for the risk factors under study so that strong correlations between multi-omics and (demographic but also clinical) patient data may be present, which might induce inflated, biased estimates. For what regards the demographics variables, we must indeed note that omics profiles have been already shown to be related to demographic descriptors [85].

On the other hand, even the most relevant biomedical studies tend to be characterized by quality issues not easy to control [86] and related to limited follow-up time and/or a large number of values being missing in patient data, which contrast with the completeness of multi-omics variables. This may bias the effect estimates of the clinical/demographic variables. Note that the demographic variables we selected (i.e. age at first pathological diagnosis, gender, ethnicity, and race) are less prone to quality issues because they do not require a follow-up and are generally nearly complete (limited number of missing data).

Data fairness refers to the fact that both supervised and unsupervised models are depending of the available dataset that, as a matter of fact, embodies the bias generated in the social context that originated it [87]. As an example, several papers using ML techniques on COVID data from US hospitals have documented the highest age-adjusted infection rates and risk for Hispanic/Latino immigrants or other minorities [88, 89], which may push the machine learning algorithm to reinforce and evidence healthcare disparities while underestimating or even disregarding the underlying biomedical causes of infection risks. Therefore, when demographic variables are used, caution should be taken when analyzing and interpreting the obtained classifications.

For what regards our study, descriptive statistics showed that age at diagnosis, race, and ethnicity descriptors have a relationship with survival events in the BLCA (p-value $< 0.05$, table S. B.1), race is also associated with overall survival events in the BRCA1 (p-value $< 0.05$, table S. B.1) dataset, gender is related to the overall survival events in the SKCM dataset (p-value $< 0.05$), and age at diagnosis is also related to overall survival events in the LUSC (table S. B.2), and OV datasets (table S. B.3).

Note that, while some authors have already reported some biological explanations for racial disparity in survival to breast cancer [90], other works have linked survival to bladder cancer with age and ethnicity [91] and therefore suggested including these factors in survival prediction, survival to ovarian cancer has been related to age at first pathological diagnosis [92] and the same relationship holds for squamous non-small cell lung cancer [93]. However, we found no literature evidence of gender-dependent survival events for skin cutaneous melanoma [94].

Figures S. F.17 and S. F.18 report the extracts of the top-winner DR+data-fusion experiments (specified also by the combination of the input views) that combine the four omics and eventually integrate the demographic view. It is undoubted that the inclusion of demographic views improves the classification performance.

#### *S. F.4. Supervised Feature Selection and hyper-parameter tuning steps*

The supervised feature selection step applied to improve RF performance used 21 internal stratified holdouts - obtained by random selection of the 90% of training cases. Each holdout was then used to train an RF, and the trained RF was then used to compute a permutation-importance value for each variable. After processing all the internal holdouts, all the importances were averaged across the 21 holdouts to obtain a mean permutation-importance for each variable.

At this stage, the most discriminative variables were selected by normalizing the mean importances to unitary sum, computing their cumulative sum, and finally selecting those (most important) variables allowing to obtain a cumulative sum greater than 0.9.

Hyper-parameter tuning was performed by using internal holdout validation; more precisely we chose the combination of parameters that allowed obtaining the best average AUCPR value across 21 stratified holdouts containing the 90% of samples in the dataset. This procedure was used to tune only the number of trees and the value of the number $mtry$ of variables that are considered to form each split. More precisely, for the number of trees we tested the following values $ntree = \{100, 250, 500, 750, 1000, 1250\}$. To define the search values for the value of $mtry$ we considered that its default value $m\bar{t}ry$ is defined as the square of the number of variables in the dataset; we set the search space based on $m\bar{t}ry$, i.e. we tested the following search space $mtry = \{0.5m\bar{t}ry, m\bar{t}ry, 1.5m\bar{t}ry\}$.

More careful supervised feature selection or hyper-parameter tuning strategies may further improve results.

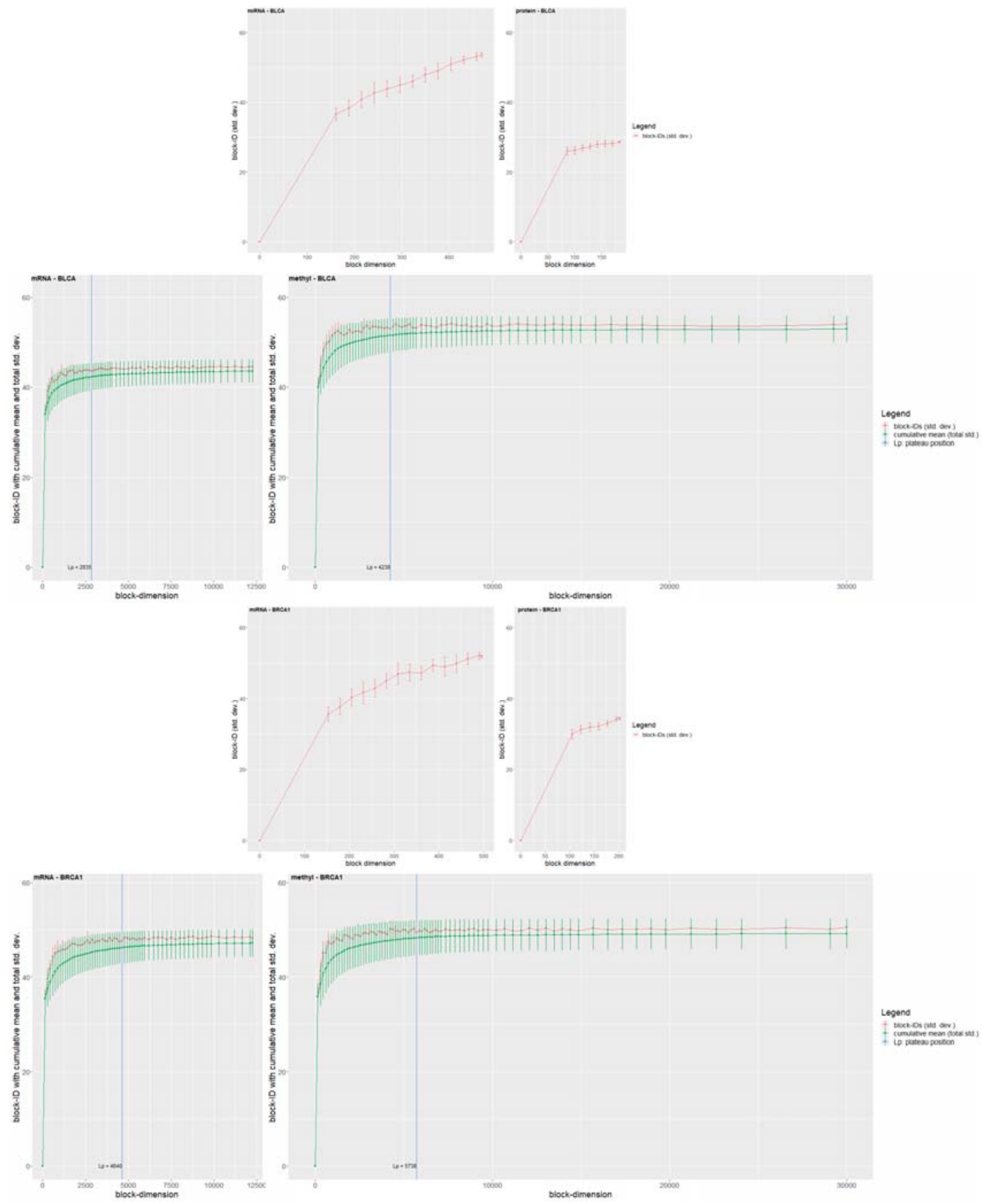

Figure S. C.1: Block analysis for BLCA and BRCA1 datasets.

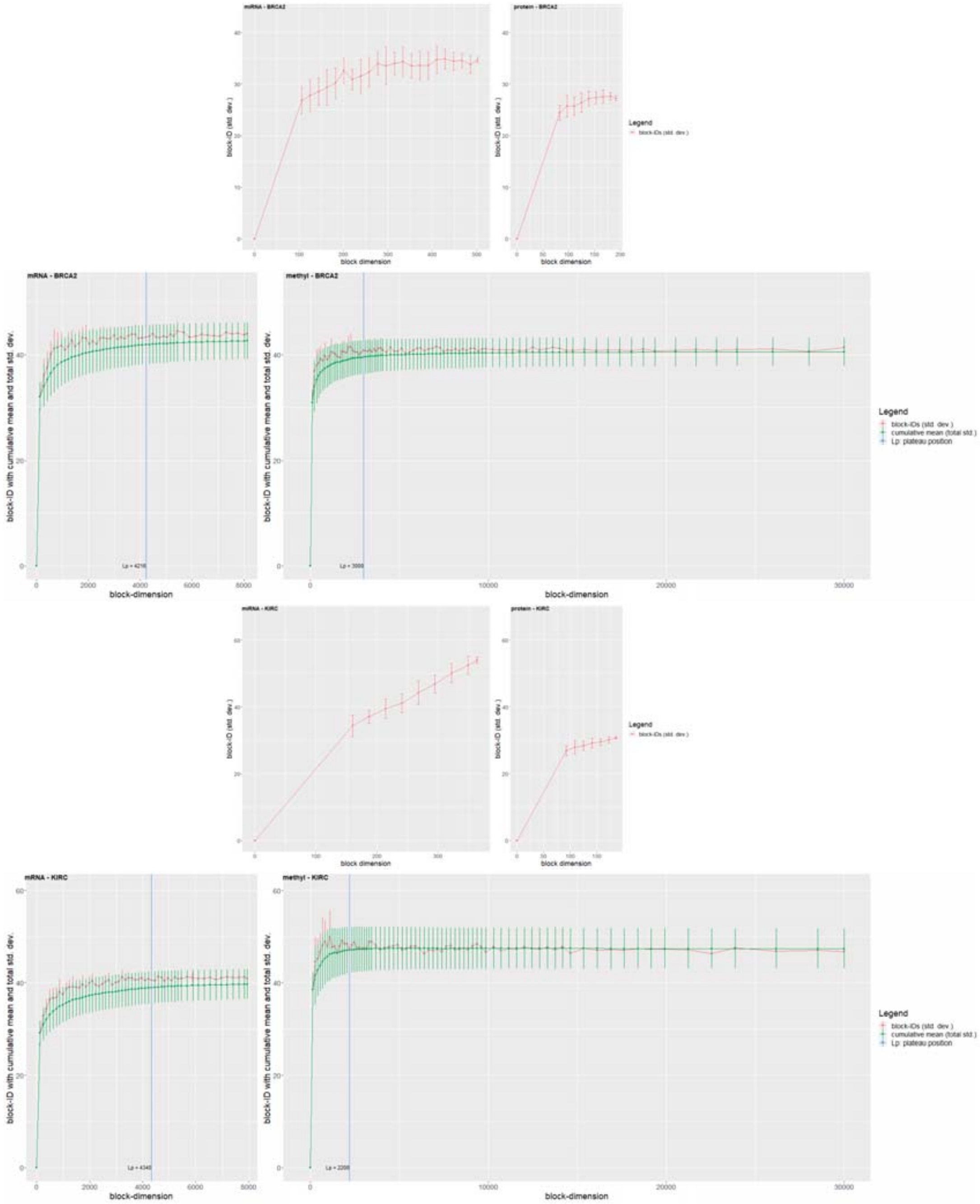

Figure S. C.2: Block analysis for BRCA2 and KIRC datasets.

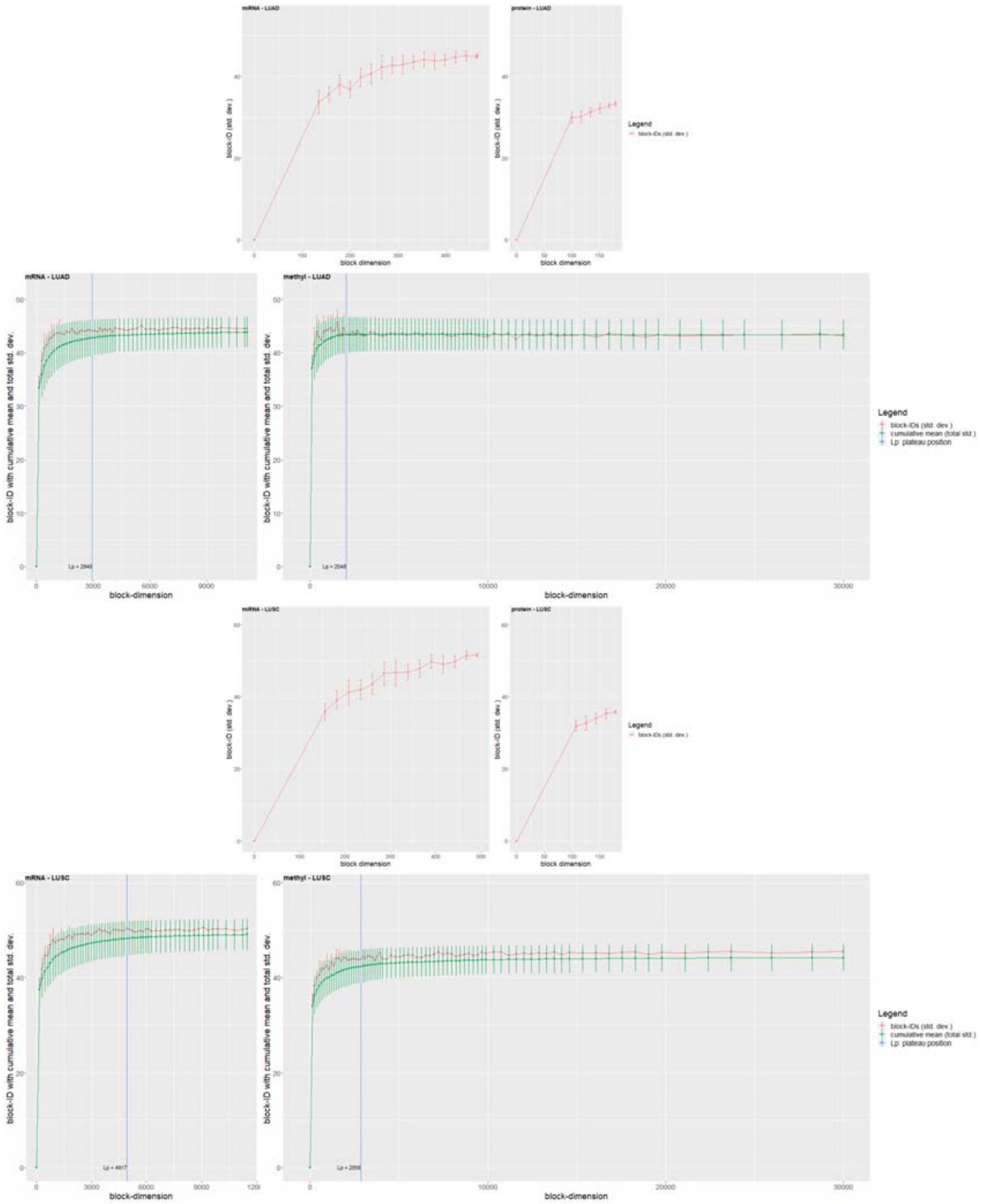

Figure S. C.3: Block analysis for LUAD and LUSC datasets.

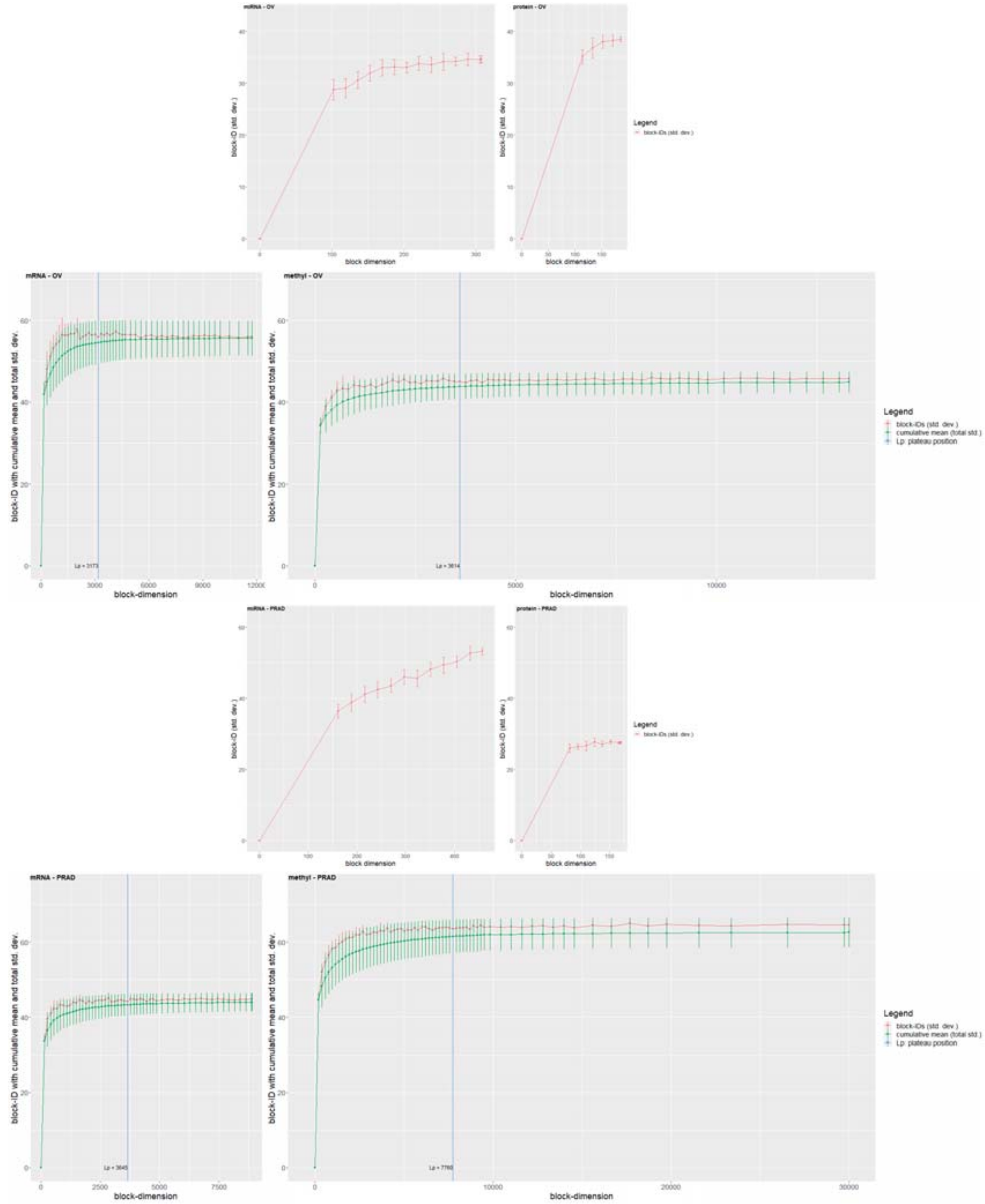

Figure S. C.4: Block analysis for the OV and PRAD datasets.

| Dataset | N | d | data fusion | aucpr | std(aucpr) | auc | std(auc) |
|---|---|---|---|---|---|---|---|
| BLCA | 335 | 336 | SNF | 0.59 | 0.07 | 0.63 | 0.06 |
| | | 16 | MOFA+ | 0.55 | 0.07 | 0.59 | 0.06 |
| | | 336 | uMKL | 0.56 | 0.06 | 0.62 | 0.05 |
| BRCA1 | 317 | 318 | SNF | 0.25 | 0.08 | 0.53 | 0.08 |
| | | 16 | MOFA+ | 0.28 | 0.07 | 0.61 | 0.09 |
| | | 318 | uMKL | 0.28 | 0.07 | 0.57 | 0.08 |
| BRCA2 | 128 | 129 | SNF | 0.48 | 0.22 | 0.55 | 0.23 |
| | | 16 | MOFA+ | 0.39 | 0.06 | 0.59 | 0.23 |
| | | 129 | uMKL | 0.47 | 0.19 | 0.53 | 0.29 |
| KIRC | 169 | 170 | SNF | 0.50 | 0.14 | 0.67 | 0.10 |
| | | 16 | MOFA+ | 0.48 | 0.10 | 0.69 | 0.08 |
| | | 170 | uMKL | 0.50 | 0.13 | 0.66 | 0.11 |
| LUAD | 300 | 301 | SNF | 0.50 | 0.06 | 0.58 | 0.06 |
| | | 16 | MOFA+ | 0.58 | 0.06 | 0.63 | 0.04 |
| | | 301 | uMKL | 0.47 | 0.07 | 0.56 | 0.07 |
| LUSC | 228 | 229 | SNF | 0.51 | 0.07 | 0.59 | 0.08 |
| | | 16 | MOFA+ | 0.45 | 0.05 | 0.50 | 0.06 |
| | | 229 | uMKL | 0.50 | 0.07 | 0.58 | 0.08 |
| OV | 226 | 227 | SNF | 0.69 | 0.05 | 0.54 | 0.09 |
| | | 16 | MOFA+ | 0.67 | 0.06 | 0.48 | 0.08 |
| | | 227 | uMKL | 0.70 | 0.06 | 0.56 | 0.10 |
| PRAD | 337 | 338 | SNF | 0.52 | 0.02 | 0.52 | 0.15 |
| | | 16 | MOFA+ | 0.52 | 0.01 | 0.32 | 0.34 |
| | | 338 | uMKL | 0.51 | 0.00 | 0.32 | 0.11 |
| SKCM | 334 | 335 | SNF | 0.65 | 0.07 | 0.68 | 0.05 |
| | | 16 | MOFA+ | 0.59 | 0.07 | 0.64 | 0.06 |
| | | 335 | uMKL | 0.64 | 0.05 | 0.65 | 0.04 |

Figure S. F.5: Tabular visualization of AUC and AUCPR obtained on all the cancer datasets when no hyper-parameter tuning nor supervised feature selection are applied prior to RF training.

| Dataset | N | d | data fusion | aucpr | std(aucpr) | auc | std(auc) |
|---|---|---|---|---|---|---|---|
| BLCA | 335 | 336 | SNF | 0.59 | 0.06 | 0.63 | 0.06 |
| | | 16 | MOFA+ | 0.54 | 0.06 | 0.57 | 0.07 |
| | | 336 | uMKL | 0.56 | 0.06 | 0.63 | 0.06 |
| BRCA1 | 317 | 318 | SNF | 0.22 | 0.06 | 0.49 | 0.07 |
| | | 16 | MOFA+ | 0.25 | 0.08 | 0.57 | 0.08 |
| | | 318 | uMKL | 0.29 | 0.08 | 0.57 | 0.08 |
| BRCA2 | 128 | 129 | SNF | 0.37 | 0.07 | 0.49 | 0.21 |
| | | 16 | MOFA+ | 0.49 | 0.25 | 0.58 | 0.30 |
| | | 129 | uMKL | 0.41 | 0.14 | 0.51 | 0.26 |
| KIRC | 169 | 170 | SNF | 0.48 | 0.12 | 0.66 | 0.09 |
| | | 16 | MOFA+ | 0.47 | 0.10 | 0.67 | 0.08 |
| | | 170 | uMKL | 0.50 | 0.12 | 0.64 | 0.10 |
| LUAD | 300 | 301 | SNF | 0.50 | 0.06 | 0.58 | 0.06 |
| | | 16 | MOFA+ | 0.55 | 0.06 | 0.61 | 0.06 |
| | | 301 | uMKL | 0.48 | 0.07 | 0.56 | 0.06 |
| LUSC | 228 | 229 | SNF | 0.51 | 0.09 | 0.59 | 0.08 |
| | | 16 | MOFA+ | 0.44 | 0.07 | 0.49 | 0.10 |
| | | 229 | uMKL | 0.51 | 0.07 | 0.58 | 0.08 |
| OV | 226 | 227 | SNF | 0.69 | 0.06 | 0.54 | 0.10 |
| | | 16 | MOFA+ | 0.66 | 0.06 | 0.48 | 0.08 |
| | | 227 | uMKL | 0.71 | 0.08 | 0.56 | 0.10 |
| PRAD | 337 | 338 | SNF | 0.52 | 0.03 | 0.57 | 0.16 |
| | | 16 | MOFA+ | 0.52 | 0.01 | 0.35 | 0.32 |
| | | 338 | uMKL | 0.51 | 0.00 | 0.28 | 0.10 |
| SKCM | 334 | 335 | SNF | 0.65 | 0.06 | 0.68 | 0.05 |
| | | 16 | MOFA+ | 0.56 | 0.06 | 0.63 | 0.06 |
| | | 335 | uMKL | 0.65 | 0.05 | 0.65 | 0.04 |

Figure S. F.6: Tabular visualization of AUC and AUCPR obtained on all the cancer datasets when supervised feature selection and hyper-parameter tuning are applied to maximize AUCPR prior to RF training.

| DR pipeline | data_integration | ID definition | wins | ties | losses | auc | std(auc) |
|---|---|---|---|---|---|---|---|
| rpca | SNF | HD_1 | 41 | 7 | 0 | 0.594 | 0.125 |
| rpca | SNF | **HD_2** | 40 | 8 | 0 | 0.589 | 0.126 |
| feature_clustering | SNF | **HD_2** | 39 | 9 | 0 | 0.591 | 0.136 |
| umap | SNF | **HD_2** | 38 | 10 | 0 | 0.583 | 0.163 |
| feature_clustering | SNF | HD_1 | 33 | 14 | 1 | 0.576 | 0.134 |
| feature_clustering | uMKL | HD_1 | 30 | 15 | 3 | 0.571 | 0.15 |
| entropy | uMKL | **HD_2** | 29 | 19 | 0 | 0.577 | 0.144 |
| entropy | SNF | **HD_2** | 29 | 19 | 0 | 0.573 | 0.158 |
| feature_clustering | concatenation | HD_1 | 28 | 16 | 4 | 0.559 | 0.162 |
| umap | SNF | HD_1 | 27 | 18 | 3 | 0.563 | 0.155 |
| entropy | concatenation | **HD_2** | 23 | 21 | 4 | 0.55 | 0.163 |
| feature_clustering | MOFA+ | HD_1 | 23 | 20 | 5 | 0.557 | 0.141 |
| rpca | uMKL | **HD_2** | 22 | 26 | 0 | 0.572 | 0.151 |
| rpca | MOFA+ | **HD_2** | 22 | 22 | 4 | 0.559 | 0.122 |
| entropy | concatenation | HD_1 | 22 | 21 | 5 | 0.547 | 0.182 |
| umap | concatenation | **HD_2** | 21 | 22 | 5 | 0.555 | 0.164 |
| feature_clustering | uMKL | **HD_2** | 21 | 19 | 8 | 0.558 | 0.14 |
| rpca | MOFA+ | HD_1 | 20 | 25 | 3 | 0.554 | 0.14 |
| feature_clustering | concatenation | **HD_2** | 20 | 23 | 5 | 0.55 | 0.151 |
| umap | concatenation | HD_1 | 20 | 22 | 6 | 0.551 | 0.164 |
| entropy | MOFA+ | HD_1 | 19 | 18 | 11 | 0.55 | 0.118 |
| tsne | concatenation | HD_1 | 15 | 24 | 9 | 0.553 | 0.166 |
| feature_clustering | MOFA+ | **HD_2** | 15 | 22 | 11 | 0.542 | 0.156 |
| entropy | SNF | HD_1 | 11 | 24 | 13 | 0.54 | 0.153 |
| rpca | concatenation | HD_1 | 11 | 20 | 17 | 0.548 | 0.14 |

Figure S. F.7: Tabular visualization of win-tie-loss of the models that had the largest number of wins when using the AUC measure in the Wilcoxon signed-rank test comparing DR and data-fusion pipelines when heuristics $HD_1$ (column "ID definition" $= HD\_1$) or $HD_2$ (column "ID definition" $= HD\_2$) are used to set the dimension of the reduced space. The average values of AUC and their standard deviation across all the cancer datasets are shown. Selecting the thirteen experiments that were the top-winners and also had zero losses (highlighted with yellow in the figure), eight (61%) used heuristic $HD_2$ (49% used $HD_1$).

| DR pipeline | data_integration | ID definition | wins | ties | losses | aucpr | std(aucpr) |
|---|---|---|---|---|---|---|---|
| entropy | uMKL | **HD_2** | 40 | 8 | 0 | 0.532 | 0.148 |
| feature_clustering | SNF | **HD_2** | 35 | 13 | 0 | 0.525 | 0.144 |
| rpca | SNF | **HD_2** | 35 | 13 | 0 | 0.522 | 0.153 |
| rpca | SNF | HD_1 | 33 | 15 | 0 | 0.523 | 0.154 |
| umap | SNF | **HD_2** | 33 | 15 | 0 | 0.522 | 0.147 |
| feature_clustering | uMKL | HD_1 | 32 | 16 | 0 | 0.524 | 0.146 |
| feature_clustering | SNF | HD_1 | 32 | 16 | 0 | 0.523 | 0.153 |
| umap | SNF | HD_1 | 31 | 16 | 1 | 0.518 | 0.149 |
| entropy | SNF | **HD_2** | 30 | 17 | 1 | 0.515 | 0.15 |
| entropy | concatenation | HD_1 | 28 | 19 | 1 | 0.515 | 0.135 |
| rpca | MOFA+ | HD_1 | 25 | 23 | 0 | 0.507 | 0.139 |
| feature_clustering | concatenation | HD_1 | 25 | 21 | 2 | 0.513 | 0.133 |
| umap | concatenation | **HD_2** | 24 | 21 | 3 | 0.512 | 0.142 |
| feature_clustering | uMKL | **HD_2** | 24 | 17 | 7 | 0.512 | 0.127 |
| feature_clustering | concatenation | **HD_2** | 23 | 20 | 5 | 0.509 | 0.124 |
| umap | concatenation | HD_1 | 22 | 18 | 8 | 0.509 | 0.151 |
| entropy | concatenation | **HD_2** | 21 | 25 | 2 | 0.505 | 0.135 |
| rpca | uMKL | **HD_2** | 20 | 27 | 1 | 0.503 | 0.143 |
| feature_clustering | MOFA+ | HD_1 | 18 | 19 | 11 | 0.495 | 0.151 |
| entropy | SNF | HD_1 | 17 | 21 | 10 | 0.497 | 0.15 |
| entropy | MOFA+ | HD_1 | 16 | 23 | 9 | 0.499 | 0.14 |
| tsne | concatenation | HD_1 | 15 | 21 | 12 | 0.488 | 0.143 |
| tsne | SNF | HD_1 | 14 | 24 | 10 | 0.501 | 0.176 |
| umap | uMKL | HD_1 | 13 | 22 | 13 | 0.511 | 0.133 |
| laplacianEigenmaps | SNF | **HD_2** | 11 | 25 | 12 | 0.491 | 0.142 |

Figure S. F.8: Tabular visualization of win-tie-loss of the models that had the largest number of wins when using the AUCPR measure in the Wilcoxon signed-rank test comparing DR and data-fusion pipelines when heuristics $HD_1$ (column "ID definition" $= HD\_1$) or $HD_2$ (column "ID definition" $= HD\_2$) are used to set the dimension of the reduced space. The average values of AUCPR across all the cancer datasets are shown, together with their standard deviations. The top winners use heuristic $HD_2$.

| Heuristics | | | |
|---|---|---|---|
| HD_1 | 15 | | |
| HD_2 | 25 | | |
| **DR algorithm** | | **Data integration algorithm** | |
| feature_clustering | 9 | 18 | SNF |
| entropy | 8 | 12 | uMKL |
| rpca | 7 | 4 | MOFA+ |
| laplacianEigenmaps | 3 | 6 | concatenation |
| tsne | 5 | | |
| umap | 8 | | |

Figure S. F.9: Frequencies of heuristics, DR, and data-fusion algorithms appearing among the top-performing experiments, when using either heuristic $HD_1$ or heuristic $HD_2$ to set the dimension of the reduced space.

| Heuristic / block-analysis guided DR | | | |
|---|---|---|---|
| block-analysis | 32 | | |
| HD_2 | 10 | | |
| **DR algorithm** | | **Data integration algorithm** | |
| feature_clustering | 10 | 14 | SNF |
| par_rcur | 7 | 11 | uMKL |
| entropy | 11 | 9 | MOFA+ |
| rpca | 16 | 8 | concatenation |
| laplacianEigenmaps | 8 | | |
| umap | 8 | | |
| tsne | 4 | | |

Figure S. F.10: Frequencies of top-performing DR+data-fusion experiments obtained when comparing heuristic $HD_2$ and the block-analysis for guiding the reductions.

| DR pipeline | data_integration | use_id | ID definition | wins | ties | losses | auc | std(auc) |
|---|---|---|---|---|---|---|---|---|
| par_rcur+rpca | SNF | **TRUE** | **block-analysis** | 99 | 1 | 0 | 0.612 | 0.121 |
| par_rcur | MOFA+ | **TRUE** | **block-analysis** | 96 | 3 | 1 | 0.596 | 0.134 |
| feature_clustering | SNF | FALSE | HD_2 | 91 | 8 | 1 | 0.591 | 0.136 |
| umap | SNF | FALSE | HD_2 | 89 | 10 | 1 | 0.583 | 0.163 |
| rpca | SNF | FALSE | HD_2 | 88 | 10 | 2 | 0.589 | 0.126 |
| feature_clustering+rpca | SNF | **TRUE** | **block-analysis** | 88 | 10 | 2 | 0.589 | 0.131 |
| feature_clustering | SNF | **TRUE** | **block-analysis** | 87 | 11 | 2 | 0.588 | 0.131 |
| rpca | SNF | **TRUE** | **block-analysis** | 85 | 13 | 2 | 0.585 | 0.13 |
| par_rcur | SNF | **TRUE** | **block-analysis** | 85 | 13 | 2 | 0.584 | 0.12 |
| entropy | uMKL | FALSE | HD_2 | 84 | 13 | 3 | 0.577 | 0.144 |
| par_rcur+laplacianEigenmaps | SNF | **TRUE** | **block-analysis** | 84 | 12 | 4 | 0.583 | 0.119 |
| entropy | SNF | FALSE | HD_2 | 82 | 15 | 3 | 0.573 | 0.158 |
| entropy+umap | SNF | **TRUE** | **block-analysis** | 78 | 15 | 7 | 0.577 | 0.132 |
| feature_clustering | uMKL | **TRUE** | **block-analysis** | 77 | 17 | 6 | 0.576 | 0.145 |
| rpca | uMKL | FALSE | HD_2 | 75 | 18 | 7 | 0.572 | 0.151 |
| feature_clustering+rpca | uMKL | **TRUE** | **block-analysis** | 69 | 22 | 9 | 0.558 | 0.17 |
| par_rcur | uMKL | **TRUE** | **block-analysis** | 69 | 20 | 11 | 0.564 | 0.155 |
| entropy+rpca | MOFA+ | **TRUE** | **block-analysis** | 66 | 23 | 11 | 0.567 | 0.141 |
| entropy | SNF | **TRUE** | **block-analysis** | 66 | 22 | 12 | 0.569 | 0.137 |
| par_rcur+umap | SNF | **TRUE** | **block-analysis** | 66 | 22 | 12 | 0.569 | 0.136 |
| entropy+laplacianEigenmaps | SNF | **TRUE** | **block-analysis** | 66 | 22 | 12 | 0.569 | 0.133 |
| rpca | MOFA+ | **TRUE** | **block-analysis** | 65 | 23 | 12 | 0.567 | 0.133 |
| feature_clustering+umap | SNF | **TRUE** | **block-analysis** | 64 | 23 | 13 | 0.565 | 0.161 |
| feature_clustering+rpca | MOFA+ | **TRUE** | **block-analysis** | 63 | 23 | 14 | 0.566 | 0.141 |
| laplacianEigenmaps | SNF | **TRUE** | **block-analysis** | 59 | 26 | 15 | 0.567 | 0.128 |

Figure S. F.11: Tabular visualization of win-tie-loss tables (where AUC is used as the evaluation measure) obtained when pipelines using heuristic $HD_2$ for setting the dimensionality of the reduced space are compared to experiments using the block analysis for guiding the reductions. We show only the twenty five top-winners experiments, but all the results are reported in supplementary table S5). Note that the most of the top-most winner pipelines that lose less than twice are guided by block-analysis.

| DR pipeline | data_integratio | use_id | ID definition | wins | ties | losses | aucpr | std(aucpr) |
|---|---|---|---|---|---|---|---|---|
| par_rcur+rpca | SNF | TRUE | **block-analysis** | 89 | 11 | 0 | 0.531 | 0.146 |
| entropy | uMKL | FALSE | HD_2 | 87 | 13 | 0 | 0.532 | 0.148 |
| feature_clustering | SNF | FALSE | HD_2 | 84 | 16 | 0 | 0.525 | 0.144 |
| rpca | SNF | FALSE | HD_2 | 82 | 18 | 0 | 0.522 | 0.153 |
| feature_clustering+rpca | uMKL | TRUE | **block-analysis** | 82 | 18 | 0 | 0.517 | 0.157 |
| rpca | SNF | TRUE | **block-analysis** | 76 | 23 | 1 | 0.52 | 0.157 |
| umap | SNF | FALSE | HD_2 | 74 | 26 | 0 | 0.522 | 0.147 |
| feature_clustering | uMKL | TRUE | **block-analysis** | 67 | 32 | 1 | 0.52 | 0.143 |
| feature_clustering+rpca | SNF | TRUE | **block-analysis** | 66 | 33 | 1 | 0.517 | 0.148 |
| par_rcur | uMKL | TRUE | **block-analysis** | 65 | 31 | 4 | 0.516 | 0.134 |
| entropy | SNF | FALSE | HD_2 | 64 | 35 | 1 | 0.515 | 0.15 |
| feature_clustering | SNF | TRUE | **block-analysis** | 64 | 32 | 4 | 0.517 | 0.151 |
| umap | uMKL | TRUE | **block-analysis** | 63 | 36 | 1 | 0.521 | 0.155 |
| entropy | SNF | TRUE | **block-analysis** | 62 | 33 | 5 | 0.517 | 0.149 |
| par_rcur+rpca | uMKL | TRUE | **block-analysis** | 59 | 40 | 1 | 0.512 | 0.15 |
| entropy+rpca | uMKL | TRUE | **block-analysis** | 58 | 41 | 1 | 0.51 | 0.158 |
| par_rcur | SNF | TRUE | **block-analysis** | 57 | 37 | 6 | 0.515 | 0.145 |
| rpca | uMKL | TRUE | **block-analysis** | 56 | 43 | 1 | 0.51 | 0.157 |
| umap | concatenation | FALSE | HD_2 | 55 | 40 | 5 | 0.512 | 0.142 |
| entropy+umap | SNF | TRUE | **block-analysis** | 54 | 43 | 3 | 0.513 | 0.146 |
| entropy+rpca | concatenation | TRUE | **block-analysis** | 54 | 42 | 4 | 0.504 | 0.146 |
| feature_clustering | concatenation | FALSE | HD_2 | 54 | 40 | 6 | 0.509 | 0.124 |
| entropy | concatenation | FALSE | HD_2 | 53 | 44 | 3 | 0.505 | 0.135 |
| entropy | uMKL | TRUE | **block-analysis** | 53 | 40 | 7 | 0.512 | 0.141 |
| feature_clustering | uMKL | FALSE | HD_2 | 53 | 39 | 8 | 0.512 | 0.127 |

Figure S. F.12: Tabular visualization of win-tie-loss tables (where AUCPR is used as the evaluation measure) obtained when pipelines using heuristic $HD_2$ for setting the dimensionality of the reduced space are compared to experiments using the block analysis for guiding the reductions. We show only the twenty five top-winners experiments, but all the results are reported in supplementary table S6). Note that the most of the top-most winner pipelines that lose less than twice are guided by block-analysis.

| DR pipeline | data_integration | wins | ties | losses | auc | std(auc) |
|---|---|---|---|---|---|---|
| par_rcur+rpca | SNF | 73 | 3 | 0 | 0.612 | 0.121 |
| feature_clustering+rpca | SNF | 63 | 12 | 1 | 0.589 | 0.131 |
| rpca | SNF | 62 | 13 | 1 | 0.585 | 0.13 |
| feature_clustering | SNF | 60 | 15 | 1 | 0.588 | 0.131 |
| par_rcur | SNF | 60 | 15 | 1 | 0.584 | 0.12 |
| par_rcur | MOFA+ | 50 | 26 | 0 | 0.596 | 0.134 |
| par_rcur+laplacianEigenmaps | SNF | 47 | 28 | 1 | 0.583 | 0.119 |
| entropy+rpca | MOFA+ | 47 | 28 | 1 | 0.567 | 0.141 |
| feature_clustering | uMKL | 46 | 28 | 2 | 0.576 | 0.145 |
| entropy | SNF | 44 | 27 | 5 | 0.569 | 0.137 |
| feature_clustering+rpca | uMKL | 43 | 33 | 0 | 0.558 | 0.17 |
| rpca | MOFA+ | 43 | 32 | 1 | 0.567 | 0.133 |
| par_rcur | uMKL | 43 | 28 | 5 | 0.564 | 0.155 |
| entropy+rpca | uMKL | 40 | 34 | 2 | 0.551 | 0.176 |
| entropy+rpca | concatenation | 38 | 33 | 5 | 0.554 | 0.153 |
| entropy+umap | SNF | 37 | 38 | 1 | 0.577 | 0.132 |
| entropy+laplacianEigenmaps | SNF | 37 | 38 | 1 | 0.569 | 0.133 |
| laplacianEigenmaps | SNF | 37 | 32 | 7 | 0.567 | 0.128 |
| rpca | concatenation | 35 | 39 | 2 | 0.543 | 0.188 |
| feature_clustering+rpca | MOFA+ | 35 | 37 | 4 | 0.566 | 0.141 |
| entropy+rpca | SNF | 35 | 36 | 5 | 0.564 | 0.134 |
| feature_clustering+umap | concatenation | 35 | 34 | 7 | 0.556 | 0.174 |
| par_rcur+umap | SNF | 34 | 36 | 6 | 0.569 | 0.136 |
| feature_clustering+umap | SNF | 33 | 36 | 7 | 0.565 | 0.161 |
| umap | SNF | 33 | 36 | 7 | 0.558 | 0.158 |

Figure S. F.13: Extracts of win-tie-loss tables obtained when comparing each DR+data-fusion pipeline guided by block-analysis against each other. The top twenty-five winners are shown. Each pipeline reduced and then integrates all the available four omics. AUC is used for comparison.

| DR pipeline | data_integration | wins | ties | losses | aucpr | std(aucpr) |
|---|---|---|---|---|---|---|
| par_rcur+rpca | SNF | 71 | 5 | 0 | 0.531 | 0.146 |
| feature_clustering+rpca | uMKL | 65 | 11 | 0 | 0.517 | 0.157 |
| rpca | SNF | 60 | 15 | 1 | 0.52 | 0.157 |
| feature_clustering | uMKL | 53 | 22 | 1 | 0.52 | 0.143 |
| feature_clustering+rpca | SNF | 52 | 23 | 1 | 0.517 | 0.148 |
| par_rcur | uMKL | 51 | 24 | 1 | 0.516 | 0.134 |
| feature_clustering | SNF | 50 | 25 | 1 | 0.517 | 0.151 |
| umap | uMKL | 49 | 26 | 1 | 0.521 | 0.155 |
| entropy | SNF | 47 | 27 | 2 | 0.517 | 0.149 |
| par_rcur+rpca | uMKL | 45 | 30 | 1 | 0.512 | 0.15 |
| entropy+rpca | uMKL | 44 | 31 | 1 | 0.51 | 0.158 |
| par_rcur | SNF | 43 | 30 | 3 | 0.515 | 0.145 |
| rpca | uMKL | 42 | 33 | 1 | 0.51 | 0.157 |
| entropy+rpca | concatenation | 42 | 33 | 1 | 0.504 | 0.146 |
| entropy+umap | SNF | 40 | 34 | 2 | 0.513 | 0.146 |
| entropy | uMKL | 40 | 32 | 4 | 0.512 | 0.141 |
| entropy+rpca | SNF | 40 | 30 | 6 | 0.507 | 0.142 |
| feature_clustering+umap | concatenation | 39 | 35 | 2 | 0.52 | 0.134 |
| par_rcur | MOFA+ | 39 | 35 | 2 | 0.515 | 0.145 |
| rpca | concatenation | 39 | 35 | 2 | 0.502 | 0.144 |
| par_rcur+laplacianEigenmaps | SNF | 38 | 36 | 2 | 0.506 | 0.138 |
| feature_clustering+rpca | MOFA+ | 38 | 35 | 3 | 0.511 | 0.123 |
| laplacianEigenmaps | SNF | 36 | 34 | 6 | 0.505 | 0.146 |
| umap | SNF | 36 | 34 | 6 | 0.503 | 0.138 |
| feature_clustering+laplacianEigenmaps | SNF | 35 | 32 | 9 | 0.498 | 0.138 |

Figure S. F.14: Extracts of win-tie-loss tables obtained when comparing each DR+data-fusion pipeline guided by block-analysis against each other. The top twenty-five winners are shown. Each pipeline reduced and then integrates the four omics. AUCPR is used for comparison.

| data | DR pipeline | data_integration | wins | ties | losses | auc | std(auc) |
|---|---|---|---|---|---|---|---|
| miRNA_mRNA | rpca | SNF | 827 | 9 | 0 | 0.614 | 0.134 |
| miRNA_methy | par_rcur | SNF | 827 | 9 | 0 | 0.612 | 0.125 |
| miRNA_methy | feature_clustering | SNF | 827 | 9 | 0 | 0.608 | 0.142 |
| miRNA_methy | par_rcur+rpca | SNF | 826 | 10 | 0 | 0.612 | 0.119 |
| miRNA_mRNA_methy | par_rcur+rpca | SNF | 825 | 11 | 0 | 0.612 | 0.122 |
| miRNA_Proteins_methy | par_rcur+rpca | SNF | 821 | 15 | 0 | 0.609 | 0.131 |
| miRNA_mRNA_Proteins_methy | par_rcur+rpca | SNF | 820 | 16 | 0 | 0.612 | 0.121 |
| miRNA_Proteins_methy | feature_clustering+rpca | SNF | 818 | 18 | 0 | 0.606 | 0.131 |
| miRNA_methy | rpca | SNF | 815 | 21 | 0 | 0.605 | 0.129 |
| miRNA_methy | feature_clustering+rpca | SNF | 809 | 24 | 3 | 0.601 | 0.133 |
| miRNA_mRNA_methy | par_rcur | SNF | 808 | 24 | 4 | 0.6 | 0.125 |
| miRNA_Proteins_methy | rpca | SNF | 806 | 25 | 5 | 0.604 | 0.133 |
| miRNA_mRNA | entropy+rpca | SNF | 805 | 26 | 5 | 0.601 | 0.136 |
| miRNA_mRNA | par_rcur+rpca | SNF | 804 | 27 | 5 | 0.599 | 0.136 |
| miRNA_mRNA | feature_clustering+rpca | SNF | 803 | 27 | 6 | 0.599 | 0.143 |
| miRNA_mRNA | feature_clustering+rpca | MOFA+ | 801 | 30 | 5 | 0.595 | 0.16 |
| miRNA_mRNA | par_rcur+rpca | MOFA+ | 800 | 29 | 7 | 0.596 | 0.145 |
| miRNA_mRNA_methy | rpca | SNF | 800 | 28 | 8 | 0.597 | 0.127 |
| miRNA_methy | feature_clustering | uMKL | 799 | 30 | 7 | 0.597 | 0.152 |
| miRNA_Proteins_methy | par_rcur | SNF | 799 | 29 | 8 | 0.597 | 0.14 |
| miRNA_mRNA_Proteins_methy | par_rcur | MOFA+ | 797 | 30 | 9 | 0.596 | 0.134 |
| miRNA_Proteins_methy | feature_clustering | SNF | 797 | 30 | 9 | 0.595 | 0.14 |
| mRNA_Proteins_methy | feature_clustering+rpca | uMKL | 793 | 35 | 8 | 0.583 | 0.152 |
| miRNA_mRNA_Proteins | feature_clustering+rpca | MOFA+ | 790 | 37 | 9 | 0.594 | 0.156 |
| miRNA_Proteins | par_rcur+rpca | SNF | 789 | 38 | 9 | 0.594 | 0.137 |

Figure S. F.15: Extracts of win-tie-loss tables obtained when comparing each DR+data-fusion pipeline against each other when at least two omics data-views are integrated. AUC is used for comparison.

| data | DR pipeline | data_integration | wins | ties | losses | aucpr | std(aucpr) |
|---|---|---|---|---|---|---|---|
| miRNA_mRNA_methy | feature_clustering | uMKL | 819 | 17 | 0 | 0.537 | 0.139 |
| miRNA_methy | entropy | SNF | 816 | 20 | 0 | 0.533 | 0.141 |
| miRNA_methy | feature_clustering | SNF | 808 | 28 | 0 | 0.533 | 0.14 |
| miRNA_methy | par_rcur | SNF | 802 | 34 | 0 | 0.533 | 0.151 |
| miRNA_methy | par_rcur+rpca | SNF | 802 | 34 | 0 | 0.531 | 0.15 |
| miRNA_mRNA_Proteins_methy | par_rcur+rpca | SNF | 800 | 36 | 0 | 0.531 | 0.146 |
| miRNA_mRNA_methy | entropy | SNF | 793 | 43 | 0 | 0.529 | 0.148 |
| mRNA_Proteins_methy | feature_clustering+ | concatenation | 791 | 45 | 0 | 0.53 | 0.13 |
| miRNA_mRNA_methy | rpca | SNF | 791 | 45 | 0 | 0.528 | 0.152 |
| miRNA_mRNA_methy | par_rcur+rpca | SNF | 790 | 46 | 0 | 0.529 | 0.14 |
| miRNA_methy | feature_clustering | uMKL | 787 | 49 | 0 | 0.528 | 0.149 |
| mRNA_methy | umap | concatenation | 786 | 50 | 0 | 0.527 | 0.138 |
| miRNA_methy | feature_clustering+ | SNF | 780 | 56 | 0 | 0.527 | 0.147 |
| mRNA_methy | entropy | SNF | 777 | 58 | 1 | 0.527 | 0.132 |
| miRNA_Proteins_methy | par_rcur+rpca | SNF | 774 | 62 | 0 | 0.528 | 0.156 |
| miRNA_Proteins_methy | entropy | SNF | 767 | 69 | 0 | 0.527 | 0.157 |
| miRNA_methy | rpca | SNF | 763 | 73 | 0 | 0.526 | 0.151 |
| miRNA_mRNA_methy | par_rcur | uMKL | 763 | 72 | 1 | 0.529 | 0.135 |
| mRNA_methy | umap | SNF | 762 | 73 | 1 | 0.522 | 0.15 |
| miRNA_mRNA_methy | entropy | uMKL | 754 | 80 | 2 | 0.526 | 0.137 |
| miRNA_mRNA_methy | par_rcur | SNF | 754 | 80 | 2 | 0.526 | 0.135 |
| miRNA_Proteins_methy | par_rcur | SNF | 753 | 83 | 0 | 0.527 | 0.162 |
| miRNA_mRNA_methy | feature_clustering | SNF | 753 | 81 | 2 | 0.525 | 0.145 |
| miRNA_methy | entropy+rpca | SNF | 753 | 81 | 2 | 0.525 | 0.139 |
| miRNA_mRNA | par_rcur+rpca | MOFA+ | 748 | 86 | 2 | 0.524 | 0.135 |

Figure S. F.16: Extracts of win-tie-loss tables obtained when comparing each DR+data-fusion pipeline against each other when at least two omics data-views are integrated. AUCPR is used for comparison.

| data | DR pipeline | data_integration | wins | ties | losses | auc | std(auc) |
|---|---|---|---|---|---|---|---|
| 4 omics + pt | rpca | MOFA+ + PT data | 208 | 1 | 0 | 0.619 | 0.154 |
| 4 omics | par_rcur+rpca | SNF | 201 | 7 | 1 | 0.612 | 0.121 |
| 4 omics + pt | par_rcur+rpca | SNF | 201 | 7 | 1 | 0.612 | 0.123 |
| 4 omics + pt | par_rcur+rpca | SNF + PT data | 201 | 7 | 1 | 0.611 | 0.118 |
| 4 omics + pt | rpca | SNF + PT data | 201 | 7 | 1 | 0.61 | 0.118 |
| 4 omics + pt | entropy+rpca | MOFA+ + PT data | 201 | 7 | 1 | 0.608 | 0.132 |
| 4 omics + pt | par_rcur | MOFA+ | 199 | 9 | 1 | 0.605 | 0.131 |
| 4 omics + pt | par_rcur | MOFA+ + PT data | 199 | 9 | 1 | 0.601 | 0.176 |
| 4 omics + pt | par_rcur | SNF + PT data | 191 | 12 | 6 | 0.6 | 0.129 |
| 4 omics + pt | par_rcur+laplacianEigenmaps | SNF + PT data | 190 | 13 | 6 | 0.599 | 0.141 |
| 4 omics | par_rcur | MOFA+ | 189 | 12 | 8 | 0.596 | 0.134 |
| 4 omics + pt | feature_clustering+laplacianEigenmaps | SNF + PT data | 184 | 17 | 8 | 0.594 | 0.151 |
| 4 omics + pt | feature_clustering+rpca | MOFA+ | 180 | 21 | 8 | 0.591 | 0.141 |
| 4 omics + pt | par_rcur | SNF | 179 | 22 | 8 | 0.593 | 0.121 |
| 4 omics + pt | feature_clustering+rpca | SNF | 179 | 22 | 8 | 0.592 | 0.125 |
| 4 omics + pt | par_rcur+rpca | MOFA+ + PT data | 179 | 22 | 8 | 0.58 | 0.181 |
| 4 omics + pt | feature_clustering+rpca | MOFA+ + PT data | 178 | 23 | 8 | 0.588 | 0.154 |
| 4 omics + pt | entropy+rpca | MOFA+ | 178 | 23 | 8 | 0.584 | 0.17 |
| 4 omics + pt | feature_clustering | SNF + PT data | 176 | 24 | 9 | 0.592 | 0.136 |
| 4 omics + pt | par_rcur+laplacianEigenmaps | SNF | 174 | 25 | 10 | 0.589 | 0.117 |
| 4 omics + pt | feature_clustering | SNF | 174 | 24 | 11 | 0.589 | 0.138 |
| 4 omics | feature_clustering+rpca | SNF | 174 | 24 | 11 | 0.589 | 0.131 |
| 4 omics + pt | feature_clustering+rpca | SNF + PT data | 173 | 25 | 11 | 0.587 | 0.129 |
| 4 omics + pt | entropy | SNF + PT data | 172 | 26 | 11 | 0.587 | 0.139 |
| 4 omics | feature_clustering | SNF | 171 | 27 | 11 | 0.588 | 0.131 |

Figure S. F.17: Extracts of win-tie-loss tables obtained when comparing each DR+data-fusion pipeline against each other when all the available omics and patients descriptors are analyzed (the combination of data views that are input to the pipeline also specifies the pipeline). AUC is used for comparison. “MOFA+ + PT data”, “SNF + PT data”, and “uMKL + PT data” refer to the data-fusion algorithms integrating the demographic view. “MOFA+”, “SNF”, and “uMKL” refer to the traditional application of the data-fusion algorithms for integrating multi-omics, followed by concatenation with the demographics views.

| data | DR pipeline | data_integration | wins | ties | losses | aucpr | std(aucpr) |
|---|---|---|---|---|---|---|---|
| 4 omics + pt | rpca | MOFA+ + PT data | 207 | 2 | 0 | 0.549 | 0.148 |
| 4 omics + pt | entropy+rpca | MOFA+ + PT data | 202 | 7 | 0 | 0.54 | 0.129 |
| 4 omics + pt | par_rcur+rpca | SNF | 200 | 8 | 1 | 0.534 | 0.15 |
| 4 omics + pt | par_rcur | MOFA+ + PT data | 199 | 9 | 1 | 0.535 | 0.141 |
| 4 omics | par_rcur+rpca | SNF | 196 | 12 | 1 | 0.531 | 0.146 |
| 4 omics + pt | par_rcur+rpca | MOFA+ + PT data | 195 | 13 | 1 | 0.534 | 0.143 |
| 4 omics + pt | feature_clustering+rpca | MOFA+ | 191 | 17 | 1 | 0.53 | 0.129 |
| 4 omics + pt | entropy+rpca | MOFA+ | 177 | 30 | 2 | 0.528 | 0.121 |
| 4 omics + pt | feature_clustering+rpca | MOFA+ + PT data | 177 | 30 | 2 | 0.527 | 0.13 |
| 4 omics + pt | feature_clustering+umap | concatenation | 170 | 36 | 3 | 0.526 | 0.133 |
| 4 omics + pt | par_rcur | MOFA+ | 165 | 40 | 4 | 0.524 | 0.15 |
| 4 omics | umap | uMKL | 165 | 40 | 4 | 0.521 | 0.155 |
| 4 omics + pt | par_rcur | uMKL | 162 | 42 | 5 | 0.525 | 0.14 |
| 4 omics + pt | par_rcur+rpca | SNF + PT data | 161 | 42 | 6 | 0.522 | 0.139 |
| 4 omics | rpca | SNF | 159 | 46 | 4 | 0.52 | 0.157 |
| 4 omics + pt | entropy | SNF | 157 | 46 | 6 | 0.519 | 0.15 |
| 4 omics + pt | feature_clustering+rpca | uMKL | 155 | 48 | 6 | 0.521 | 0.158 |
| 4 omics + pt | feature_clustering | SNF | 155 | 48 | 6 | 0.52 | 0.153 |
| 4 omics + pt | rpca | SNF + PT data | 155 | 47 | 7 | 0.52 | 0.146 |
| 4 omics | feature_clustering | uMKL | 154 | 48 | 7 | 0.52 | 0.143 |
| 4 omics + pt | feature_clustering | uMKL | 154 | 48 | 7 | 0.52 | 0.142 |
| 4 omics + pt | par_rcur | SNF | 154 | 48 | 7 | 0.52 | 0.144 |
| 4 omics | entropy | SNF | 154 | 48 | 7 | 0.517 | 0.149 |
| 4 omics + pt | umap | uMKL | 153 | 49 | 7 | 0.518 | 0.155 |
| 4 omics | feature_clustering+rpca | SNF | 153 | 49 | 7 | 0.517 | 0.148 |

Figure S. F.18: Extracts of win-tie-loss tables obtained when comparing each DR+data-fusion pipeline against each other when all the available omics and patients descriptors are analyzed (the combination of data views that is input to the pipeline also specifies the pipeline). AUCPR is used for comparison. "MOFA+ + PT data", "SNF + PT data", and "uMKL + PT data" refer to the data-fusion algorithms integrating the demographic view. "MOFA+", "SNF", and "uMKL" refer to the traditional application of the data-fusion algorithms for integrating multi-omics, followed by concatenation with the demographics views.
